# Generative AI Framework SynGlue for the Rational Design of Clinically relevant Protein Degraders

**DOI:** 10.1101/2025.08.28.672835

**Authors:** Saveena Solanki, Sanjay Kumar Mohanty, Shiva Satija, Sonam Chauhan, N.V.M. Rao Bandaru, Sandeep Dukare, Nirbhay Kumar Tiwari, R Naveen Kumar, A B Aravind, Subhendu Mukherjee, Dinesh Chikkanna, Wesley Roy Balasubramanian, Srinivasa Raju Sammeta, Vishakha Gautam, Sakshi Arora, Suvendu Kumar, Subhadeep Duari, Arushi Sharma, Raidhani Shome, Debarka Sengupta, Chandrasekhar Abbineni, Susanta Samajdar, Gaurav Ahuja

**Author notes:** Shared First Authors (Saveena Solanki, Sanjay Kumar Mohanty). **Correspondence:** Gaurav Ahuja; Susanta Samajdar.

## Abstract

The rational design of protein degraders, such as proteolysis-targeting chimeras (PROTACs), requires the simultaneous optimization of multiple molecular properties, a complex challenge that limits efficient discovery. Here, we introduce SynGlue, a generative artificial intelligence (AI) framework that addresses this challenge through two core modules: data-driven, leveraging large-scale protein-ligand intelligence, and structure-guided, for physics-aware molecular design. SynGlue harness MagnetDB, a curated database of 6.37 million experimental protein-ligand interactions, and couples it with deep learning models that quantitatively predict degradation potency (DC_50_), maximal degradation (D_max_), and guide ternary-complex-compatible linker design. Benchmarked against 6,935 compounds, SynGlue demonstrates superior performance in relevant pharmacology prediction. To validate SynGlue, we engineered degraders for BRD4 and GSPT1. Our data-driven design for BRD4 yielded compounds with novel warhead scaffolds (<50% warhead similarity with known PROTACs), which proved to be potent degraders *in vitro* (DC_50_ = 0.19 nM) and efficacious *in vivo* in mouse models. Independently, our structure-guided *de novo* design for GSPT1 produced ultrapotent degraders (DC_50_ ≈ 0.0011 μM) that are also effective both *in vitro* and *in vivo*, uncovering a new oncogenic dependency. By unifying data-driven and physics-aware design, SynGlue establishes a generalizable AI framework for the rapid development of clinically relevant protein degraders, with principled extension to other multi-target modalities.

## INTRODUCTION

The drug discovery paradigm is rapidly evolving beyond the traditional “one target, one drug” framework toward strategies that better reflect the biological complexity of human disease^1,2^. Polypharmacological molecules, which modulate multiple targets or pathways simultaneously, have emerged as a powerful approach to achieve enhanced efficacy and overcome resistance, particularly in multifactorial diseases such as cancer^3,4,5–7^. These molecules can act through diverse mechanisms, including synergistic inhibition ^8^, induced protein-protein proximity ^9^, and Targeted Protein Degradation (TPD) ^10^, most notably via Proteolysis Targeting Chimeras (PROTACs) ^11,12^. By recruiting an E3 ubiquitin ligase to a protein of interest, PROTACs have substantially expanded the druggable proteome ^13^. Despite their potential, the rational design of potent PROTACs remains a significant challenge ^14,15^. Productive degradation requires the coordinated optimization of three interdependent components, the target warhead, the E3 ligase ligand, and the linker, which together govern ternary complex formation and degradation efficiency (DC_50_, D_max_)^16,17^. Small structural changes can lead to large, non-intuitive shifts in activity, rendering empirical optimization inefficient. While structure-based modeling is computationally intensive and conformation-sensitive, many AI-driven methods rely on whole-molecule representations that obscure component-specific contributions, limiting interpretability and generalizability ^18–20^.

A key conceptual limitation is this reliance on whole-molecule similarity as a proxy for function, which fails to capture how individual binding fragments encode specificity in modular architectures. Fragment-centric inference is needed to deconvolve multi-target activity while preserving mechanistic interpretability. Existing platforms offer either broad modeling capabilities (e.g., MOE, ICM) or specialized focus (e.g., PRosettaC ^21^ for ternary complexes, DiffPROTACs ^22^ for linkers), but face trade-offs in accuracy, computational cost, accessibility, or generalizability beyond PROTACs to other multi-target modalities ^23^. Furthermore, the optimal design strategy is context-dependent. For data-rich targets like BRD4, data-driven fragment mapping can efficiently prioritize tractable solutions. For ligand-poor targets like the translation termination factor GSPT1, an essential protein with sparse chemical precedent, structure-guided *de novo* discovery is necessary.

To address these challenges, we developed SynGlue, an integrated AI platform that unifies large-scale fragment-centric interaction intelligence with structure-guided molecular design. SynGlue functions as a decision framework that learns how individual molecular components encode target specificity and functional outcomes, systematically narrowing chemical space into actionable candidates. It integrates a comprehensive ligand-target knowledge base (MagnetDB) with scalable fragment indexing and predictive modeling. Supporting both data-driven and structure-guided workflows, SynGlue enables degrader discovery across heterogeneous contexts. Using BRD4 and GSPT1 as orthogonal case studies, we demonstrate its ability to generate selective, novel degraders from data-rich to ligand-poor regimes, with validation from computation to *in vitro* and *in vivo* efficacy.

## RESULTS

### SynGlue: An Integrated Platform for Multi-Target Mapping and PROTAC Design

SynGlue is an integrated computational platform that unifies large-scale data-driven interaction mapping with structure-guided molecular design for the systematic discovery and generation of multi-target and bifunctional therapeutics. Its framework consists of two complementary, interoperable modules: a data-driven module and a structure-guided module, that together integrate curated interaction databases, fragment-based indexing, quantitative scoring, and *de novo* design within a single workflow **(Figure 1a)**. This architecture enables SynGlue to perform accurate target inference, interpret polypharmacology, predict quantitative efficacy, and generate molecules, even without prior structural knowledge. At the core of the data-driven workflow is MagnetDB, a rigorously curated repository of 6.37 million high-confidence, experimentally validated protein-ligand interactions, distilled from over 37.5 million raw entries and involving 1.94 million unique compounds and 20,129 protein targets **(Supplementary Figure 1a; Supplementary Table 1)**. To enable scalable substructure searching across this space, all compounds are decomposed into terminal fragments using the RECAP algorithm, yielding ~2.4 million unique fragments **(Figure 1b; Supplementary Figure 1b-c)**. These are indexed using a TRIE (prefix tree) data structure, which preserves complete chemical fidelity and enables exact fragment reconstruction **(Supplementary Figure 1d)**. Benchmarking confirms that TRIE-based retrieval operates with near-linear O(n) time complexity, outperforming brute-force O(n^2^) matching by orders of magnitude **(Figure 1c; Supplementary Figure 1d-e)**. Search time scales modestly with fragment length and remains stable across diverse queries, confirming robustness **(Supplementary Figure 1f-i)**.

**Figure 1:**
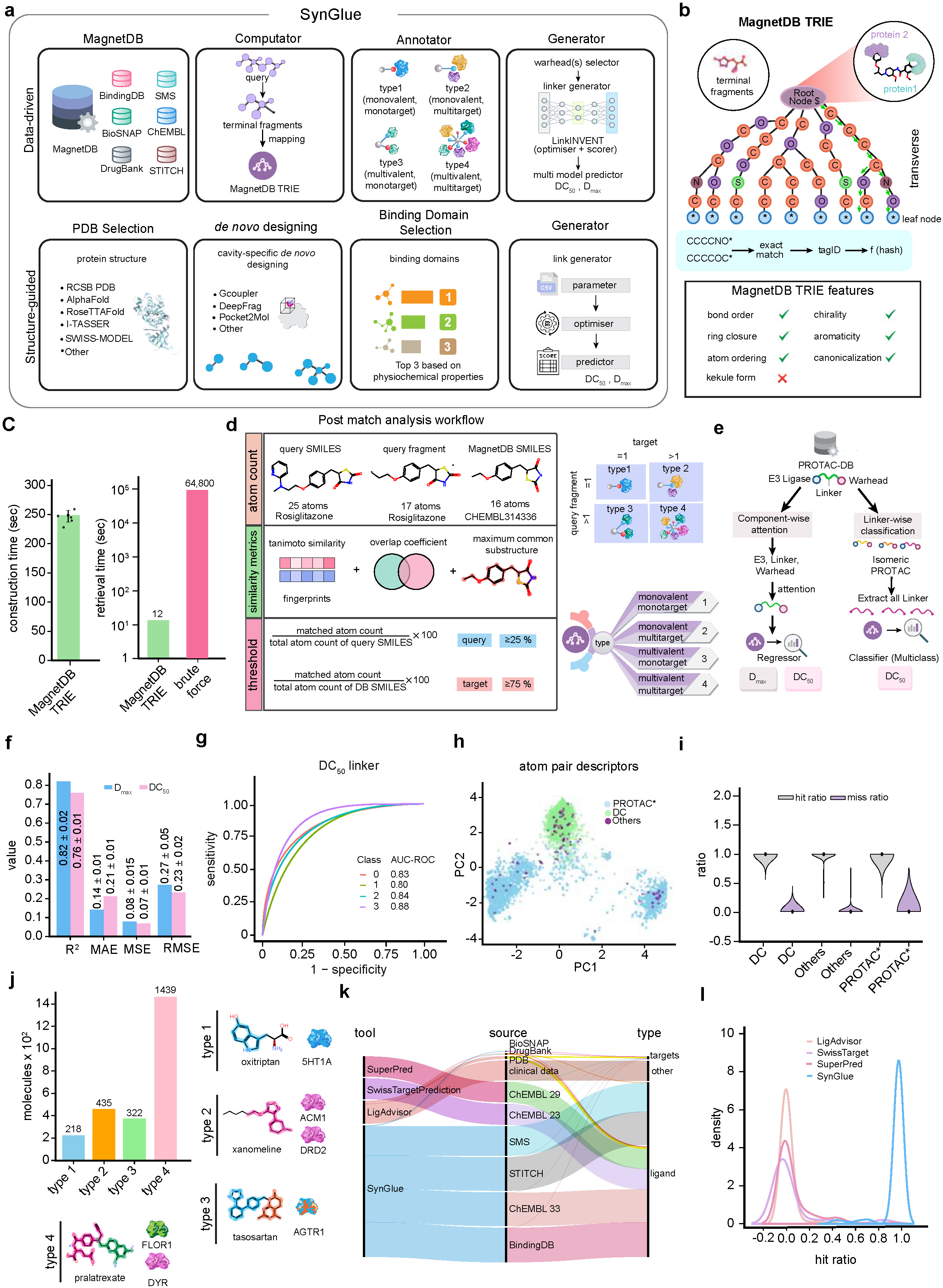
SynGlue’s data-driven architecture and MagnetDB performance. **(a)** Schematic detailing SynGlue’s dual workflows. The data-driven path (top) utilizes the MagnetDB interaction database, the TRIE-based Computator for fragment mapping, and the Annotator for classification, feeding into the Generator. The structure-guided path (bottom) starts with a target PDB structure, proceeds through de novo warhead design (using preferred tools) and pharmacological filtering, before inputting prioritized warheads and selected E3 ligases into the SynGlue Generator for de novo synthesis of multi-targeting molecules (MTMs) like PROTACs. **(b)** Schematic of the MagnetDB TRIE data structure used for efficient fragment matching. Terminal fragments are indexed as canonical SMILES into a trie, enabling rapid exact-match traversal from root to leaf nodes. Supported chemical features include bond order, chirality, aromaticity, atom ordering, and canonicalization. **(c)** Performance benchmarking comparing database construction and retrieval time between MagnetDB TRIE and brute-force approaches, demonstrating substantial acceleration in both indexing and query retrieval. **(d)** Post-match analysis workflow illustrating fragment-level comparison between query molecules and MagnetDB entries. Multiple similarity metrics, including fingerprint similarity, overlap coefficient, and maximum common substructure, are combined with atom-count-based thresholds to assign interaction types (type 1-4). **(e)** Machine-learning workflow for PROTAC analysis integrating component-wise attention over E3 ligase, linker, and warhead features. The pipeline supports both regression (DC_50_, D_max_) and classification (linker class, isomeric PROTACs). **(f)** Predictive performance of SynGlue regression models for DC_50_ and D_max_ shown using R^2^, MAE, MSE, and RMSE metrics across cross-validation runs. **(g)** Receiver operating characteristic (ROC) curves for multi-class DC_50_ linker classification, with area-under-the-curve (AU-ROC) values reported for each interaction type. **(h)** Principal component analysis (PCA) of atom-pair descriptors comparing PROTAC-positive compounds, DC-active molecules, and other chemical space, highlighting distinct clustering of functional degraders. **(i)** Violin plots showing hit-to-miss ratios across DC-active compounds, PROTAC-positive sets, and background molecules, illustrating improved enrichment for SynGlue-identified hits. **(j)** Distribution of molecules across four interaction types (types 1-4), with representative examples shown for each category, spanning monovalent/multivalent and mono-/multi-target interactions. **(k)** Sankey diagram comparing target recovery across tools (SynGlue, SuperPred, SwissTargetPrediction, LigAdvisor) and source databases, highlighting broader ligand–target coverage achieved by SynGlue. **(l)** Density distribution of hit ratios across target-prediction tools, demonstrating superior enrichment and reduced false positives for SynGlue compared to baseline methods.

SynGlue applies a dual-stage fragment matching strategy to map query molecules onto known interaction space **(Figure 1d)**, combining fast TRIE-based prefix searches with similarity refinement. This enables high-resolution fragment-target mapping and classifies compounds into four polypharmacological types (mono-/multivalent, mono-/multitarget), supporting interpretation of complex modalities like PROTACs **(Figure 1d)**. To enable quantitative efficacy prediction, SynGlue integrates a linker-centric machine-learning layer with transformer-based regression models for DC_50_ and D_max_ prediction **(Figure 1e-g)**. By decomposing PROTACs into warhead, linker, and E3 ligase components, the models directly capture linker-dependent effects. A linker-centric multiclass framework robustly stratifies PROTAC behavior **(Supplementary Figure 4a)**, showing consistent performance across representations **(Supplementary Figure 4b-k)**. Ten-fold cross-validation confirms generalizable classification with a median test AUC of ~0.80 **(Supplementary Figure 4l-n)**. In parallel, a transformer-based attention model predicts DC_50_ and D_max_ by learning component-wise contributions, achieving strong quantitative performance **(Figure 1f)**. When DC_50_ prediction is analyzed in a linker-aware setting, the model achieves high AUC-ROC values across potency classes **(Figure 1g)**. Together, these establish linker chemistry as a dominant, learnable determinant of degradation efficiency, forming the quantitative core of SynGlue’s Generator module **(Supplementary Figure 5b-h)**, which supports an integrated PROTAC design and prioritisation workflow **(Supplementary Figure 3a-c)**.

The accuracy of SynGlue’s data-driven and predictive components was validated using a curated benchmark of 6,935 compounds, including DrugCentral drugs, known PROTACs, and other multitarget entities **(Supplementary Figure 2a-c)**. Chemical space analysis reveals pronounced structural heterogeneity **(Figure 1h; Supplementary Figure 2h,i)**. Target recovery analysis demonstrates consistently high accuracy across compound classes **(Figure 1i; Supplementary Figure 2d,e)**. SynGlue successfully classified 4,621 compounds (66.7%) into defined interaction types, including complex PROTACs **(Figure 1j, Supplementary Table 3)**. Comparative benchmarking against LigAdvisor, SwissTargetPrediction, and SuperPred highlights SynGlue’s superior target recovery, particularly for multitarget and linker-containing molecules **(Figures 1k, l; Supplementary Figure 2f-l, Supplementary Table 4)**. Collectively, these results establish SynGlue as a complete, end-to-end platform that unifies fragment-based interaction mapping, polypharmacology classification, linker-aware machine learning, quantitative efficacy prediction, and structure-guided generative design.

### SynGlue Enables Design of PROTACs and Multi-Target Molecules

To evaluate SynGlue’s generative design capabilities, we applied it to the *de novo* creation of novel therapeutics. First, as an *in silico* proof of concept, we targeted the androgen receptor (AR) using both data-driven and structure-guided modalities. SynGlue generated AR-targeting PROTAC candidates by integrating warhead selection, linker optimization, and E3 ligase recruitment **(Figure 2a, b)**. The data-driven approach yielded candidates resembling more of the known AR degraders, while the structure-guided workflow, in addition to the known AR degraders, also possessed more diverse chemical space **(Supplementary Figure 5a-b)**. Additional characterization of atom count distributions and linker properties provides a comprehensive view of component diversity **(Supplementary Figure 3d-f)**. Using the data-driven approach, SynGlue identified potential AR-binding warheads from MagnetDB, selecting a validated candidate (CHEMBL146794) **(Figure 2b; Supplementary Figure 5a; Supplementary Table 5)**. The Generator module coupled this warhead to a known XIAP E3 ligase ligand to produce PROTAC candidates. Concurrently, we employed a structure-guided design workflow using the human AR crystal structure (PDB ID: 1E3G) with the Gcoupler tool ^24^ to design 404 *de novo* cavity-specific ligands, demonstrating SynGlue’s dual-workflow capability **(Figure 2i)**. The Gcoupler Authenticator module evaluated candidates based on molecular interaction profiles, and the selected warhead was used by SynGlue’s Generator with optimized linker parameters and the same XIAP E3 ligase. Comparative analysis showed that the data-driven approach produced analogs with high similarity to validated AR PROTACs, while the structure-guided approach generated more chemically diverse candidates, highlighting SynGlue’s complementary modalities for expanding ligand novelty. Beyond PROTACs, SynGlue supports the design of non-degrading multitarget molecules (MTMs). For an EGFR/CDK1 dual-target case, SynGlue identified validated warheads (CHEMBL939 and CHEMBL4442620) and designed candidate MTMs predicted to retain binding to both targets **(Supplementary Figure 6a)**. A similar successful outcome was achieved for the PRKX/DRD1 pair using identified warheads (CHEMBL2152768 and CHEMBL1201356) **(Supplementary Figure 6b; Supplementary Table 6)**. These examples underscore SynGlue’s versatility, as highlighted by linker property and chemical taxonomy analyses **(Supplementary Figure 6c-e)**. Finally, in a cross-platform comparison, SynGlue outperformed existing PROTAC prediction tool PDP by accurately ranking compound potency and quantitatively predicting both DC_50_ and D_max_ across VCaP and LNCaP cell lines, uniquely integrating linker-aware modeling with *de novo* design **(Supplementary Figure 7a-c)**.

**Figure 2:**
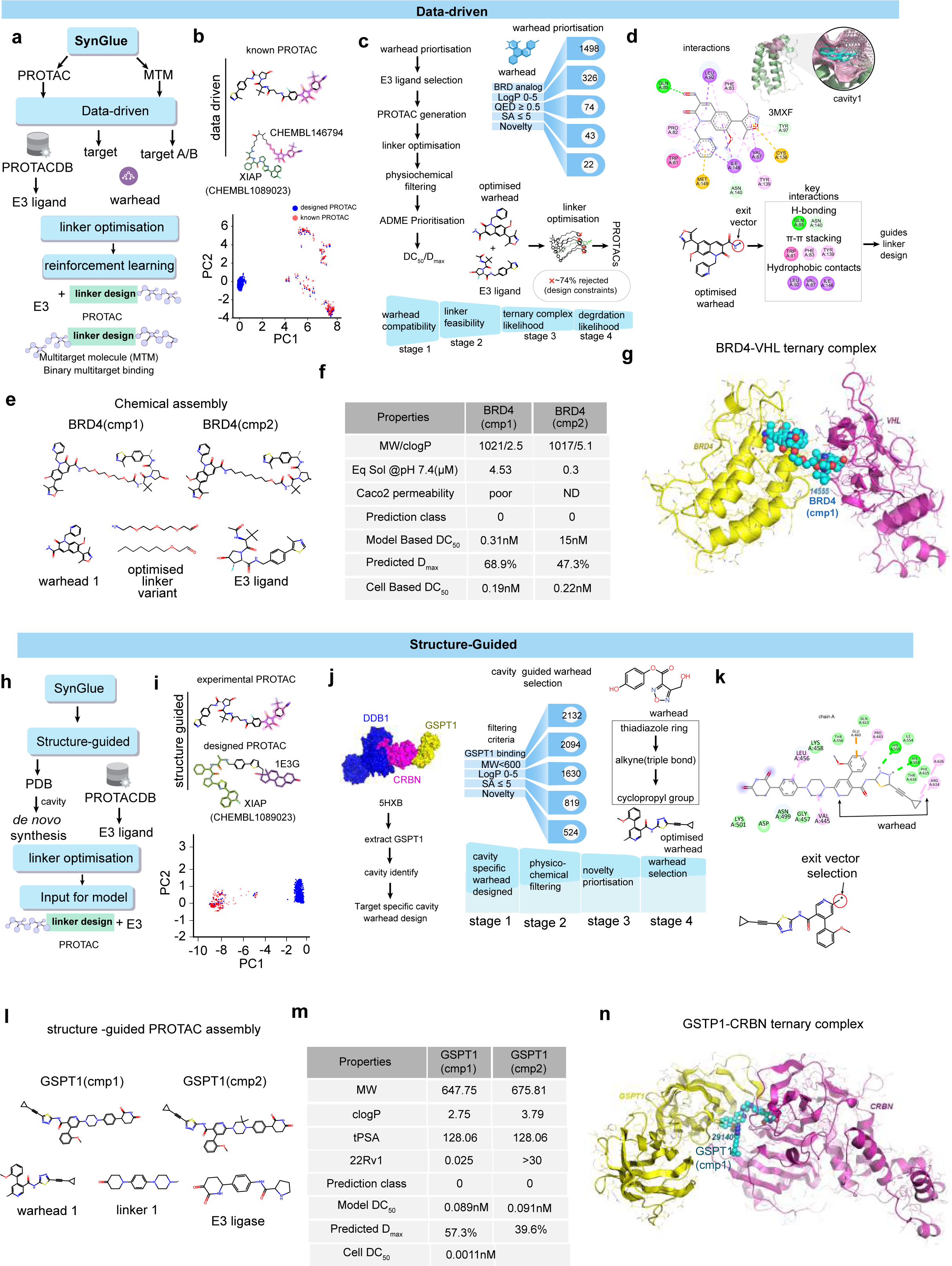
SynGlue enables data-driven and structure-guided design and prioritization of PROTACs. **(a)** Schematic overview of the data-driven SynGlue workflow for PROTAC and multitarget molecule (MTM) design. Starting from curated interaction data in PROTACDB and MagnetDB, SynGlue integrates warhead and E3 ligase selection, linker optimization, and reinforcement-learning-based linker design to generate candidate PROTACs and MTMs. **(b)** Chemical space projection comparing known Androgen Receptor (AR) PROTACs and SynGlue-designed PROTACs, illustrating that designed molecules occupy regions proximal to validated compounds while extending into adjacent chemical space. Representative examples include a known XIAP-based PROTAC and a data-driven designed analog derived from CHEMBL interaction data. **(c)** Data-driven warhead prioritization and PROTAC generation funnel. Starting from 1,498 BRD4-associated warheads identified from MagnetDB, successive filtering by physicochemical properties (logP, QED, SA) and novelty reduces the pool to 22 optimized warheads. These are combined with E3 ligands, followed by linker optimization and ADME-based prioritization to yield final PROTAC candidates, with ~74% of designs rejected by constraint-based filtering. **(d)** Representative interaction-level analysis of a prioritized BRD4 warhead docked into the BRD4 bromodomain (PDB: 3MXF). Key hydrogen bonding, π–π stacking, and hydrophobic interactions are highlighted, and the exit-vector orientation guiding linker attachment is shown, informing rational linker design for ternary complex formation. **(e)** Chemical assembly of data-driven BRD4 PROTACs, showing optimized warhead, linker variants, and E3 ligase components for two isomeric candidates, BRD4 (cmp1) and BRD4 (cmp2). **(f)** Comparative predicted and experimental properties of BRD4 (cmp1) and BRD4 (cmp2), including molecular weight, lipophilicity, solubility, permeability, predicted degradation metrics (DC_50_ and D_max_), and cell-based degradation potency, illustrating SynGlue’s ability to resolve activity differences between closely related isomers. **(g)** Modeled BRD4-PROTAC-VHL ternary complex, highlighting productive spatial arrangement and target-E3 proximity for BRD4 (cmp1), supporting its predicted degradation capability. **(h)** Overview of the structure-guided SynGlue pipeline, in which experimentally resolved protein structures are used for cavity-specific de novo warhead design, followed by linker optimization and PROTAC assembly. **(i)** Comparison of experimentally validated PROTACs and structure-guided designed PROTACs projected into chemical space, demonstrating that structure-guided designs explore distinct regions while maintaining relevance to known degraders. **(j)** Cavity-guided warhead selection funnel for GSPT1 using the CRBN-GSPT1 complex (PDB: 5HXB). De novo warheads are generated based on cavity geometry and filtered sequentially by binding compatibility, physicochemical properties, and novelty to yield optimized, cavity-specific warheads. **(k)** Structure-guided exit-vector determination for a selected GSPT1 warhead, showing residue-level interactions and linker attachment points derived directly from cavity orientation. **(l)** Chemical assembly of structure-guided GSPT1 PROTACs, illustrating warhead, linker, and E3 ligase components for GSPT1 (cmp1) and GSPT1 (cmp2). **(m)** Predicted and experimental physicochemical and degradation properties of GSPT1 PROTACs, including molecular weight, clogP, TPSA, predicted DCLL and D_max_, and cell-based degradation potency. **(n)** Modeled GSPT1-PROTAC-CRBN ternary complex, confirming spatial compatibility and stable interface formation that supports efficient target degradation.

To benchmark SynGlue across complementary design challenges, we selected BRD4 and GSPT1 as representative targets for PROTAC design using data-driven and structure-guided modalities, respectively **(Figure 2)**. BRD4, a well-characterized epigenetic target with extensive ligand data, tests SynGlue’s ability to achieve selectivity and novelty. In contrast, GSPT1, a translation termination factor with sparse direct ligands, provides a stringent test for *de novo*, cavity-specific warhead discovery. We initiated BRD4 PROTAC design by interrogating MagnetDB for BRD4-associated fragments, yielding 1,498 candidate warheads satisfying a stringent interaction-coverage threshold (>75%). To ensure paralog selectivity, candidates were cross-referenced against fragment profiles for BRD2, BRD3, and BRD9, removing 14 promiscuous fragments **(Supplementary Figure 9a)**. Sequential physicochemical filtering refined this pool: logP (0-5) reduced the set to 326, QED ≥ 0.5 narrowed it to 74, and synthetic accessibility (SA ≤5) yielded 43 tractable warheads **(Figure 2c; Supplementary Figure 9a-d, 5a)**. Novelty assessment against 42 known BRD-targeting warheads from PROTAC-DB 3.0, using ECFP4 Tanimoto similarity, eliminated compounds with TS≥0.5, resulting in 22 structurally novel warheads **(Figures 2c, d; Supplementary Figure 9a)**. Hierarchical clustering and RDKit analysis confirmed these span multiple chemotypes **(Figure 2e; Supplementary Figure 9b)**. Sequential physicochemical filtering reduced the initial compound set to a small, high-quality subset, reflecting stringent drug-likeness criteria **(Supplementary Figure 10a).** Docking into the BRD4 bromodomain (PDB: 3MXF) validated biological relevance, confirming preservation of key interactions (hydrogen bonding with Asn140, π–π stacking with Tyr97) and viable exit vectors **(Figure 2f, Supplementary Figure 8a-l)**. Selected warheads were coupled to a VHL E3 ligase ligand, with linker optimization guided by a multiclass feasibility model and attention-based regression models predicting DC_50_ and D_max_ **(Figure 2g; Supplementary Figure 9h, e-n),** designing diverse BRD4-VHL PROTACs using SynGlue **(Supplementary Table 7)**. SynGlue discriminated between closely related isomers, prioritizing two candidates, BRD4 (cmp1) and BRD4 (cmp2), based on balanced linker properties and predicted ternary stability **(Figure 2h)**, in silico characterization for prioritized BRD4-VHL PROTAC candidates **(Supplementary Table 8).** Similarity analysis against all reported BRD4 PROTACs confirmed low maximum Tanimoto similarity (0.493 and 0.465, respectively), with distributions centered between 0.30-0.45 and no scaffold-level overlap **(Figure 2i; Supplementary Figure 10b-e)**.

For GSPT1, we applied SynGlue’s structure-guided workflow using the CRBN-GSPT1 complex (PDB: 5HXB) as input **(Figure 2h-n)**. The pipeline first isolated the GSPT1-specific interaction surface from the multimeric assembly, eliminating E3-ligase-dominated features to anchor design exclusively to GSPT1 geometry **(Supplementary Figure 8e-i)**. Explicit cavity detection on this interface identified an addressable pocket proximal to the CRBN interaction region, defining the geometric and physicochemical constraints for *de novo* warhead generation **(Figure 2h; Supplementary Figure 8f)**. Candidate warheads were funneled through structure-aware prioritization: first filtered for cavity fit and steric compatibility, then evaluated using physicochemical criteria (molecular weight, logP, TPSA) conditional on cavity engagement **(Figure 2j; Supplementary Figure 9a, b)**. Novelty prioritization eliminated redundant chemotypes, yielding a compact set of GSPT1-specific warheads with well-defined exit vectors derived from cavity orientation **(Figure 2j, Supplementary Figure 9c)**. These warheads were assembled into full degraders using linker optimization constrained by the spatial relationship between GSPT1 and CRBN. Linkers were prioritized using a weighted geometric mean emphasizing flexibility, ring balance, and effective length **(Figure 2j-l, Supplementary Figure 10e-i; 11a-d, Supplementary Table 10)**. Degradation-prediction models revealed structure-guided GSPT1 PROTACs achieve sub-nanomolar predicted DC_50_ (≈0.089-0.091 nM) with high predicted degradation efficiency (D_max_ up to 57.3%), despite no prior warhead templates **(Figure 2m)**. Ternary complex modeling confirmed stable, coherent GSPT1-PROTAC-CRBN assemblies with favorable interface properties **(Figure 2n)**; this was followed by *in silico* characterization of the final prioritized GSPT1-CRBN PROTAC candidates, which supported their potential efficacy and selectivity **(Supplementary Figure 8g; Supplementary Table 11)**. Together, these orthogonal benchmarks demonstrate SynGlue’s generalizable framework. BRD4 validates precise, novel prioritization in data-rich contexts, while GSPT1 establishes *de novo* degrader design in ligand-poor regimes. By integrating interaction intelligence, structural reasoning, and AI-based prioritization, SynGlue consistently narrows vast chemical spaces into actionable PROTAC candidates.

### Data-driven Module designed BRD4 PROTACs Exhibit Potent *in vitro* Activity and *in vivo* Anti-Tumor Efficacy

To evaluate the bioactivity of SynGlue data-driven module designed BRD4 PROTACS, i.e., BRD4 (cmp1) and BRD4 (cmp2), we first chemically synthesized them and validated using HPLC and ^1^H/^13^C NMR, respectively **(Supplementary Figure 12a-i).** First, the anti-cancer properties of SynGlue-designed BRD4 PROTACs were evaluated via anti-proliferation assays. BRD4 (cmp1) exhibited potent nanomolar activity (IC_50_: 1.1 nM in MV-4-11; 3 nM in VCaP), while BRD4 (cmp2) was less potent **(Figure 3a)**. Please note, the observed IC_50_ pattern is in line with the prediction from the SynGlue **(Figure 2)**. Specificity was underscored by its inactive stereoisomer, which showed significantly reduced potency (IC_50_: 1.4 µM in VCaP vs. 4 nM for BRD4 (cmp1)) **(Supplementary Figure 12a-b, f)**. Western blot analyses further confirmed potent, dose-dependent BRD4 degradation in prostate cancer lines (VCaP, and LNCaP) by both compounds **(Figure 3b-c)**. BRD4 (cmp1) also degraded BRD2 and BRD3 in these cells but showed selective BRD4 degradation over other isoforms in HeLa cells **(Figure 3c)**. In MV-4-11 cells, it achieved 50% degradation (DC_50_) at 0.193 nM, outperforming BRD4 (cmp2) (DC_50_ = 0.228 nM) **(Supplementary Figure 12c)**. Degradation was confirmed as ubiquitin-proteasome system-dependent, as co-treatment with the proteasome inhibitor MLN2238 rescued BRD4 levels **(Figure 3d)**. The inactive isomer showed minimal degradation **(Figure 3e; Supplementary Figure 12a)**. Based on this promising profile, BRD4 (cmp1) was advanced for *in vivo* evaluation. Intravenous pharmacokinetics in CD-1 mice showed dose-proportional exposure and a plasma half-life of ~0.83 hours **(Figure 3f; Supplementary Figure 12d; Supplementary Table 9)**. The compound was well-tolerated, with no adverse effects observed up to 10 mg/kg/day in a 14-day maximum tolerated dose study **(Figure 3g-h; Supplementary Table 9)**. In an MV-4-11 xenograft model, BRD4 (cmp1) induced statistically significant, dose-dependent tumor growth inhibition: 47% TGI at 5 mg/kg (q.d.), 120% TGI at 10 mg/kg (q.d.), and 75% TGI at 10 mg/kg (q48h) **(Figures 3i-l; Supplementary Figure 12e; Supplementary Table 9)**. The 10 mg/kg daily regimen demonstrated optimal tumor targeting, achieving a 2.2-fold higher concentration in tumor versus plasma (246 ± 35 ng/g vs. 110 ± 19 ng/g) **(Figure 3m)**. Anti-tumor efficacy correlated with *in vivo* BRD4 degradation and downregulation of c-Myc in excised tumors **(Figure 3n)**, with sustained target engagement confirmed up to 24 hours post-treatment **(Supplementary Table 9)**. In summary, the SynGlue-designed PROTAC BRD4 (cmp1) is a potent, selective degrader with favorable drug-like properties, pharmacokinetics, and tolerability, demonstrating significant mechanism-driven anti-tumor efficacy. These findings validate SynGlue’s ability to guide the discovery of therapeutically relevant multi-target molecules.

**Figure 3:**
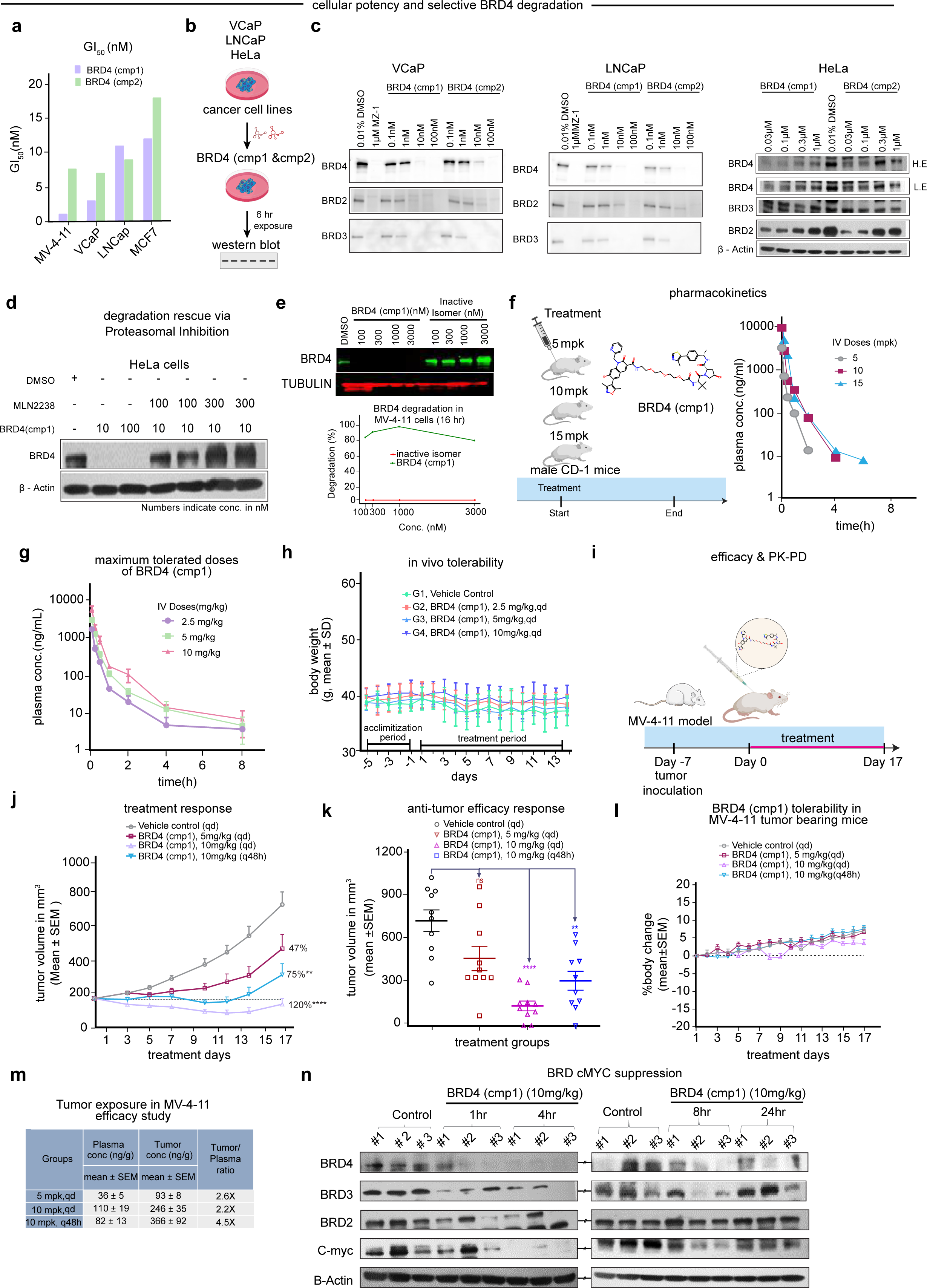
Preclinical Validation of SynGlue-Data Driven Module Designed BRD4 PROTACs. **(a)** A bar plot displays the Growth Inhibition 50 (GI_50_) values in nanomolar (nM) for SynGlue-designed PROTACs, BRD4 (cmp1) and BRD4 (cmp2), as determined by CellTiter-Glo® assay across various cancer cell lines, including MV-4-11, VCaP, LNCaP, and MCF7. **(b)** A schematic representation outlines the experimental strategy employed to assess the in vitro Bromodomain-containing protein 4 (BRD4) protein degradation potential of the SynGlue-designed PROTACs in VCaP, LNCaP, and HeLa cell lines following compound treatment for a specified duration (e.g., 6 hours). **(c)** Western blots illustrating the dose-dependent degradation of BRD4, BRD2, and BRD3 proteins in VCaP, LNCaP, and HeLa cell lines treated with increasing concentrations (0.1 nM to 100 nM) of BRD4 (cmp1) and BRD4 (cmp2) over 6 hours, with β-Actin or β-Tubulin serving as loading controls; the selective BRD4 degradation by BRD4 (cmp1) is particularly evident in HeLa cells. **(d)** A Western blot demonstrating the proteasomal degradation of BRD4 (cmp1)-mediated BRD4 degradation, showing that pre-treatment of HeLa cells with a proteasome inhibitor (e.g., MG132 or Bortezomib) followed by BRD4 (cmp1) treatment rescues BRD4 protein levels compared to treatment with BRD4 (cmp1) alone. **(e)** A Western blot illustrates the requirement for a functional PROTAC structure for degradation, as treatment of cells (e.g., HeLa) with an inactive isomer of BRD4 (cmp1) (AU-16914) at equivalent concentrations fails to induce BRD4 degradation compared to the active BRD4 (cmp1). **(f)** Schematic of the in vivo pharmacokinetic (PK) study design, outlining single intravenous (IV) administration of BRD4 (cmp1) in male CD-1 mice at 5, 10, and 15 mg/kg, with the corresponding plasma concentration time profiles (ng/mL versus hours) shown in line plots to determine PK parameters (C_max_, AUC, half-life). **(g)** Line plots showing the maximum tolerated doses (MTD) and PK profiles of BRD4 (cmp1), with consistent body weight maintenance across 2.5, 5, and 10 mg/kg/day IV doses over the 14-day administration period, indicating good tolerability. **(h)** Tolerability assessment of BRD4 (cmp1) in vivo, with line plots tracking average body weight (% change) of male CD-1 mice during the administration and treatment periods, confirming the absence of significant systemic toxicity at tested doses. **(i)** A schematic representation outlines the in vivo anti-tumor efficacy study design using a relevant xenograft model (e.g., MV-4-11 Cell Line Derived Xenograft (CDX) in athymic nude mice), including treatment groups (vehicle control, BRD4 (cmp1) at 10 mg/kg daily or alternate day IV), treatment duration (e.g., 17 days), and primary endpoints (tumor volume, body weight). **(j)** Line plots tracking tumor volume (mm^3^, mean ± SEM) in treated mice, demonstrating robust tumor growth inhibition by BRD4 (cmp1) relative to vehicle control, with dose-dependent efficacy. **(k)** A scatterplot summarizes the effect of BRD4 (cmp1) treatment on tumor volume (cubic millimeters (mm^3^), Mean ± Standard Error of the Mean (SEM)) at the study endpoint in the xenograft model, comparing vehicle control and BRD4 (cmp1) treatment groups, with statistical significance indicated. **(l)** Line plots of average body weight changes during the in vivo efficacy study, confirming tolerability across all treatment groups. Experimental setup for evaluating the efficacy of BRD4 (cmp1) in mice: Male athymic nude mice (7 to 8 weeks old, 10 mice per group) were inoculated with MV4-11 cells (ATCC) at a density of 15 million cells per mouse in 200LμL (1:1 HBSS & ECM gel). Treatment was initiated 8 days after inoculation, when the mean tumor volume reached approximately 170Lmm^3^, and continued for 17 days. Statistical analysis was performed using GraphPad Prism (version 10.2.3), applying Brown-Forsythe and Welch ANOVA tests. **(m)** Table summarizing the measured concentrations of BRD4 (cmp1) in plasma and tumor tissues at selected post-treatment time points in the efficacy study, demonstrating sustained intratumoral exposure and a favorable pharmacokinetic–pharmacodynamic (PK-PD) correlation. BRD4 (cmp1) exposure was assessed in plasma samples (nL=L9-10/group) and tumor tissues (nL=L8-10/group) at the 1-hour time point following the end of the efficacy study. **(n)** Representative Western blots show the protein levels of BRD4, BRD2, BRD3, and the downstream target c-Myc in tumor tissue lysates harvested from mice at baseline (pre-treatment) and at early time points (1 and 4 hours) post-treatment with BRD4 (cmp1), demonstrating pharmacodynamic target engagement and pathway modulation. Representative Western blots show the protein levels of BRD4, BRD2, BRD3, and c-Myc in tumor tissue lysates harvested from mice at later time points (6, 8, and 24 hours) post-treatment with BRD4 (cmp1), illustrating the duration of pharmacodynamic effects.

### Structure-guided Module designed GSPT1 PROTACs Exhibit Translational dependency across Cancer Models

GSPT1 (eRF3a), a core component of the translation termination machinery, has been considered undruggable due to its essential role. We hypothesized targeted degradation could selectively eliminate translationally addicted cancer cells. Using SynGlue’s structure-guided workflow, we designed and synthesized lead GSPT1 degraders. The chemical identity and purity of AU-29140 (GSPT1 (cmp1)) and AU-30489 (GSPT1 (cmp2)) were rigorously confirmed via HPLC, high-resolution mass spectrometry (HRMS), and ^1^H/^13^C NMR spectroscopy, yielding high chemical purity (>98%) and validating the synthetic routes **(Supplementary Figure 14a-h)**. We first evaluated the antiproliferative activity of GSPT1 (cmp1) across a diverse panel of hematologic and solid tumor cell lines. It displayed potent and broad growth inhibition, with sub-100 nM GI_50_ values in multiple models including Jurkat, 22Rv1, SW48, MDA-MB-436, CL-34, and HCC-1569 cells, consistent with a strong dependency on GSPT1-mediated translational control **(Figures 4a-b)**. Limited activity in lines such as DU145 and HepG2 indicated context-dependent vulnerability rather than nonspecific cytotoxicity, establishing GSPT1 (cmp1) as a selective, mechanistically driven degrader **(Supplementary Table 12a-c)**. To assess early molecular consequences, we performed TMT-based global proteomic profiling in 22Rv1 cells after 6-hour treatment with GSPT1 (cmp1). Quantitative analysis identified a subset of significantly altered proteins, with GSPT1 among the most downregulated, confirming robust target degradation and early perturbation of translational homeostasis pathways **(Figure 4c-d, Supplementary Table 13)**. Biochemical validation via Western blot in 22Rv1 cells demonstrated potent, dose-dependent degradation, with a calculated DC_50_ of ~1.1 nM and a Hill slope of 1.4 **(Figures 4e-f)**. Notably, degradation potency was uncoupled from antiproliferative activity, as substantial GSPT1 depletion occurred even in contexts with limited growth inhibition, highlighting a defining feature of degradation: protein elimination precedes biological collapse.

**Figure 4:**
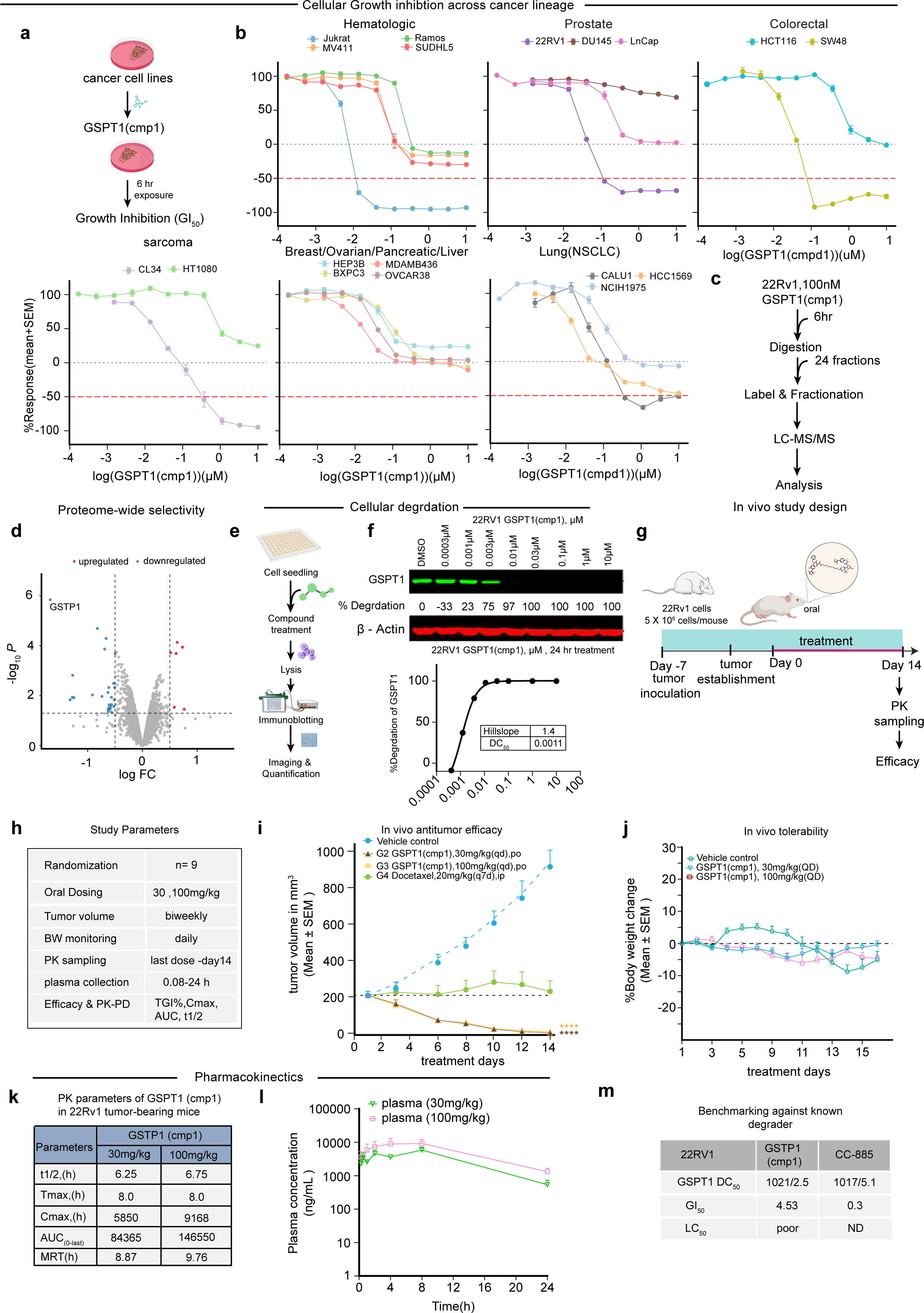
Experimental validation of structure-guided GSPT1 degradation reveals context-dependent translational vulnerability. **(a)** Schematic overview of the experimental workflow used to evaluate the antiproliferative effects of GSPT1 (cmp1) across cancer cell lines. Cells were treated with GSPT1 (cmp1) for 6 hours, followed by assessment of growth inhibition (GI_50_). **(b)** Dose-response growth inhibition profiles of GSPT1 (cmp1) across a diverse panel of hematologic and solid tumor cell lines, including hematologic, prostate, colorectal, sarcoma, breast/ovarian/pancreatic/liver, and lung (NSCLC) cancers. Highly sensitive cell lines exhibit deep cytotoxic responses, while resistant lines show limited growth inhibition, indicating context-dependent vulnerability. **(c)** Experimental design for TMT-based quantitative proteomics in 22Rv1 cells treated with GSPT1 (cmp1) (100nM, 6h), including digestion, labeling, fractionation, LC-MS/MS analysis, and downstream data processing. **(d)** Volcano plot of global proteomic changes in 22Rv1 cells following short-term treatment with GSPT1 (cmp1). Significantly upregulated and downregulated proteins are highlighted, with GSPT1 among the most significantly depleted targets. **(e)** Schematic of the biochemical workflow used to assess GSPT1 degradation by immunoblotting, including cell seeding, compound treatment, lysis, immunoblotting, and quantitative analysis. **(f)** Western blot analysis demonstrating dose-dependent degradation of GSPT1 in 22Rv1 cells following 24-hour treatment with GSPT1 (cmp1), with β-actin as a loading control. Quantification reveals near-complete target depletion at nanomolar concentrations, yielding a DC_50_ of ~1.1 nM with a Hill slope of 1.4. **(g)** In vivo study design illustrating oral administration of GSPT1 (cmp1) in 22Rv1 tumor-bearing mice, including treatment schedule, pharmacokinetic sampling, and efficacy evaluation. **(h)** Summary of key in vivo study parameters, including randomization, dosing regimen, tumor volume monitoring, body weight tracking, pharmacokinetic sampling, and efficacy endpoints. **(i)** Tumor growth curves for 22Rv1 xenografts treated with vehicle, GSPT1 (cmp1) (30 mg/kg and 100 mg/kg, oral), or docetaxel. GSPT1 (cmp1) induces robust, dose-dependent tumor growth inhibition compared with vehicle controls (**** p < 0.0001). **(j)** Body weight changes during the treatment period, demonstrating good tolerability of GSPT1 (cmp1) at both dose levels without significant treatment-related weight loss. **(k)** Pharmacokinetic parameters of GSPT1 (cmp1) in 22Rv1 tumor-bearing mice following oral administration, including half-life, C_max_, AUC, and mean residence time. **(l)** Plasma concentration-time profiles of GSPT1 (cmp1) following oral dosing at 30 mg/kg and 100 mg/kg, showing sustained systemic exposure and dose-proportional pharmacokinetics over 24 hours. **(m)** Comparative cellular activity of GSPT1 degraders in 22Rv1 cells. Despite similar GSPT1 degradation potency (DC_50_), AU-29140 (GSPT1 (cmp1)) exhibits reduced antiproliferative activity (GI_50_) and poor cytotoxicity (LC_50_) relative to CC-885, highlighting dissociation between degradation potency and cytotoxic effects.

We next assessed in vivo efficacy in 22Rv1 prostate cancer xenografts following oral administration. Treatment with GSPT1 (cmp1) resulted in robust, dose-dependent tumor growth inhibition at 30 mg/kg and 100 mg/kg compared to vehicle controls **(Figures 4g-i)**. Statistical analysis (Brown-Forsythe and Welch ANOVA) revealed highly significant treatment effects (p < 0.0001). The compound was well-tolerated over 14 days, with no significant body weight loss **(Figure 4j)**. Pharmacokinetic analysis confirmed favorable oral properties. GSPT1 (cmp1) exhibited sustained, dose-proportional exposure with moderate terminal half-lives (6.25 h at 30 mg/kg; 6.75 h at 100 mg/kg) and peak plasma concentrations at ~8 hours post-dose. AUC (0-last) values were 84,365 ng·h/mL (30 mg/kg) and 146,550 ng·h/mL (100 mg/kg), with mean residence times of 8.87-9.76 hours, supporting once-daily dosing **(Figure 4k)**. In tumor-bearing mice, the compound achieved dose-dependent plasma exposure, maintaining nanogram-per-milliliter concentrations for at least 24 hours **(Figure 4l)**. Finally, we benchmarked GSPT1 (cmp1) against the reference degrader CC-885 in 22Rv1 cells. While both exhibited comparable GSPT1 degradation potency (DC_50_ ≈ 1-2 nM), GSPT1 (cmp1) displayed markedly reduced antiproliferative activity (GI_50_ = 4.53 µM) and cytotoxicity compared to CC-885 (GI_50_ = 0.3 µM) **(Figure 4m)**. This dissociation underscores that GSPT1 (cmp1) induces selective, context-dependent effects rather than generalized toxicity. In summary, targeted degradation of GSPT1 induces selective proteome remodeling, drives cytotoxicity in translationally addicted cancers, and achieves durable tumor suppression in vivo with favorable tolerability and pharmacokinetics. These findings validate GSPT1 as a viable degradation target and demonstrate that SynGlue’s structure-guided pipeline can therapeutically exploit translational vulnerabilities previously considered inaccessible.

## DISCUSSION

The development of multi-targeting therapeutics, particularly via targeted protein degradation strategies such as PROTACs, represents a significant advancement in medicinal chemistry and pharmacology ^4,10,11,14^. These modalities hold the potential to address complex diseases more effectively than conventional single-target drugs by modulating multiple signaling pathways or eliminating disease-driving proteins. However, the rational design of potent, selective, and drug-like multi-target molecules remains a substantial bottleneck due to the intricate structure-activity relationships involved and the expansive chemical space, especially in the context of linker optimization and ternary complex formation ^4,14,16^. Bridging this gap requires a platform that integrates diverse data modalities and design principles.

In this study, we introduced SynGlue, a generative AI platform that unifies large-scale, data-driven fragment intelligence with structure-guided design to overcome these challenges. Its core innovation is the synergistic integration of two complementary modules within a single, decision-oriented framework. This architecture is powered by MagnetDB, one of the largest curated repositories of experimentally validated protein-ligand interactions explicitly designed for generative multi-target design. Coupled with a highly efficient TRIE-based indexing system, it enables rapid and interpretable fragment-centric target mapping, forming a robust foundation for *de novo* molecular assembly ^16,17^. This approach transitions from retrospective analysis to prospective design, allowing systematic navigation of vast chemical space while enforcing novelty and synthesizability. SynGlue’s performance was rigorously validated through extensive benchmarking, where it demonstrated superior target recovery and polypharmacology classification compared to existing tools like LigAdvisor ^25^, SwissTargetPrediction ^26^, and SuperPred ^27^. Its predictive core, integrating linker-aware multiclass models with transformer-based regressors for DC_50_ and D_max_, provides a quantitative bridge between molecular architecture and functional outcome, addressing a critical gap in degrader design.

The platform’s utility was demonstrated through its application to two orthogonal and clinically relevant targets: BRD4 and GSPT1. For BRD4, a well-characterized oncology target with extensive ligand data, SynGlue’s data-driven module enabled the discovery of novel, selective warheads. The resulting PROTAC, BRD4 (cmp1), exhibited potent degradation (DC_50_ = 0.19 nM) and selective anti-proliferative activity, which translated to significant tumor growth inhibition in vivo. This case validates SynGlue’s ability to achieve high-resolution prioritization and novelty in data-rich contexts, generating degraders with low structural similarity to known chemotypes. Conversely, for GSPT1, a translation termination factor with sparse direct ligand precedent, SynGlue’s structure-guided module enabled de novo warhead discovery directly from protein complex geometry. The designed degrader GSPT1 (cmp1) achieved sub-nanomolar predicted potency, induced selective proteome remodeling, and demonstrated robust oral efficacy in xenograft models. This success establishes that SynGlue can rationally address targets considered intractable to conventional discovery, exploiting therapeutic vulnerabilities via degradation. Existing computational tools often address isolated aspects of this pipeline, such as ternary complex modeling (e.g., PRosettaC ^21^) or linker design (e.g., DiffPROTACs ^22^). While valuable, they lack the integrated data scope, fragment-aware reasoning, and modular design flexibility necessary for end-to-end degrader discovery. SynGlue provides a holistic solution by unifying interaction intelligence, structural biophysics, and activity prediction within a generative framework, efficiently narrowing chemical space to experimentally actionable candidates.

Current limitations of SynGlue include the dependency of its data-driven module on the coverage of public interaction databases and the challenge of modeling highly rigid or macrocyclic scaffolds. Predictive accuracy for degradation endpoints is also constrained by the available public training data. Future iterations will expand MagnetDB with newly published data, refine fragmentation algorithms for complex chemotypes, integrate more advanced physics-based ternary complex modeling, and employ active learning cycles with experimental feedback to improve prediction robustness. Furthermore, incorporating assay metadata and uncertainty quantification will enhance the generalizability of predictions across cellular contexts. In conclusion, SynGlue establishes a generalizable, experimentally validated framework for the rational design of multi-targeted degraders. By demonstrating a complete pipeline from *de novo* design to in vivo efficacy for two distinct targets, BRD4 and GSPT1, we bridge a critical gap between computational innovation and therapeutic translation. SynGlue provides a scalable and versatile platform poised to accelerate the discovery of next-generation multi-target therapeutics.

## MATERIAL AND METHODS

### Construction and Preparation of the MagnetDB Interaction Database

MagnetDB was constructed by systematically aggregating and curating experimentally validated protein-ligand interaction data from six major public repositories: DrugBank ^28^ (v5.1.10), BindingDB ^29^ (downloaded 28-May-2023), ChEMBL^30^ (v33, released 31-May-2023), STITCH^31^ (v5.0), BioSNAP - ChG-InterDecagon, ChG-Miner, ChG-TargetDecagon (August 2018 release)^32^, and the Small Molecule Suite^33^. Database-specific extraction pipelines were applied, including XML parsing (DrugBank), SDF processing (BindingDB), API-based species-restricted queries (ChEMBL), network-based extraction (BioSNAP and STITCH), and internal mapping within the Small Molecule Suite, yielding 37.55 million raw interaction records. Ligands were standardized to canonical SMILES using OpenBabel (v3.0.0), and protein identifiers were harmonized to UniProt accessions. Redundant and inconsistent entries were resolved through automated normalization and manual curation, consolidating the dataset to 35.21 million unique ligand-target pairs. To ensure biological relevance, only experimentally validated, direct binding interactions were retained and filtered for Homo sapiens, Mus musculus, and Saccharomyces cerevisiae, reducing the dataset to 21.39 million species-specific entries. After removal of duplicates and low-confidence associations, the final curated MagnetDB comprises 6.37 million unique protein-ligand interactions, encompassing 1.94 million distinct compounds and 20,129 protein targets, each annotated with standardized identifiers, source information, assay details, and species. To enable scalable and chemically precise fragment mapping, we implemented a custom TRIE (prefix tree) indexed by reversed canonical SMILES strings with a terminal symbol ($). Compounds were fragmented using the RECAP algorithm in RDKit (v2023.09.02), and only terminal fragments representing single, synthetically meaningful cleavage events were retained, with attachment points marked by an asterisk (*). This strategy preserves key binding motifs and avoids ambiguities from non-terminal or multiple cleaved fragments. Terminal fragments were inserted into the TRIE and linked to MagnetDB interaction identifiers via hash maps. The TRIE preserves stereochemistry, bond order, and atom connectivity, enabling exact and prefix-based matching with O(n) complexity. In total, ~2.4 million fragments were indexed, and the TRIE was constructed in ~3.5 minutes on a standard CPU (Intel i9-10900X, 256 GB RAM). For downstream use, the TRIE and fragment-interaction mappings were serialized using Python pickle, enabling rapid structure-aware fragment retrieval within SynGlue.

### TRIE Performance Benchmarking

The TRIE data structure implemented in SynGlue was rigorously benchmarked to evaluate its efficiency and scalability, focusing on insertion time, search time, and prefix matching time. Two distinct fragment sets were employed for testing: (a) A random subset of 1,000 terminal fragments from MagnetDB, and (b) Fragments representing the 10^th^ and 90^th^ percentiles of structural diversity across MagnetDB’s constituent sources. For each dataset, average insertion and search times were recorded, and retrieval accuracy was evaluated using retrieval rate metrics. Empirical time complexity (Big O notation) was compared against theoretical expectations and brute-force substructure matching. Performance was further assessed across varying fragment lengths to evaluate structural scalability **(Figure 1c, Supplementary 1c-h)**. Performance was further evaluated across varying fragment lengths, confirming the TRIE’s scalability.

### SynGlue Data-Driven Analysis Workflow

SynGlue employs a data-driven, fragment-based pipeline to rapidly infer protein targets and polypharmacological profiles from input molecules. Query compounds, provided as SMILES, are canonicalized using OpenBabel (v3.0.0) and fragmented via RECAP in RDKit (v2023.09.02); the entire techstack is in **Supplementary Table 2**. Terminal fragments are mapped to MagnetDB using a TRIE-based exact-match search, enabling fast and scalable target association. To validate predicted interactions, SynGlue integrates multi-modal similarity scoring combining ECFP4 (1024-bit) and RDKit topological fingerprints (2048-bit) with Tanimoto and overlap coefficients, alongside maximum common substructure (MCS) analysis. Stringent filters-including minimum atom count (>10 atoms), MCS overlap thresholds (≥25% query, ≥75% database fragment), and exclusion of trivial fragments-ensure biological relevance and high-confidence matches. Based on fragment-target associations, compounds are classified into four polypharmacological categories (types 1-4), providing mechanistic insight and prioritization guidance. The final output includes matched fragments, associated targets, similarity scores, annotations, and classification metadata, enabling high-throughput, informed decision-making for multi-target drug design.

### Performance Evaluation and Comparative Benchmarking of Analysis Workflow

The SynGlue data-driven workflow was applied to a benchmark dataset of 6,935 compounds with known targets. Predicted targets were compared against ground truth targets (excluding known targets not present in MagnetDB for exact match analysis). Accuracy was quantified using hit rate and miss rate.

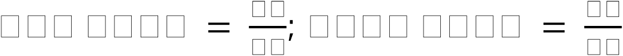

where;

TP = True Positives

FN = False Negatives

TT = Total Known Targets

Performance was assessed across different compound categories (Drug Central, PROTAC, and Others) **(Supplementary Figure 2d)**. SynGlue was further benchmarked against LigAdvisor, SwissTargetPrediction, and SuperPred using a representative subset of 100 structurally diverse molecules (selected via ECFP4 clustering) **(Supplementary Figures h-j)** for clustering methodology. Target predictions were obtained from all tools for this subset. These metrics enabled direct comparison of performance across all tools.

### SynGlue Generative Design Workflow

The SynGlue Generator enables de novo PROTAC and MTM design using the Link-INVENT architecture, an encoder-decoder RNN with LSTM units (embedding size 256; three hidden layers of 512 units) trained via reinforcement learning. Warhead prioritization filters MagnetDB fragments using physicochemical and structural constraints (MW 200-500 Da, LogP 0-5, TPSA 20-120 Å^2^, rotatable bonds 0-10, rings 0-5, aromatic proportion <0.45, SA ≤6, Csp^3^ 0.3-0.8) and required reactive groups (Cl, Br, I, NH_2_), ensuring drug-like and chemically tractable warheads. Prioritized warheads are paired with compatible E3 ligases from PROTAC-DB using property-matched filtering, typically yielding 5-30 warhead-E3 pairs per input. The exit vector defines the chemically viable attachment point on the warhead that preserves key target-binding interactions while enabling linker conjugation. In SynGlue, exit vectors are selected based on fragment attachment sites, steric accessibility, and compatibility with ternary complex geometry to ensure productive PROTAC assembly. For each pair, SynGlue generates linkers optimized using a multi-objective scoring function that balances hydrogen bond donors, ring content, and graph length, promoting flexibility. conformational adaptability, and ternary complex compatibility. High-scoring designs (score ≥0.8) are annotated with unique identifiers, SMILES, scaffolds, computed physicochemical properties, and design metadata. Generative runs typically produce 2,000-4,000 PROTACs per warhead-E3 pair within ~10 minutes on a 12-core CPU (128 GB RAM). SynGlue’s modular design supports future integration of additional generative constraints, such as ternary complex energetics, off-target prediction, or disease-context filtering, enabling scalable and extensible multi-functional therapeutic design.

### SynGlue Data Driven Used for Multitarget Molecules

SynGlue’s data-driven design module streamlines the creation of multitarget molecules (MTMs) by leveraging MagnetDB-derived fragments that demonstrate a high degree of matching (fragment coverage over 75%) to target binding sites, ensuring strong baseline affinity. These candidate warheads are further refined through a stringent filtering process incorporating critical drug-likeness and developability parameters, including molecular weight (200-500 Da), logP (0-5), topological polar surface area (TPSA, 20-120 Å^2^), number of rotatable bonds (0-10), aromatic ring content (<0.45), synthetic accessibility (SA ≤ 6), and Csp^3^ fraction (0.3-0.8). This comprehensive profile ensures that the selected warheads possess optimal solubility, permeability, and synthetic feasibility while maintaining a balanced conformational flexibility essential for engaging multiple binding sites simultaneously. After the target warhead selection, SynGlue’s Generator module is used for the design and optimization of linkers to maintain the ideal spatial arrangement, chemical stability, and flexibility needed for robust multitarget binding. This integrated approach maximizes cooperative interactions and enhances the functional synergy of the resulting MTMs. The effectiveness of this workflow has been demonstrated in case studies such as CDK1-EGFR and PRKX-DRD1 multitarget molecules **(Supplementary 6a-b)**, where SynGlue’s data-driven methodology enabled rapid prototyping of MTMs with tailored therapeutic profiles by combining predictive analytics with precise structural alignment.

### SynGlue Structure-Guided Design Workflow

For targets that lack sufficient data coverage in MagnetDB, such as novel or less-studied protein targets, SynGlue employs a complementary structure-guided workflow. This approach leverages curated or predicted 3D protein structures (from RCSB PDB) processed using standard protein preparation pipelines, followed by Gcoupler to identify relevant pharmacophores in druggable binding pockets and ligand design. The structure-guided design is particularly valuable in these data-sparse scenarios because it leverages three-dimensional pharmacophore complementarity to pinpoint targetable features and binding hot spots that might be invisible to purely ligand-based approaches. These pharmacophores guide the *de novo* design, leading to the selection of compatible warheads based on structural complementarity and properties (molecular weight, LogP, halogenation, heteroatoms, and TPSA). Selected warheads are paired with compatible E3 ligases (as in the data-driven workflow). The SynGlue Generator designs linkers that respect the geometric and steric constraints of the identified pockets, ensuring that final candidates are not only synthetically feasible but also structurally compatible with the target’s unique binding landscape. Final candidates can be refined through molecular docking (AutoDock Vina v1.2) and geometric analyses (RMSD ≤ 2.0 Å from pharmacophore sites) to confirm proper binding orientations and energy. This workflow was applied in the AR-XIAP, BRD4-VHL, GSPT1-CRBN case studies using RCSB PDB and Gcoupler-based pharmacophore models to generate high-confidence candidates. Final molecules are output as SMILES, accompanied by design and scoring metadata for downstream prioritization.

### Predictive Prioritization of Designed Compounds

To systematically rank designed BRD4-VHL PROTACs, predictive models for degradation potency were developed: (1) A regression-based attention learning model estimates DC_50_ and D_max_ values using molecular representations and structural embeddings. (2) A linker-based classification model predicts the DC_50_ activity class (high, medium, low) and categorizes linker potency based on the DC_50_ quartiles. The regression framework employed a component-aware attention model, wherein GROVER-derived graph embeddings (4800-dimensional) were independently generated for each PROTAC component: warhead, linker, and E3 ligase. An attention mechanism was implemented to learn component-wise weights reflecting the relative contribution of each substructure to overall degradation efficacy. These attention-weighted embeddings were used with a separate GROVER embedding of the full PROTAC molecule and passed through a deep feedforward regressor consisting of three dense layers (512, 256, and 128 units) with ReLU activations and dropout (rate = 0.2). The model was trained using mean squared error loss with the Adam optimizer over 200 epochs with early stopping. For comparative analysis, we also trained multiple traditional regressors, including Random Forest, Gradient Boosting, AdaBoost, Ridge, Lasso, ElasticNet, Support Vector Regression, k-Nearest Neighbors, XGBoost, Multilayer Perceptron, Stochastic Gradient Descent, and Decision Trees, using the same GROVER embeddings as input. All regressors were implemented using scikit-learn (v1.3.0), XGBoost (v1.7+), and LightGBM (v3.3+) and evaluated using ten-fold cross-validation with R^2^, MAE, and RMSE as performance metrics. To balance expressive power and generalization, we employ a hybrid framework in which attention-based deep learning captures component-specific interactions, followed by classical machine-learning models that provide robust prediction under limited and noisy PROTAC data.

To complement continuous prediction and enable rapid ranking of linker designs, we constructed a multiclass classification model for potency class prediction, using a curated dataset of 2,320 PROTACs with experimentally validated DC_50_ values. Linkers were grouped into four potency classes (low, medium-low, medium-high, and high) based on DC_50_ quartile thresholds (Q1 = 6.28 ng/mL, Q2 = 54.96 ng/mL, Q3 = 585 ng/mL). Feature extraction was performed using five orthogonal descriptor sets: bioactivity-based Signaturizer (v1.1.14) ^34^, graph-based GROVER (https://github.com/tencent-ailab/grover) ^35^, physicochemical Mordred (1.2.0) ^36^, image-based ImageMol (https://github.com/ChengF-Lab/ImageMol) ^37^, and language model-based embeddings from ChemBERTa (ChemBERTa-77M-MLM) ^38^, trained as a multi-class classifier **(Supplementary Figure 4a)**. After data preprocessing and an 80:20 stratified train-test split, feature selection was performed using the Boruta algorithm to identify the most informative variables. A suite of twelve classifiers, including SVM, Logistic Regression, kNN, Random Forest, Extra Trees, SGD, Gradient Boosting, XGBoost, Gaussian Naive Bayes, Multilayer Perceptron, AdaBoost, and Decision Trees were trained and evaluated using one-vs-all AUC-ROC, accuracy, and F1-score. Hyperparameter tuning was conducted via GridSearchCV from the scikit-learn (v1.5.2) library, and final models were validated using 10-fold cross-validation. Class probabilities were used to prioritize linkers predicted to fall into high-potency classes with high confidence, supporting targeted design decisions. **(Supplementary Figure 4b-k).** We acknowledge that DC_50_ and D_max_ values are inherently assay-dependent and can vary across cell lines, experimental protocols, and data sources; thus, while our predictive models demonstrate robust performance within the available training distribution, their generalizability may be influenced by inter-study variability in degradation assays.

### Chemical Space Analysis of Designed Compounds

The chemical diversity of SynGlue-designed PROTACs targeting the androgen receptor (AR) was evaluated relative to known AR PROTACs. ECFP6 fingerprints (radius 3, 2048 bits) were computed using RDKit (v2019.03.1), and principal component analysis (PCA) was performed on the binary fingerprint matrix using scikit-learn (v1.5.2) to compare the distribution, novelty, and chemical-space coverage of designed versus known compounds **(Figure 2a, 2i)**.

### Novel BRD4 Warhead Discovery Pipeline

To identify potent, selective, and structurally novel warheads for BRD4, we implemented a multi-step filtering pipeline combining data-driven selection with stringent physicochemical and synthetic feasibility criteria. We began by retrieving BRD4-associated fragments from MagnetDB and retained those with a high matching percentage (>75% fragment coverage). To ensure specificity, we included related bromodomain targets such as BRD2, BRD3, and BRD9, and removed any overlapping fragments (Tanimoto similarity ≥ 0.9 using ECFP4), thereby isolating BRD4-specific candidates. The filtered warheads were then evaluated for favorable drug-like properties, retaining those with logP values between 0 and 5 (indicating balanced solubility and permeability), QED scores ≥ 0.5 (suggesting drug-likeness), and Synthetic Accessibility (SA) scores ≤ 5 (indicating ease of synthesis). To further ensure novelty, we sourced known warheads for BRD family targets from PROTACDB (3.0) and computed ECFP4-based Tanimoto similarity against our MagnetDB-derived BRD4 candidates **(Supplementary Figure 9a)**. Only those fragments with similarity scores below 0.5 were retained, ensuring that the final set represented structurally novel BRD4 warheads. Finally, we performed hierarchical clustering and dendrogram analysis to visualize chemical diversity and identify distinct warhead families, from which we selected the most promising novel candidates **(Supplementary Figure 9b)**. These candidates were further optimized by assessing synthetic feasibility using RDKit-based synthetic accessibility scoring and structure optimization.

### Structural Similarity Benchmarking of In-House BRD4 PROTACs

Two in-house BRD4 PROTACs (cmp1 and cmp2) were compared against known BRD4 degraders curated from PROTAC-DB 3.0 and PROTACpedia. All structures were standardized to canonical SMILES, and structural similarity was quantified using Tanimoto similarity computed on Morgan fingerprints (radius 2, 2048 bits). Similarity distributions were visualized using violin and kernel density plots, and the highest-similarity reference compound for each query was identified and highlighted with its corresponding similarity score and 2D structure **(Supplementary Figure 10b)**.

### Structure Guided Module for GSPT1

Structure-based design of GSPT1 degraders was performed using the CRBN-GSPT1 crystal structure (PDB: 5HXB), which contains a cereblon modulator bound to GSPT1. In this structure, chains A and X both correspond to GSPT1 and exhibit highly similar sequences and conformations. Based on sequence and residue-level analysis, chain X was selected as the representative GSPT1 structure for cavity identification and de novo warhead design, while the full CRBN-GSPT1 complex was retained to preserve biologically relevant interaction geometry. The presence of a bound cereblon modulator supported CRBN selection as the E3 ligase. Binding cavities on GSPT1 were identified using an internal cavity-detection module and used to guide de novo warhead generation. Optimized GSPT1 warheads were subsequently integrated into a PROTAC design workflow to identify compatible E3 ligase recruiters and linker architectures that promote efficient ternary complex formation. PROTAC-DB ligands were filtered based on physicochemical properties, reported binding affinity (Kd/IC_50_), attachment points, synthetic feasibility, TPSA, and logP, leading to prioritization of cereblon ligands and selection of the top three candidates. Warheads were combinatorially paired with CRBN ligands and diversified through linker optimization, exploring linker length, flexibility, and attachment geometry. Generated PROTACs were filtered using predefined property windows (MW 850-1100 Da; logP 3-6, allowed up to 7; halogen count ≤3; heteroatoms ≤22; TPSA <230 Å²) and evaluated using a linker-based scoring framework incorporating binding complementarity, conformational feasibility, and steric compatibility at the GSPT1-CRBN interface. Candidates were prioritized based on predicted DC_50_ and D_max_ followed by ADME-based triage and molecular docking. Iterative optimization was performed until convergence on PROTACs with balanced degradation efficiency, physicochemical properties, and translational potential.

### Ternary Complex Modelling

Ternary complex models for BRD4 (cmp1) and GSPT1 (cmp1) **(Figure 2g and 2n)** were generated using Aurigene Oncology’s proprietary bifunctional degrader modelling platform, ALMOND <u>(Samajdar 2021)</u>, and the commercial ICM PROTAC Modeller (MolSoft). Structural assemblies and molecular visualizations were rendered using PyMOL (Schrödinger). The ALMOND workflow is a validated protocol for precision modelling of bifunctional degraders and has been previously applied across multiple target classes, including SMARCA2/4 <u>(Xiao et al.</u> <u>2024) (He et al. 2024)</u> and AR-V7 <u>(Bhumireddy et al. 2022)</u>.

### Computational Tool Benchmarking

To compare contemporary computational platforms used in PROTAC design and evaluation, a structured benchmarking framework was applied across seven widely used tools. Each tool was examined based on publicly described features, including its primary function, underlying computational methodology, databases or training sets used, component sourcing strategy, potency assessment approach, and final output format. Information was systematically extracted from published documentation, associated databases, and original tool descriptions and then organized into a unified comparative table **(Supplementary Fig 7c).** This allowed qualitative assessment of methodological diversity from data-driven transformer models and GNN-based scoring systems to physics-based docking workflows, as well as differences in input requirements, predictive capabilities, and expected outputs across tools.

### Evaluation of SynGlue and PDP Predictive Performance

SynGlue and the PDP tool were benchmarked for PROTAC potency prediction in VCaP and LNCaP cells by comparing predicted rankings with experimental potency and western blot degradation profiles. SynGlue outputs quantitative DC_50_ and D_max_ values, enabling direct comparison with biochemical measurements, whereas the PDP tool provides normalized activity scores (0-1) without dose-response resolution. Ranking accuracy was assessed by concordance with experimental ordering (BRD4 cmp1 > cmp2) and correlation to western blot data by agreement in degradation direction and magnitude. SynGlue showed higher interpretability due to its mechanistic outputs, while the PDP tool provided only moderate interpretability owing to its non-quantitative scoring.

### Chemical Synthesis and Characterization

Compounds for BRD4 (cmp1) (AU-14555), BRD4 (cmp2) (AU-14849), and an Inactive Isomer (AU-16914, or negative control) were synthesized via multi-step procedures involving peptide couplings (HATU/DIPEA), protecting group manipulations (Boc deprotection with TFA), and purification by CombiFlash® chromatography. Detailed procedures and intermediate steps are provided^39^ **(Supplementary Note 1)**. All non-aqueous reactions were performed under nitrogen in oven-dried glassware. Reaction progress was monitored by TLC (Merck silica gel 60 F254, UV 254 nm). Reagents and solvents were commercial grade. Structural elucidation used NMR (^1^H at 400 MHz, ^13^C at 101 MHz, DMSO-d^6^, TMS internal standard for ^1^H). LC-MS used an Agilent 1100-LC/MSD VL (0.1% formic acid in ACN or water/MeOH). HRMS used a Thermo Orbitrap Q-Exactive Plus LC-MS/MS (ESI). Final purity was confirmed by HPLC (BRD4 (cmp1): 98.37%; BRD4 (cmp2) 98.26%; Inactive Isomer: 97.69%). The inactive isomer of BRD4 (cmp1) (AU-16914) was synthesized as a stereochemical negative control to confirm that the observed biological activity is driven by the specific configuration of the active compound. Additional compounds for GSPT1 (GSPT1 (cmp1) AU-29140, GSPT1 (cmp2) AU-30489) evaluated in downstream biological studies were synthesized and characterized using analogous synthetic strategies, as described in **(Supplementary Note 1)**.

### Cell culture

VCaP (Vertebral-Cancer of the Prostate), LNCaP (Lymph Node Carcinoma of the Prostate), HeLa (cells of Henrietta Lacks), and MV-4-11 (MonoMac Variant 4-11 cells) cell lines were purchased from the American Type Culture Collection (ATCC, USA). VCaP cells were maintained in Dulbecco’s modified Eagle’s medium (DMEM) supplemented with 10% (v/v) fetal bovine serum (FBS); LNCaP cells were cultured in RPMI-1640 medium supplemented with 10% FBS; MV-4-11 cells were maintained in Iscove’s Modified Dulbecco’s Medium with 10% FBS; HeLa and MCF7 cells were maintained in Eagle’s Minimum Essential Medium with 10% FBS. All the cells were cultured at 37 °C in a humidified atmosphere with 5% CO_2_. For GSPT1 additional cancer cell lines used for extended antiproliferative profiling are listed in **(Supplementary Table 12)**. Human cancer cell lines including Jurkat, 22Rv1, SW48, HT1080, MV-4-11, OVCAR-8, BxPC-3, NCI-H1975, Ramos, LNCaP C4-2B4, Hep-3B, DU145, SU-DHL-5, HepG2, HCT116, MDA-MB-436, CL-34, HCC-1569, and Calu-1 were cultured under standard conditions at 37°C in a humidified incubator with 5% COL. Cells were maintained in their respective culture media supplemented with 10-20% fetal bovine serum (FBS) as detailed in **Supplementary Table 12**. Cells were routinely tested to ensure viability and appropriate growth characteristics prior to experimental use.

### Proliferation assay

LNCaP, VCaP, HeLa cells, and MV-4-11 were seeded in 96-well black polystyrene microplates (Corning, Cat #CLS3603) and treated with BRD4 (cmp1) and BRD4 (cmp2) at a final DMSO concentration of 0.1%. Cell viability was assessed using the CellTiter-Glo® Luminescent Cell Viability Assay. The GI_50_ values were calculated based on luminescence measurements, indicating the concentration at which 50% growth inhibition was observed. Western blot analysis of BRD4, BRD3, and BRD2 expression was performed in VCaP, LNCaP, HeLa, and MV-4-11 cell lines following treatment with BRD4 (cmp1) and BRD4 (cmp2) across a concentration range (0.1 nM to 1000 nM). For GSPT1 degradation studies, 22Rv1 cells were treated with test compounds under similar conditions and analyzed by Western blot to assess target degradation. Anti-proliferative activity of GSPT1 (cmp1) was evaluated using the CellTiter-Glo (CTG) luminescent viability assay. Cells were seeded in 96-well plates at optimized densities specific to each cell line and allowed to equilibrate prior to compound treatment. GSPT1 (cmp1) was tested using an 11-point, three-fold serial dilution, with 10 µM as the highest concentration, in triplicate wells. The final DMSO concentration was maintained at 0.5%. Following compound treatment, cells were incubated for 3-7 days depending on cell line-specific growth kinetics. CTG reagent was added according to the manufacturer’s instructions, and luminescence was measured using a multimode plate reader. Blank-subtracted luminescence values were used to calculate GI_50_ values by nonlinear regression using GraphPad Prism.

### Western Blotting

For Western blotting, cell lysates were prepared in RIPA buffer (ThermoFisher, #89900) supplemented with a protease inhibitor cocktail (Sigma-Aldrich, #P8340), and total protein was quantified using a Pierce BCA Assay (ThermoFisher, #23225). An equal quantity of protein from each sample was separated by 8% SDS-PAGE and transferred to polyvinylidene difluoride (PVDF) membranes. Membranes were probed with primary antibodies specific for BRD2 (Cell Signaling Technology [CST], #5848), BRD3 (Santa Cruz Biotechnology, #sc81202), BRD4 (CST, #13440), and c-Myc (CST, #5605) and GSPT1/eRF3 (CST, #14980S) β-Actin (CST, #3700S) or β-Tubulin (CST, #86298) was used as a loading control. For detection, membranes were incubated with either HRP-conjugated secondary antibodies (CST, #7074 and #7076), followed by enhanced chemiluminescence development, or with near-infrared IRDye® secondary antibodies (LI-COR, #926-68070 and #926-32211) before being imaged on an Odyssey Imaging System (LI-COR Biosciences). 22Rv1 cells were seeded in 6-well plates at a density of 1.5 × 10L cells per well and allowed to attach overnight. Cells were treated with GSPT1 (cmp1) across an 8-point dose range (10 µM to 0.003 µM) for 24 hours. Following treatment, cells were lysed using RIPA buffer supplemented with protease and phosphatase inhibitors. Lysates were clarified by centrifugation, and protein concentrations were determined using a BCA assay. Equal amounts of protein (40 µg) were resolved by 10% SDS-PAGE, transferred onto PVDF membranes, and blocked prior to incubation with primary antibodies against GSPT1 (eRF3) and β-actin. After incubation with fluorescent secondary antibodies, membranes were imaged using a LI-COR Odyssey system, and band intensities were quantified using Image Studio software. GSPT1 levels were normalized to β-actin and expressed as percentage degradation relative to control.

### *In Vivo* Pharmacokinetics (PK) and Pharmacology Studies

Pharmacokinetic studies used male CD-1 mice (7-9 weeks). For IV (3 mg/kg): BRD4 (cmp1) was formulated by dissolving in 2% v/v DMA + 20% w/v hydroxy beta cyclodextrin in water (pH ~7). For IV dose escalation (5, 10, 15 mg/kg): BRD4 (cmp1) dissolved in 10% v/v PEG-400 + 3% w/v Cremophor EL + 87% v/v PBS. Dose administration was done via the tail vein (IV). Blood was collected via the submandibular vein at specified time points (0.083-24 h). Plasma is isolated and stored at −80°C. BRD4 (cmp1) was quantified using a fit-for-purpose LC-MS/MS method using Carbamazepine as an internal standard (calibration range 5-1000 ng/mL). PK parameters were calculated using non-compartmental analysis (Phoenix WinNonlin). Pharmacology studies were conducted using both BRD4 and GSPT1 xenograft models. A M-V-4-11 cell line-derived xenograft (CDX) model in male athymic nude mice. Cells were injected SC (15 × 10L cells in HBSS: ECM gel). Treatment was initiated at tumor volumes of ~150-200 mm^3^. Mice (n=10/group) received BRD4 (cmp1) IV (5 mg/kg QD, 10 mg/kg QD, or 10 mg/kg QAD) for 17 days. Tumor volumes (TV = length × width^2^ × 0.5) were measured thrice weekly, and body weights were measured daily. Tumor growth inhibition (TGI%) calculated: %TGI = 100 – ((ΔT/ΔC) × 100). For mechanism-of-action studies, tumor tissues were harvested at study termination, snap-frozen, pulverized, lysed (CST lysis buffer), and analyzed via Western blot for BRD2, BRD3, BRD4, c-Myc, along with β-actin as a loading control. A 14-day MTD study used male CD-1 mice (n=6/group) which received BRD4 (cmp1) IV at 2.5, 5, and 10 mg/kg/day (formulation: 10% PEG-400, 3% Cremophor EL, 87% PBS), monitored for clinical signs, and measured body weight. Hematology, biochemistry, and histopathology were performed post-study.

### GSPT1 Pharmacology study

All in vivo pharmacology and pharmacokinetic studies were conducted in accordance with protocols approved by the Institutional Animal Ethics Committee (IAEC) of Aurigene Oncology Ltd. and in compliance with CPCSEA (Government of India) guidelines. The GSPT1 degrader AU-29140 aka GSPT1 (cmpd1) was formulated for oral administration in a vehicle consisting of 6% (v/v) dimethyl acetamide, 59% (v/v) Labrasol ALF, 25% (v/v) Labrafil M1944 CS, and 10% (v/v) Capryol 90. Human prostate cancer 22Rv1 cells (ATCC) were cultured in RPMI-1640 medium supplemented with glucose, sodium bicarbonate, sodium pyruvate, fetal bovine serum, and penicillin–streptomycin, harvested during exponential growth, and resuspended in a 1:2 mixture of Hank’s Balanced Salt Solution and extracellular matrix gel. Tumors were established by subcutaneous injection of 5 × 10^6^ cells per mouse into the right flank of 7-week-old male athymic nude mice. Treatment was initiated when mean tumor volumes reached approximately 150-250 mm^3^, after which animals were randomized into treatment groups (n = 9 per group). GSPT1 (cmp1) was administered orally at 30 and 100 mg/kg once daily at a dose volume of 10 mL/kg for 14 consecutive days. Body weights were monitored daily, and tumor volumes were measured at least twice weekly using digital calipers, with tumor volume calculated as length × width^2^ × 0.5. Tumor growth inhibition was calculated as %TGI = 100 − ((ΔT/ΔC) × 100), where ΔT and ΔC represent changes in mean tumor volume from the start of treatment for treated and control groups, respectively. Pharmacokinetic evaluation of GSPT1 (cmpd1) was conducted following oral dosing on the final day of treatment. Blood samples were collected at 0.08, 0.25, 0.5, 1, 2, 4, 8, and 24 hours post-dose. Plasma concentrations were quantified using a validated LC-MS/MS method, and pharmacokinetic parameters were derived using non-compartmental analysis.

### TMT-Based Global Proteomics Analysis

Human prostate cancer 22Rv1 cells were treated with 100 nM GSPT1 (cmp1) or DMSO (vehicle control) for 6 h in biological duplicates, after which cells were harvested and lysed in 0.4 mL lysis buffer using pulse sonication (Fisher Scientific Sonic Dismembrator Model 500; 15% amplitude with 10 s rest intervals). Lysates were clarified by centrifugation at 15,000 × g for 10 min at 4°C, and protein concentrations were determined using a BCA reducing reagent-compatible assay kit (Thermo Fisher Scientific). For proteomic analysis, 50 µg of protein from each sample was reduced with TCEP, alkylated with iodoacetamide for 30 min at room temperature in the dark, and digested with trypsin at 37°C overnight (16 h). Resulting peptides were labeled with TMT reagents according to the manufacturer’s instructions and fractionated into 24 fractions by basic reverse-phase liquid chromatography on an Agilent 1290 Infinity UHPLC system using a Poroshell HPH-C18 column (2.1 × 150 mm, 2.7 µm). Each fraction was loaded onto a peptide trap cartridge and separated on a 25 cm reversed-phase C18 Easy-Spray column (Thermo Fisher) using a 110 min linear gradient of 3-36% acetonitrile in 0.1% formic acid at a flow rate of 0.3 µL/min. Peptides were ionized with a Nano EasySpray source (spray voltage 1.8 kV, capillary temperature 275°C) and analyzed on an Orbitrap Exploris 240 mass spectrometer operated in data-dependent acquisition mode. Full MS scans (m/z 350–1800) were acquired at 35,000 resolution with an AGC target of 3 × 10^6^ and a maximum injection time of 100 ms, followed by HCD MS/MS of the top 20 multiply charged precursor ions (z ≥ 2) at a normalized collision energy of 30, acquired at 17,500 resolution with an AGC target of 1 × 10^5^ and a maximum injection time of 400 ms; dynamic exclusion was set to 20 s, with exclusion of unassigned and ≥7+ charge states. Raw data were searched against the UniProtKB human protein database using Proteome Discoverer (v2.5), and differential protein abundance was assessed using the MSqROB linear modeling framework with Benjamini-Hochberg correction for multiple testing. Proteins with an adjusted p-value < 0.05 and a fold change ≥ 1.25 between GSPT1 (cmp1)-treated and control samples were considered differentially expressed.

### Statistical analysis

All statistical analyses were performed using the statistical packages provided within the R and Python programming frameworks, along with Past-4 and GraphPad PRISM software. The p-value cutoff for significance was 0.05. Significance levels are * < 0.05, ** < 0.01, *** < 0.001, and **** < 0.0001. Statistical analysis was performed using GraphPad Prism (v10.2.3). Tumor volume comparisons were analyzed using the Brown-Forsythe and Welch ANOVA tests to account for unequal variances. Data are presented as mean ± standard error of the mean (SEM), and significance was defined at *P*L<L0.05. No statistical method was used to predetermine the sample size. The experiments were not randomized. The investigators were not blinded to allocation during experiments and outcome assessment. No statistical methods were used to predetermine sample sizes. Data distribution was assumed to be normal, but this was not formally tested. Data collection and analysis were not performed blind to the conditions of the experiments.

## Data Availability

The source code of SynGlue is available on the GitHub project: https://github.com/the-ahuja-lab/SynGlue. The validation datasets of PROTACs (PROTAC-DB 3.0, PROTACpedia), DrugCentral, and Literature are accessible in the Supplementary tables. The study utilized data from DrugBank 24 (v5.1.10), BindingDB (downloaded 28-May-2023), ChEMBL (v33, released 31-May-2023), STITCH (v5.0), BioSNAP - ChG-InterDecagon, ChG-Miner, ChG-TargetDecagon (August 2018 release), the Small Molecule Suite (Harvard, 2019), and PROTAC-DB 3.0.

## Code Availability

A Python package for SynGlue is provided via pip: https://test.pypi.org/project/SynGlue/. SynGlue is free for academic institutions; however, a commercial license key is mandatory for commercial usage. The source code of SynGlue is available on the project GitHub page: https://github.com/the-ahuja-lab/SynGlue.

## Ethics Statement

All animal experiments were conducted in accordance with protocols approved by the Institutional Animal Ethics Committee (IAEC) of Aurigene Oncology Ltd. and complied with the guidelines of the Committee for Control and Supervision of Experiments on Animals (CPCSEA), Government of India.

## Acknowledgments

The authors thank the IT-HelpDesk team of IIIT-Delhi for assisting with the computational resources. We thank all the members of the Ahuja lab for their intellectual contributions at various stages of this project. The Ahuja lab is supported by the Ramalingaswami Re-entry Fellowship (BT/HRD/35/02/2006), a re-entry scheme of the Department of Biotechnology, Ministry of Science & Technology, Government of India, and an intramural Start-up grant from Indraprastha Institute of Information Technology-Delhi. Saveena Solanki, Sanjay Kumar Mohanty, are supported by the INSPIRE fellowship from the Department of Science & Technology, India.

## Author Contributions

G.A. designed the SynGlue Computational Workflow, initiated the computational part of the project, and supervised the overall study. S.S. (Susanta Samajdar) designed, planned, and ensured the smooth execution of all experimental validations. S.S. (Saveena Solanki) spearheaded the SynGlue workflow and designed and coded the entire codebase. S.K.M. assisted in code refinement and implementation. S.S. (Shiva Satija) and S.C. assisted with SynGlue pipeline validation, testing, and MagnetDB data compilation. N.V.M.R.B., S.D. (Sandeep Dukare), N.K.T., N.K.R., and A.A.B. performed all the experiments under the guidance of S.S. (Susanta Samajdar). D.C., W.B., and S.R.S. helped with GSPT1 experiments. V.G., S.A., S.K., S.D. (Subhadeep Duari), A.S., and R.S. assisted in SynGlue testing and manuscript editing. D.S. assisted with algorithm implementation and manuscript editing. C.A. assisted in all wet lab experiment execution, data compilation, and figure making. All authors have read and approved the final manuscript.

## Supplementary Figure Legends

**Supplementary Figure 1: MagnetDB Fragmentation, TRIE Characteristics, and Workflow Schematic**

**(a)** The stepwise workflow of MagnetDB construction is shown, including data ingestion from multiple ligand-target sources, RECAP-based hierarchical fragmentation, terminal fragment selection, and TRIE indexing. **(b)** A schematic illustrates the data-driven query workflow, where canonicalized SMILES are mapped onto the MagnetDB TRIE followed by similarity-based matching and annotation. **(c)** Pie charts show the proportional contribution of terminal fragments from individual source databases. **(d)** A table summarizes MagnetDB statistics, including the number of unique compounds, terminal fragments, and associated targets across databases. **(e)** A complexity table compares insertion, search, and deletion operations for TRIE-based indexing versus brute-force search. **(f)** A scatter plot depicts fragment count as a function of fragment length, highlighting traversal frequency trends. **(g)** A line plot shows fragment-wise search time across indexed fragments. **(h)** Box plots compare average search time and prefix-match time across databases, reporting 10th-90th percentiles. **(i)** An example molecule (cisapride) is shown with its hierarchical fragmentation, highlighting selected terminal fragments indexed into the TRIE and excluded fragments.

**Supplementary Figure 2: Validation datasets, hit–miss definitions, and chemical heterogeneity**

**(a)** A pie chart illustrates the composition of the validation dataset comprising PROTAC*, DrugCentral, and other multitarget molecules. **(b)** A table reports dataset sizes, ligands, and target counts used for validation. **(c)** Bar plots show functional group distributions across validation datasets. **(d)** A schematic defines hit ratio and miss ratio metrics used for performance evaluation. **(e)** Density plots compare hit and miss ratios for PROTAC*, DrugCentral, and other datasets. **(f)** Density plots show LogP distributions across datasets. **(g)** t-SNE plots illustrate chemical heterogeneity of compounds based on molecular descriptors. **(h)** t-SNE clustering highlights separation of interaction types (types 1-4). **(i)** Bar plots compare ligand and target coverage across datasets. **(j)** Bar plots compare target recovery across prediction tools. **(k)** Venn diagrams show overlap of targets among validation datasets.

**Supplementary Figure 3: PROTAC generation and prioritization workflow**

**(a)** A schematic outlines the PROTAC design pipeline, including warhead selection, E3 ligase selection, linker generation, and optimization. **(b)** A funnel diagram illustrates PROTAC prioritization criteria across successive filtering stages. **(c)** A table lists generative model parameters used for linker design. **(d)** Atom count density plots for warheads, E3 ligands, and linkers are shown. **(e)** Line plots summarize E3 ligand physicochemical property distributions. **(f)** Scatter plots compare linker graph length and effective length across generated PROTACs.

**Figure 4: SynGlue PROTAC generation workflow, benchmarking, and model stability**

**(a)** The overall workflow of the linker-based multiclass classification model is illustrated, including feature extraction, selection, and training. **(b-c)** Bar plots show training and testing performance metrics for Signaturizer-based representations. **(d-e)** Training and testing performance for ChemBERTa-based molecular embeddings are shown. **(f-g)** Bar plots report ImageMol-based training and testing performance. **(h-i)** Training and testing performance for GROVER graph-based representations are shown. **(j-k)** Training and testing performance for Mordred descriptor-based models are shown. **(l-m)** Box plots summarize 10-fold cross-validation performance for the gradient boosting classifier. **(n)** A statistical table reports quartiles, whiskers, and sample size for each evaluation metric.

**Supplementary Figure 5: Attention-based regression modeling for PROTAC potency**

**(a-b)** Heatmaps compare data-driven and structure-guided similarity patterns between original and generated PROTACs. **(c)** A schematic illustrates the transformer-based regressor integrating component-wise attention over warhead, linker, and E3 ligase features. **(d)** Line plots show attention weight shifts across DC_50_ potency quantiles. **(e)** Box plots display distributions of attention weights assigned to PROTAC components. **(f)** A line plot shows training loss convergence over epochs. **(g-h)** Bar plots report regression performance metrics for D_max_ and DC_50_ prediction.

**Supplementary Figure 6: Multitarget molecule (MTM) design and diversity analysis**

**(a)** A schematic illustrates MTM generation using SynGlue’s data-driven pipeline. **(b)** Chemical structures of representative MTMs targeting EGFR, CDK1, PRKX, and DRD1 are shown. **(c)** Density plots show linker property distributions for MTMs. **(d)** Box plots assess physicochemical feasibility of MTMs. **(e)** A dot-matrix plot shows chemical class diversity and functional group enrichment across MTMs.

**Supplementary Figure 7: Benchmarking SynGlue against existing PROTAC tools**

**(a)** Bar plots compare predicted DC_50_ values across tools for selected compounds. **(b)** Bar plots show ranking accuracy in VCaP and LNCaP cell lines. **(c)** A comparative table summarizes tool functionality, methodology, training data, and outputs.

**Supplementary Figure 8: BRD4 warhead prioritization and binding analysis**

**(a)** A schematic outlines the BRD4 warhead prioritization workflow. **(b-g)** Residue proximity maps show atomic contacts between candidate warheads and BRD4 residues. **(h-l)** Interaction diagrams highlight key hydrogen bonds, π–π stacking, and hydrophobic interactions for prioritized warheads.

**Supplementary Figure 9: Docking, warhead diversity, and linker optimization**

**(a)** Density plots show Tanimoto similarity distributions of novel BRD4 warheads. **(b)** A network diagram illustrates warhead chemical diversity. **(c)** A schematic describes the docking grid setup and parameters. **(d)** Bar plots show docking binding energies for candidate warheads. **(e-g)** Density plots depict linker length, effective length, and ring count distributions. **(h-i)** Density plots show predicted Dmax and DC_50_ distributions for selected PROTACs.

**Supplementary Figure 10: Physicochemical filtering and cavity analysis**

**(a)** Bar plots show molecule counts retained after sequential physicochemical filters. **(b)** Box plots show maximum Tanimoto similarity to known PROTACs. **(c)** Density plots illustrate similarity distributions for designed PROTACs. **(d)** Structural visualizations depict BRD4 cavity druggability metrics. **(e-h)** Tables summarize interface residues, cavity properties, and chain interactions.

**Supplementary Figure 11: Linker optimization and final PROTAC selection**

**(a)** Density plots show optimized linker physicochemical properties. **(b)** Bar plots depict molecule retention after filtering stages. **(c)** Density plots show predicted DC_50_ and D_max_ distributions for selected GSPT1 PROTACs.

**Supplementary Figure 12: Synthesis, Characterization, and Functional Evaluation of SynGlue-designed PROTACs for BRD4.**

**(a)** High-performance liquid chromatography (HPLC) chromatogram of BRD4 (cmp1), showing a primary peak at 6.450 min retention time (RT) with an area percentage of 98.37%, confirming its purity. **(b)** High-resolution mass spectrometry (HRMS) data for BRD4 (cmp1), including the extracted ion chromatogram and the observed molecular ion peaks, confirming its exact mass (1020.4415 Da) and molecular formula (C_52_H_64_N_8_O_11_S). **(c)** ^1^H NMR (top) and ^13^C NMR (bottom) spectra of BRD4 (cmp1) in DMSO-d_6_, demonstrating expected chemical shifts and confirming the compound’s structural features. **(d)** HPLC chromatogram of BRD4 (cmp 2), with a primary peak at 7.455 min RT and an area percentage of 98.26%, indicative of high compound purity. **(e)** HRMS data for BRD4 (cmp2) displaying extracted ion chromatograms and molecular ion peaks consistent with its exact mass (1016.4830 Da) and molecular formula (C_53_H_66_N_8_O_10_S). **(f)** ^1^H NMR (top) and ^13^C NMR (bottom) spectra of BRD4 (cmp2) in DMSO-d_6_, demonstrating expected chemical shifts and confirming the compound’s structural features. **(g)** High-performance liquid chromatography (HPLC) chromatogram of BRD4 (cmp1) inactive isomer, showing a prominent peak at 6.357 minutes retention time (RT) with an area percentage of 97.69%, confirming its purity. **(h)** High-resolution mass spectrometry (HRMS) spectrum of the inactive isomer of BRD4 (cmp1), displaying the observed molecular ion peaks and confirming the exact mass (1020.4415 Da) and the molecular formula (C_53_H_64_N_8_O_11_S) of the compound. **(i)** Nuclear magnetic resonance (NMR) spectra of BRD4 (cmp2) in DMSO-d_6_: the upper panel shows the ^1^H NMR spectrum, with characteristic proton signals across the chemical shift range, while the lower panel presents the ^13^C NMR spectrum, depicting distinct carbon chemical shifts, together confirming the structural integrity and chemical identity of the BRD4 (cmp1) inactive isomer.

**Supplementary Figure 13: In Vivo Pharmacokinetics, Tolerability, and Pharmacodynamics of SynGlue-designed BRD4 PROTAC.**

**(a, b)** Dose-response curves for cell proliferation inhibition by BRD4 (cmp1) and its inactive isomer in MV-4-11 cell lines, showing differential potency across varying concentrations (nM range). **(c, d)** Western blot analysis of BRD4 degradation in MV-4-11 cells treated with increasing concentrations (0.001-1000 nM) of BRD4 (cmp1) and BRD4 (cmp2) for 16 hours, with tubulin as a loading control. The lower panels show dose-response curves for BRD4 degradation in MV-4-11 cells, with DC_50_ values calculated as 0.193 μM for BRD4 (cmp1) and 0.228 μM for BRD4 (cmp2).

**Supplementary Figure 14: Analytical characterization of GSPT1 PROTACs**

**(a)** High-performance liquid chromatography (HPLC) chromatogram of GSPT1 (cmp1), showing a primary peak at 6.797 min retention time (RT) with an area percentage of 98.70%, confirming its purity. **(b)** Liquid chromatography-mass spectrometry (LC-MS) data for GSPT1 (cmp1), including the extracted ion chromatogram and the observed molecular ion peaks, confirming its exact mass (647.23 Da) and molecular formula (C35H33N7O4S). **(c)** ¹H NMR spectra of GSPT1 (cmp1) in DMSO-d_6_, demonstrating expected chemical shifts and confirming the compound’s structural features. **(d)** HPLC chromatogram of GSPT1 (cmp2), with a primary peak at 6.317 min RT and an area percentage of 98.89%, indicative of high compound purity. **(e)** Liquid chromatography-mass spectrometry (LC-MS) data for GSPT1 (cmp2), including the extracted ion chromatogram and the observed molecular ion peaks, confirming its exact mass (675.26 Da) and molecular formula (C37H37N7O4S). **(f)** H NMR spectra of GSPT1 (cmp2) in DMSO-d_6_, demonstrating expected chemical shifts and confirming the compound’s structural features.

## Supplementary Data

**Supplementary Table 1:** Tabular representation detailing information about the statistical summary of the curated MagnetDB database, including counts of unique protein-ligand interactions, compounds, protein targets, and terminal fragments, broken down by species and originating source database.

**Supplementary Table 2:** Tabular representation detailing information about the core modules comprising the SynGlue platform and the key external software packages, libraries, and databases utilized or integrated within the workflow, along with their version numbers where applicable.

**Supplementary Table 3:** Tabular representation of information about the composition of the benchmarking dataset: CompoundID, their SMILES, and class.

**Supplementary Table 4:** Tabular representation detailing information about the calculated hit ratio and miss ratio values for SynGlue and the benchmark tools (LigAdvisor, SwissTargetPrediction, and SuperPred) based on target predictions for the selected subset of 100 representative molecules, allowing direct comparison.

**Supplementary Table 5:** Tabular representation detailing information about key characteristics of Androgen Receptor (AR)-XIAP PROTACs designed *de novo* using both the data-driven and structure-guided workflows of SynGlue, potentially including compound identifiers, generating workflow, predicted properties, scores, and comparison data for experimentally validated reference AR PROTACs.

**Supplementary Table 6:** Tabular representation detailing information about key characteristics of Multi-Target Molecules (MTMs) designed *de novo* using SynGlue to target specific pairs (EGFR/CDK1 and PRKX/DRD1), potentially including compound identifiers, targeted protein pair, generating workflow, and predicted properties.

**Supplementary Table 7:** Tabular representation detailing information about key characteristics of BRD4-VHL PROTACs designed *de novo* using both the data-driven and structure-guided workflows of SynGlue, potentially including compound identifiers, generating workflow, predicted properties, scores from predictive models, and comparison data for experimentally validated reference BRD4 PROTACs.

**Supplementary Table 8:** Tabular representation detailing information about the *in silico* characterization of the final prioritized BRD4-VHL PROTAC candidates selected for experimental validation, including compound identifiers, key computed physicochemical properties, predicted degradation potency (DC_50_, D_max_ values), and predicted potency class.

**Supplementary Table 9:** Plasma and tumor concentration profiles of BRD4 (cmp1) in MV-4-11 tumor-bearing mice. The table summarizes compound exposure measured at defined time points following dosing during the efficacy study, including data from the 14-day maximum tolerated dose (MTD) assessment. Reported values capture systemic and intratumoral drug levels observed under repeated dosing to evaluate exposure, tolerability, and pharmacokinetic-pharmacodynamic (PK/PD) relationships. The table also includes a summary of the PK/PD study design, detailing the dosing regimen, sampling schedule, analytical endpoints, and exposure–response metrics derived from single-dose and repeated-dose analyses in plasma and tumor lysates.

**Supplementary Table 10:** List of designed GSPT1-targeting molecules generated during the study, including compound identifiers and structural information used for subsequent prioritization.

**Supplementary Table 11:** Tabular summary of in silico characterization of the final prioritized GSPT1-CRBN PROTAC candidates, including computed physicochemical properties, predicted degradation potency (DC_50_ and D_max_), and assigned potency classes.

**Supplementary Table 12:** Detailed information on cell lines used in GSPT1 studies. The table summarizes cell type, tissue of origin, and experimental context for each model, along with optimized experimental conditions including seeding density (cells per well), culture medium, and treatment duration. It also reports dose-response data across multiple cell lines, presenting log-transformed compound concentrations (log µM), individual biological replicate responses (response1-response3), and corresponding mean response values.

**Supplementary Table 13:** Differential protein expression analysis identifying significantly regulated proteins, including UniProt identifiers, gene names, log fold change (logFC), average expression (AveExpr), statistical test values, raw and adjusted P values, and log-odds of differential expression.

