## supplementary Figure 1 for "Generative AI Framework SynGlue for the Rational Design of Clinically relevant Protein Degraders"

a

| Database | Version | Unique Compound | Fragment | Terminal Fragment | Targets |
|---|---|---|---|---|---|
| DrugBank | 5.1.10 | 5,347 | 9,576 | 18,312 | 798 |
| BindingDB | 28-05-23 | 670,995 | 321,421 | 261,340 | 3,060 |
| ChEMBL | ChEMBL33 | 487,964 | 933,792 | 307,929 | 2,055 |
| STITCH | 5.0 | 646,590 | 3,070,431 | 1,645,549 | 4,289 |
| Small Molecule Suite | 2019 | 328,709 | 244,121 | 210,421 | 3,094 |
| BioSNAP ** | 2018 | 165,148 | 144,110 | 103,212 | 6,833 |
| Total | | | | | 20,129 |

**- ChG-Miner, ChG-TargetDecagon, ChG-Interdecagon

b

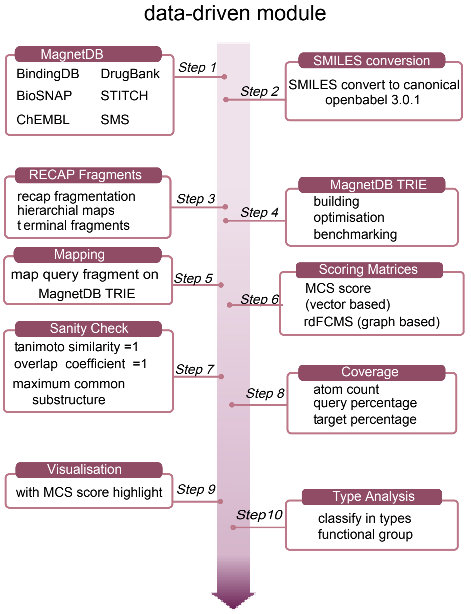


c

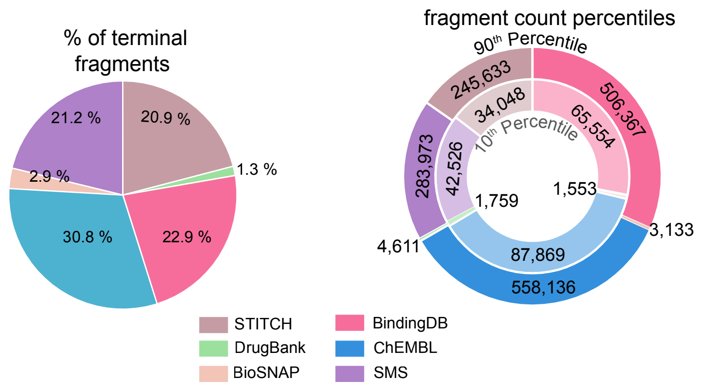


d

| parameter | count |
|---|---|
| branches | 2,471,351 |
| nodes | 5,727,812 |
| avg build time(sec) | 252 |
| depth | 1,568 |
| avg frequecy | 0.0406 |
| max branching factor | 8,492 |
| efficiency | 99.98 |
| master node | $ |

e

| Operation | Time Complexity | Auxillary Space |
|---|---|---|
| Insert | O(n) | O(n*m) |
| Search | O(n) | O(1) |
| Delete | O(n) | |

n - number of words  m- max length of word

f

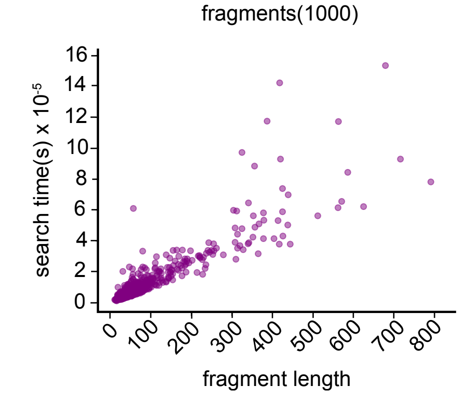


g

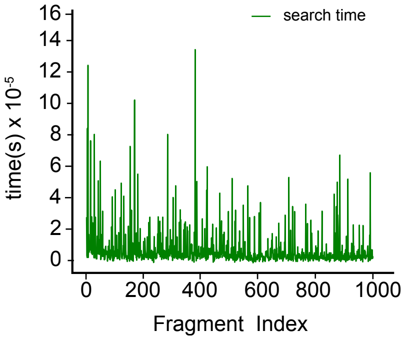


h

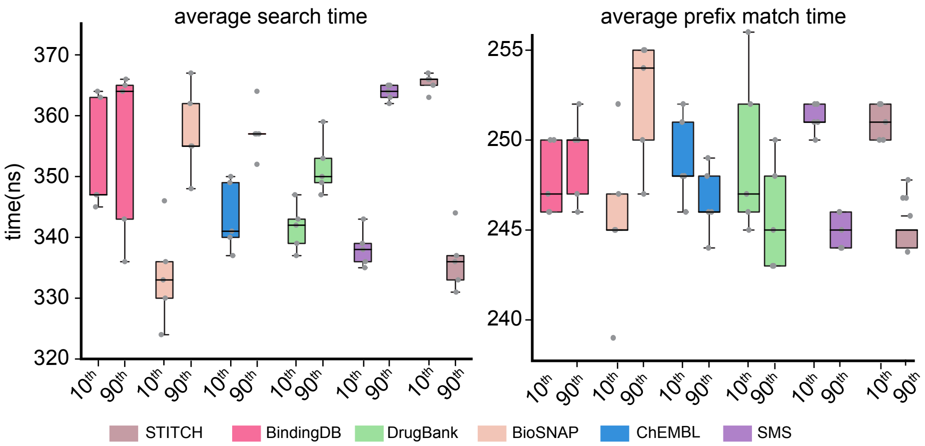


i

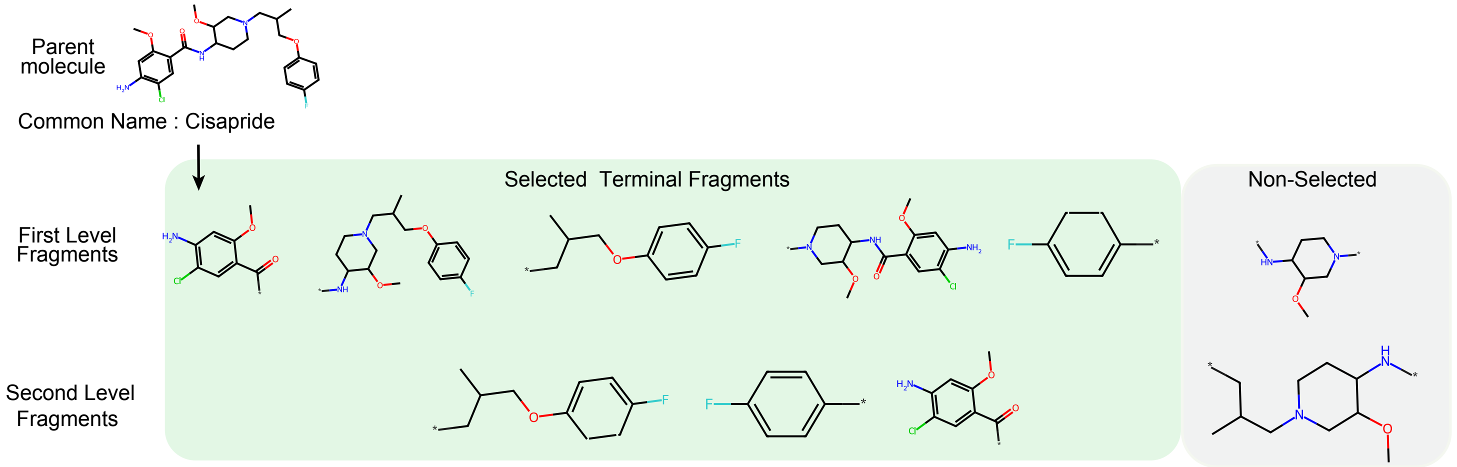


**Supplementary Figure 1**
