## supplementary Figure 2 for "Generative AI Framework SynGlue for the Rational Design of Clinically relevant Protein Degraders"

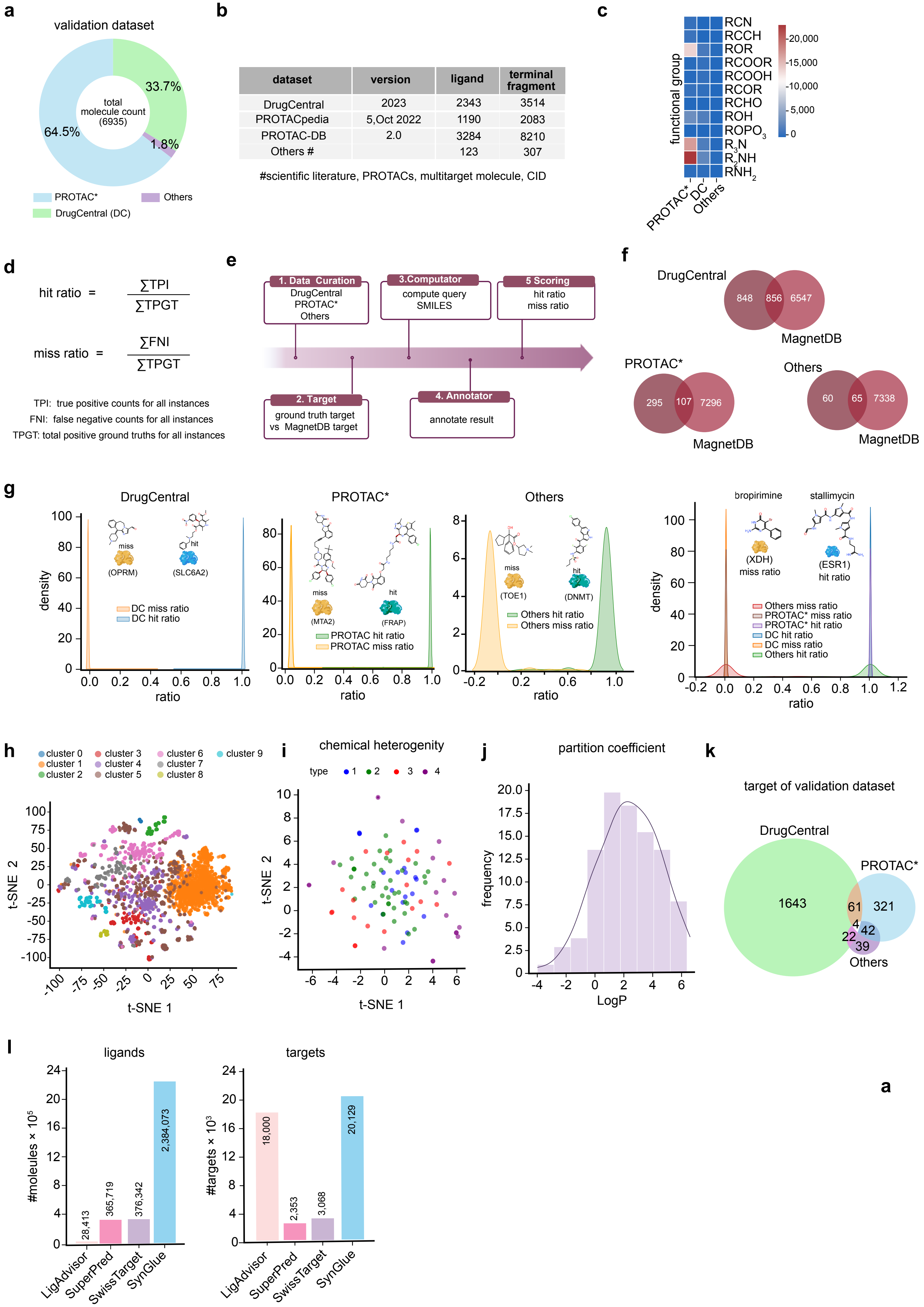


| dataset | version | ligand | terminal fragment |
|---|---|---|---|
| DrugCentral | 2023 | 2343 | 3514 |
| PROTACpedia | 5,Oct 2022 | 1190 | 2083 |
| PROTAC-DB | 2.0 | 3284 | 8210 |
| Others # | | 123 | 307 |

**Supplementary Figure 2**
