## Supplementary figures and images for "Generative AI Framework SynGlue for the Rational Design of Clinically relevant Protein Degraders"

### supplementary Figure 3

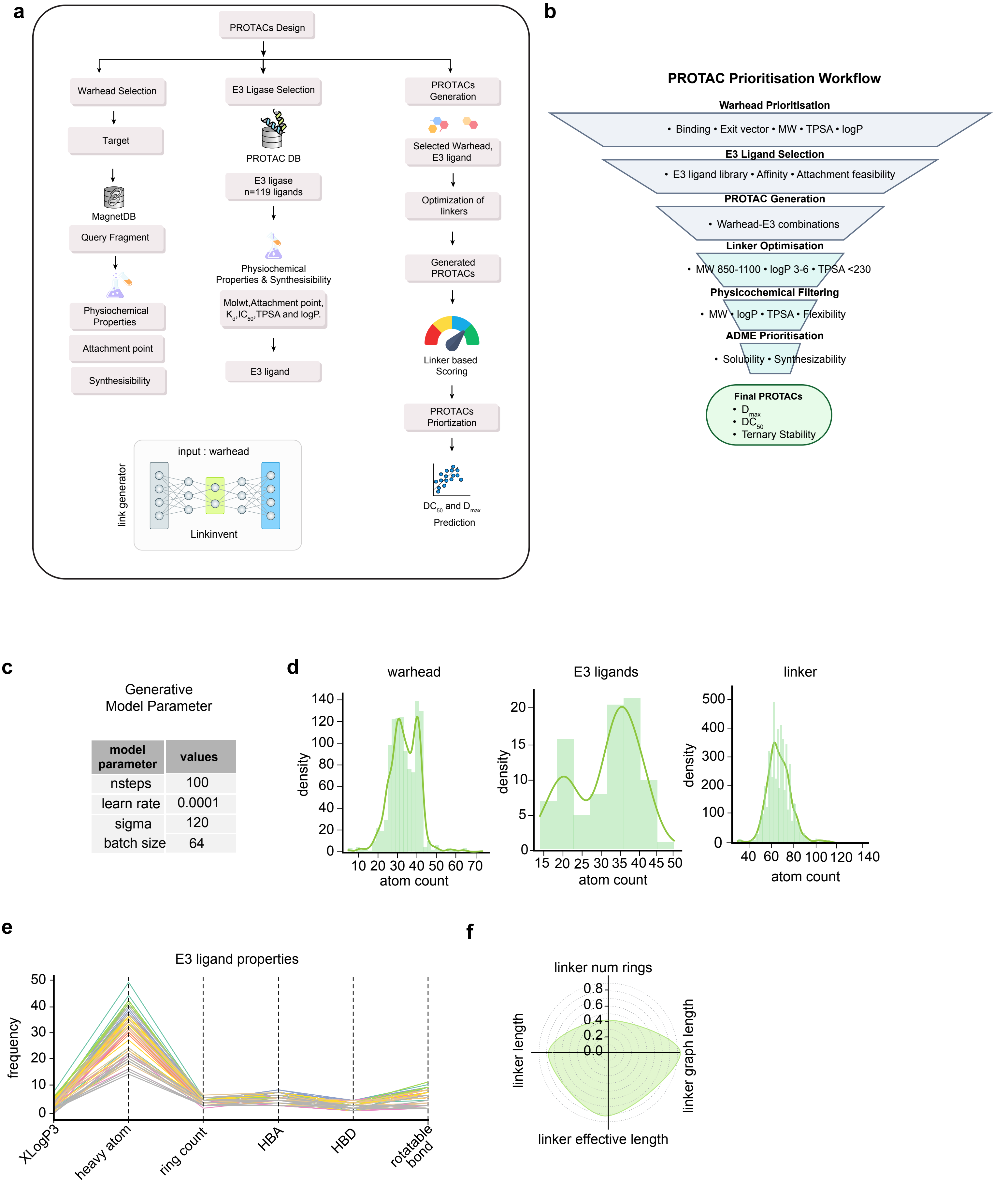


Supplementary Figure 3

### supplementary Figure 5

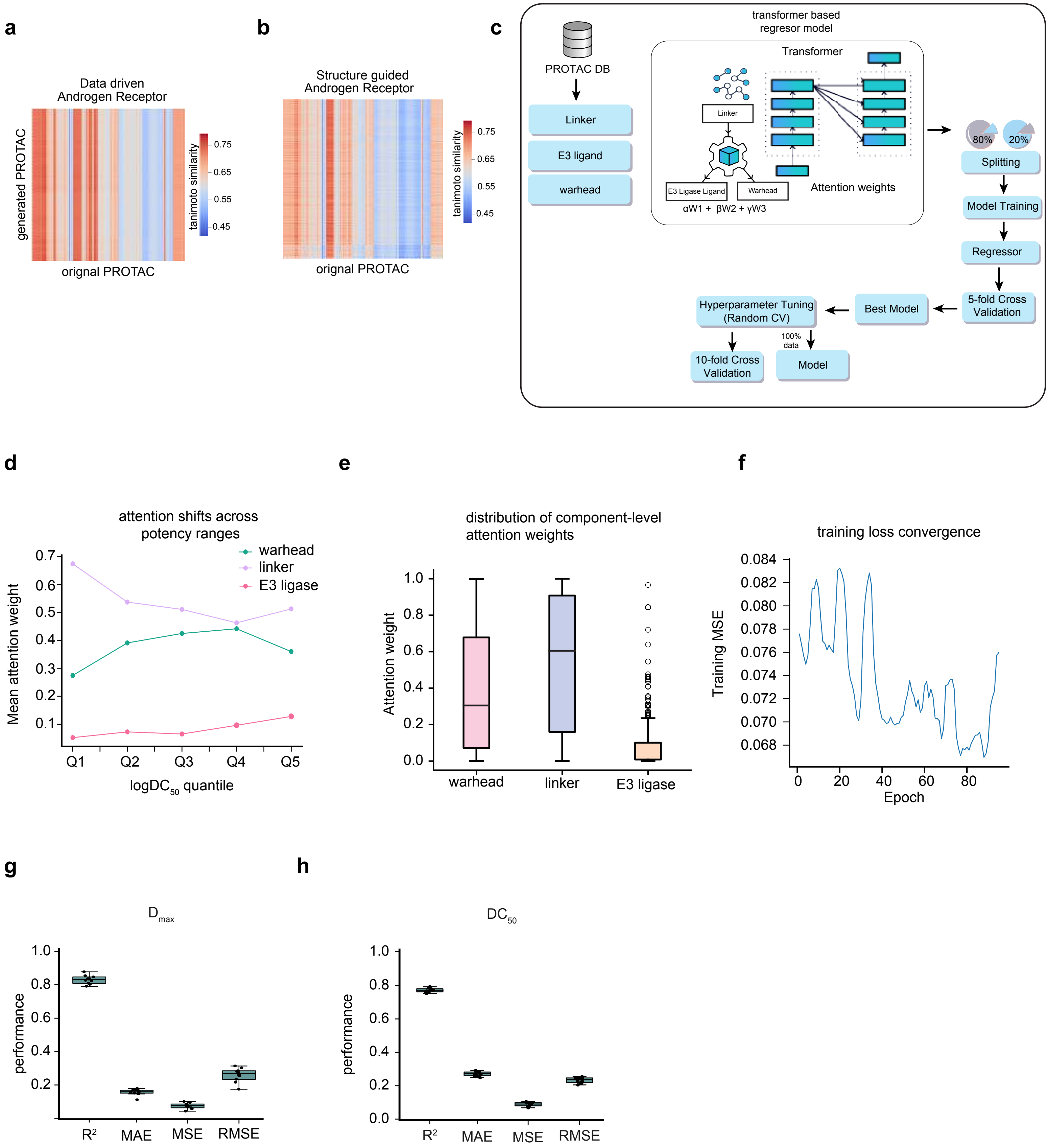


**Supplementary Figure 5**

### supplementary Figure 6

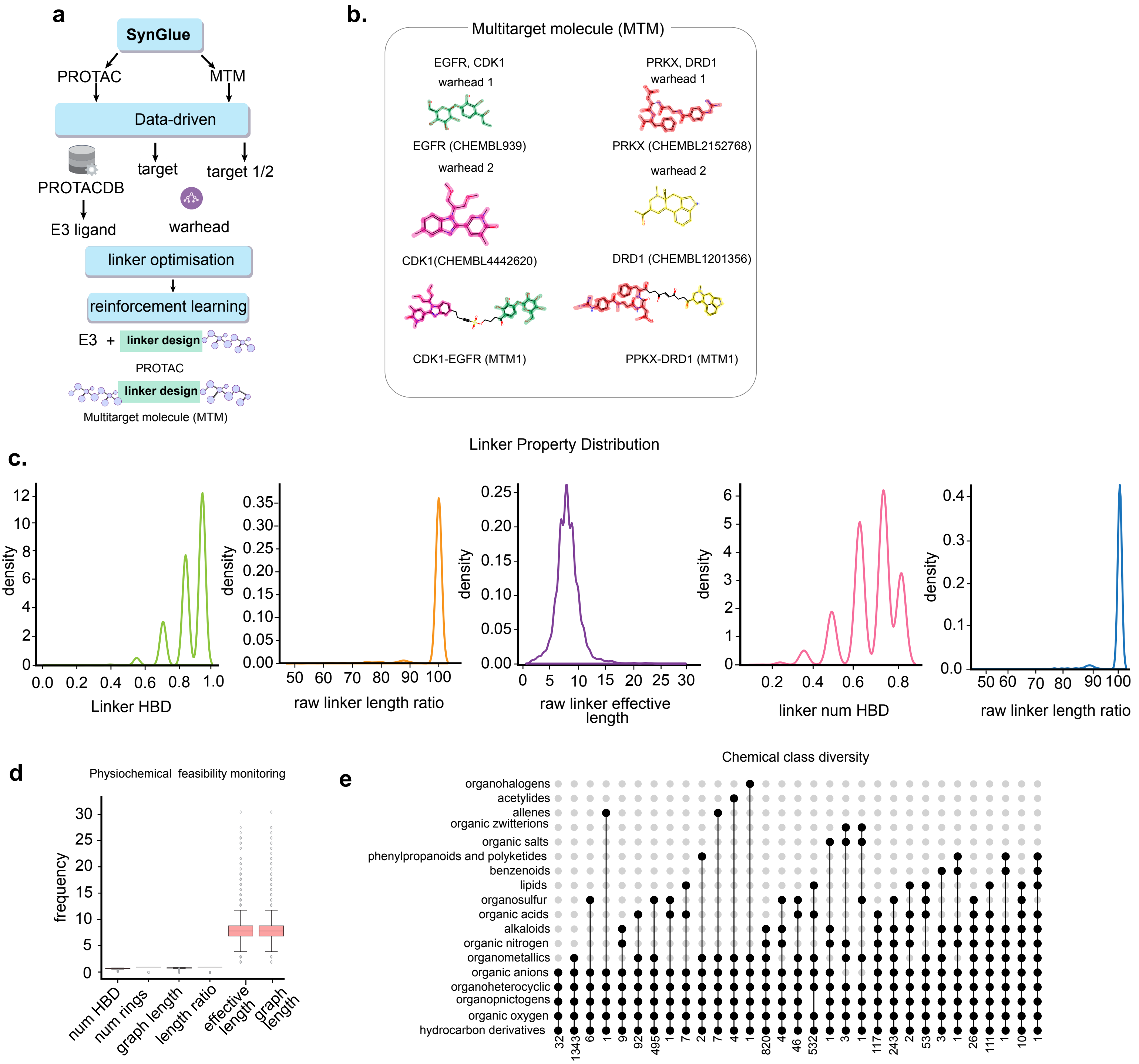


**Supplementary Figure 6**

### supplementary Figure 8

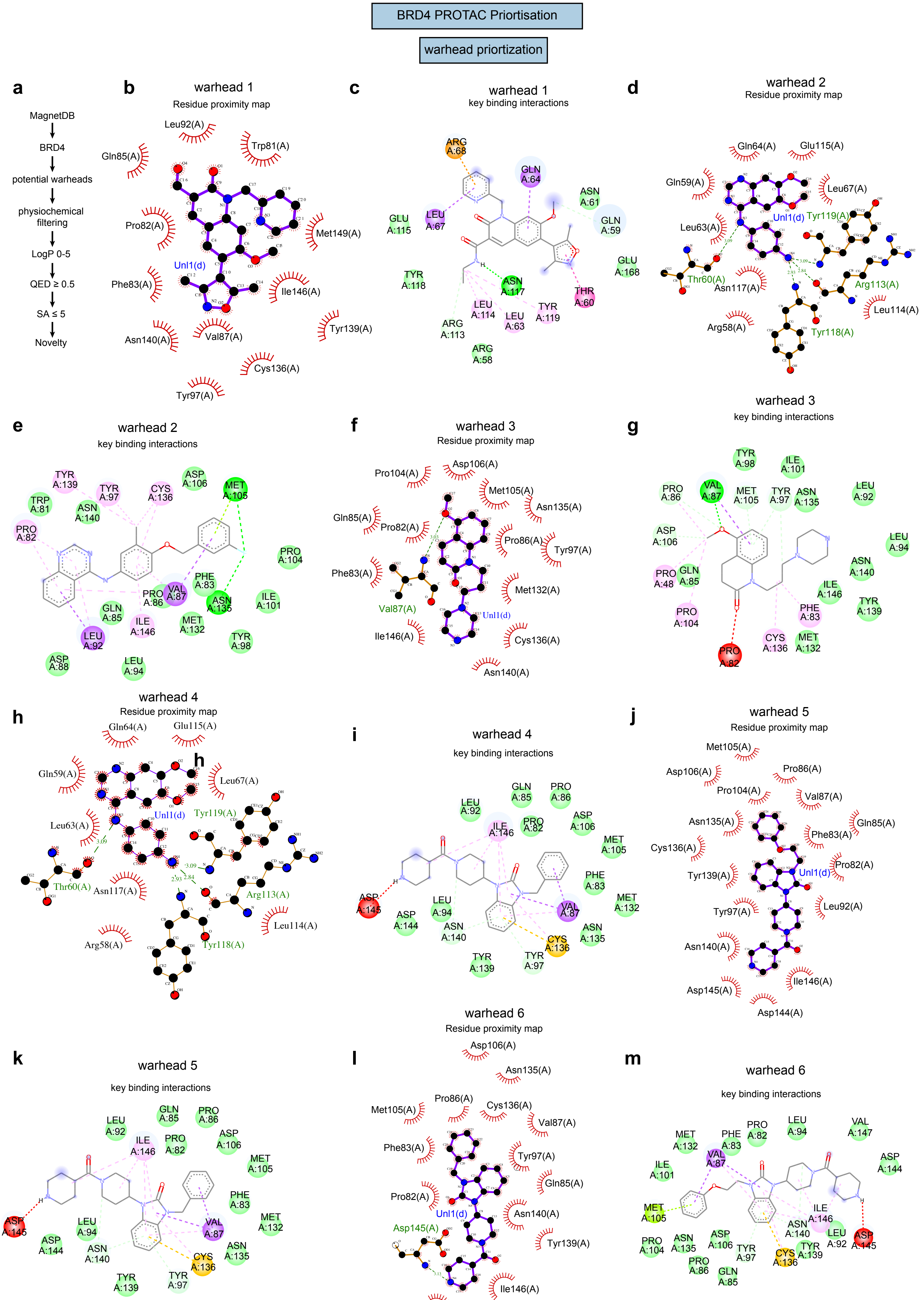


Supplementary Figure 8

### supplementary Figure 9

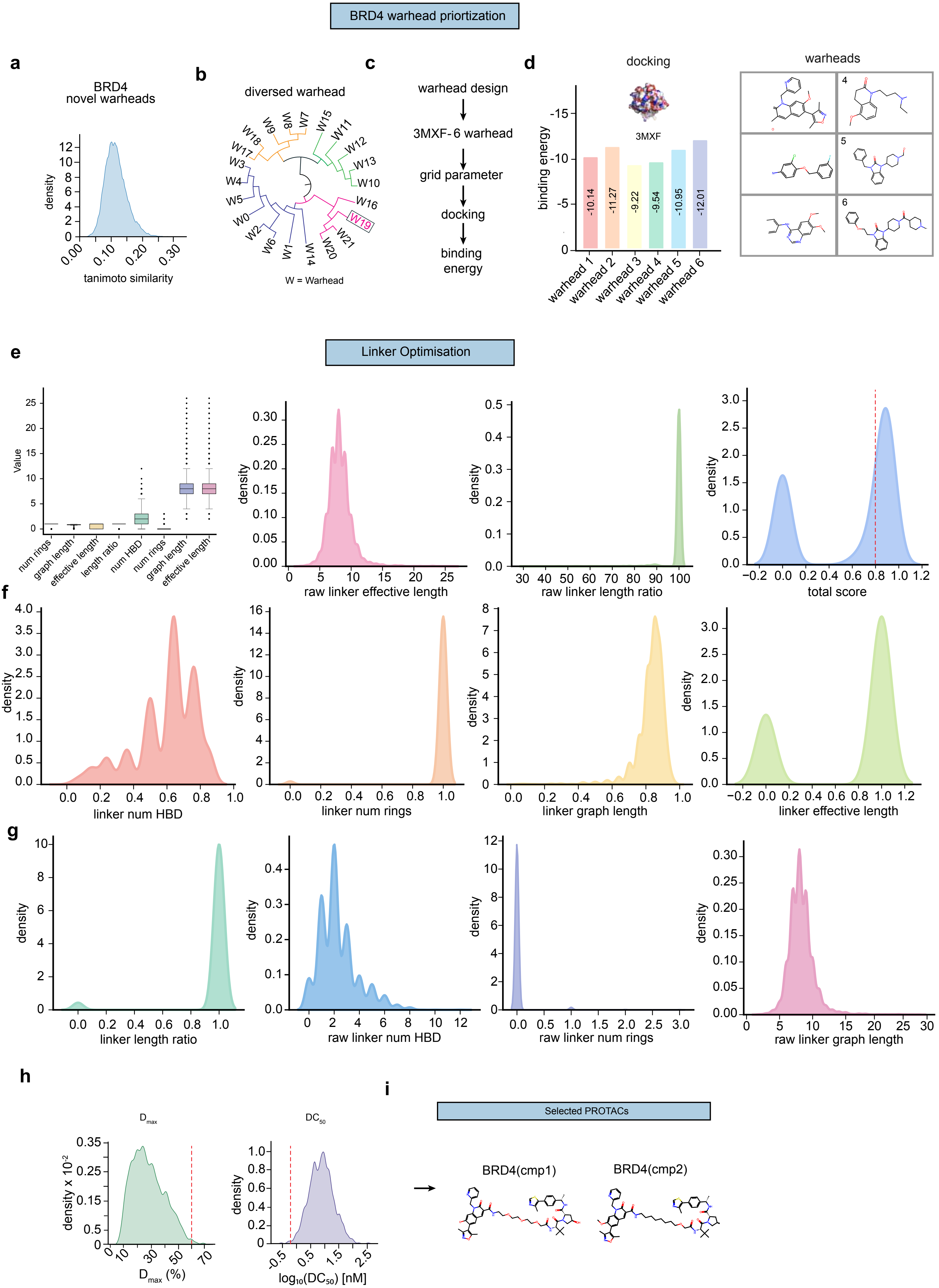


**Supplementary Figure 9**

### supplementary Figure 11

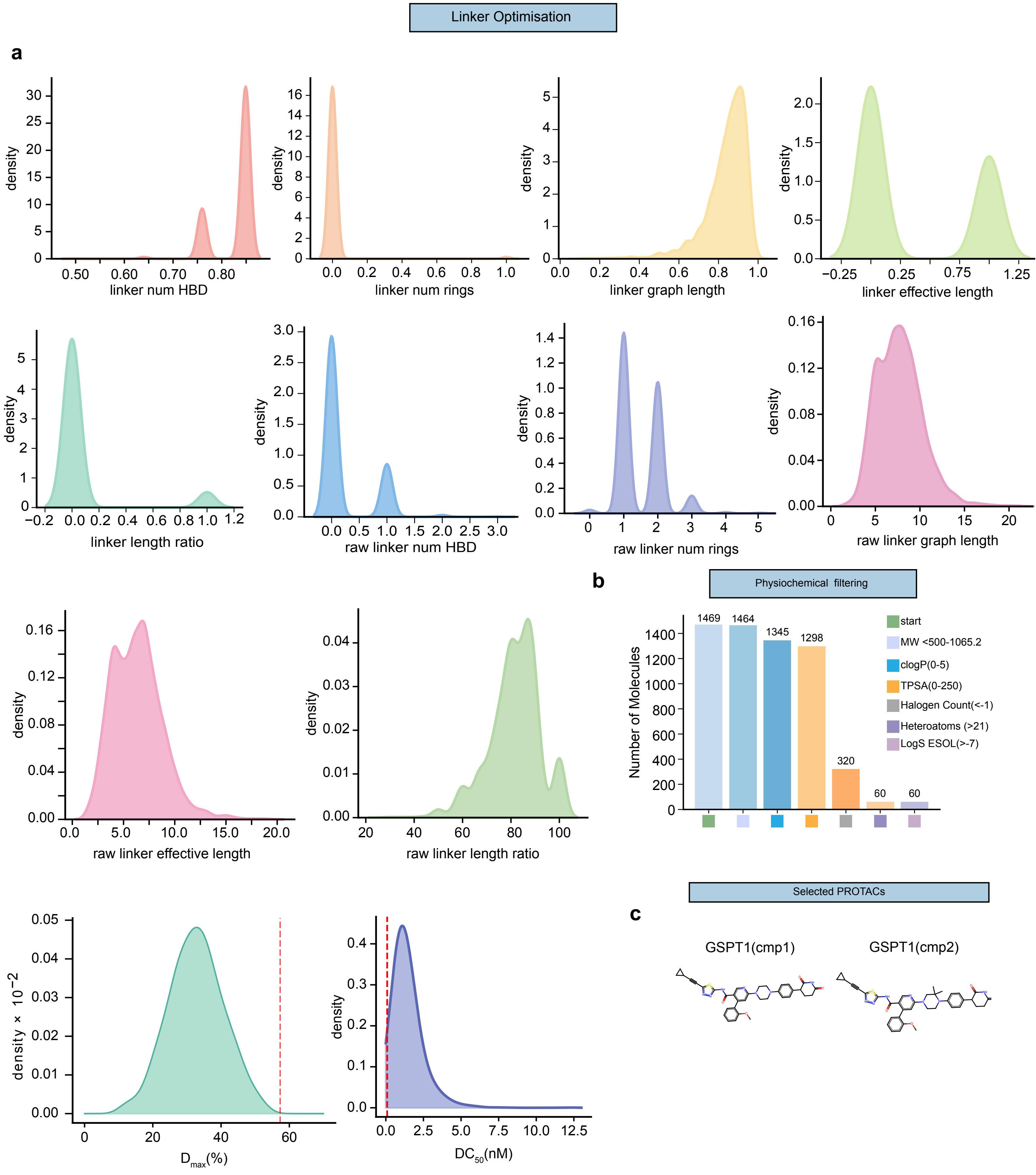


**Supplementary Figure 11**
