## supplementary Figure 4 for "Generative AI Framework SynGlue for the Rational Design of Clinically relevant Protein Degraders"

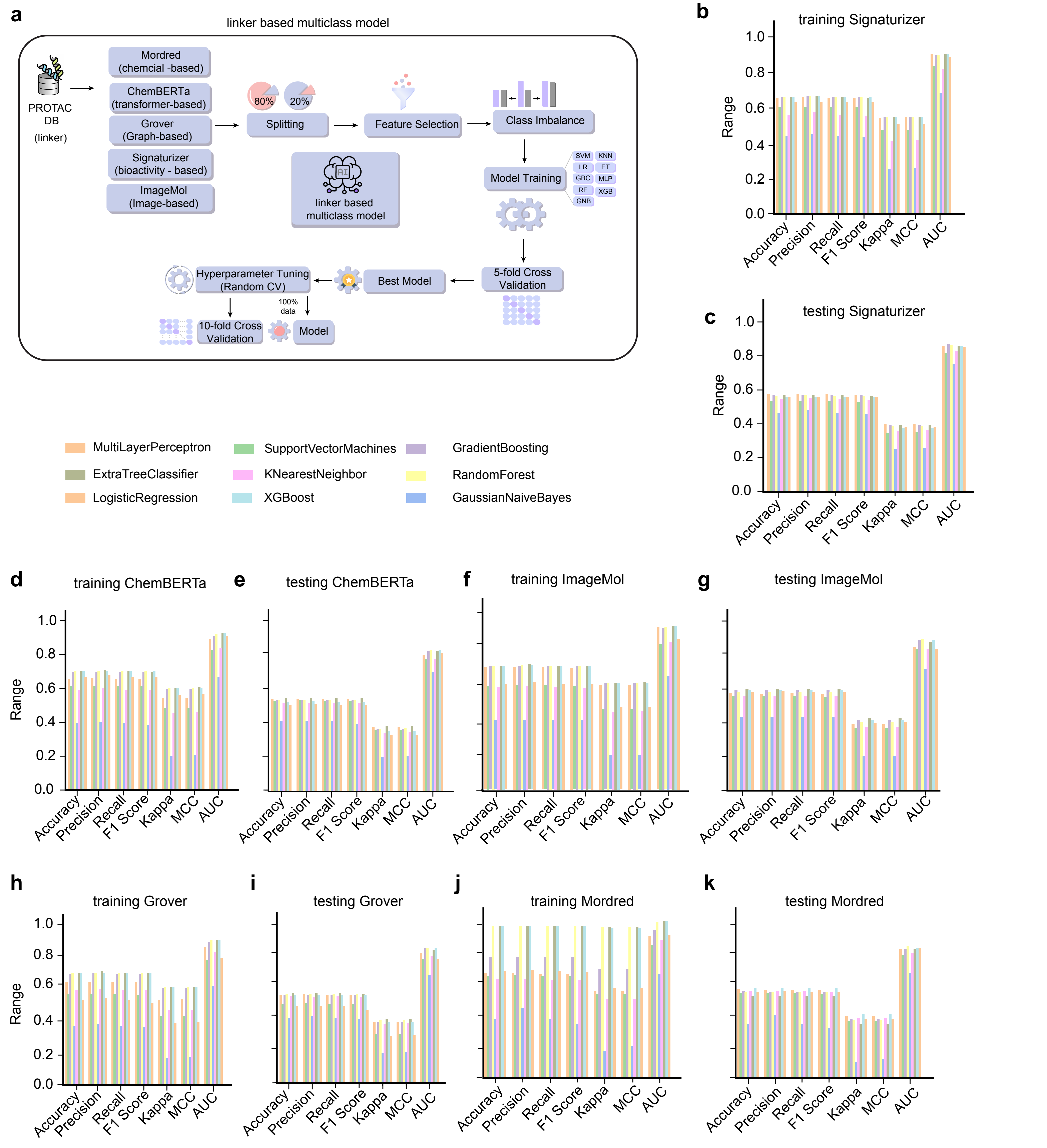


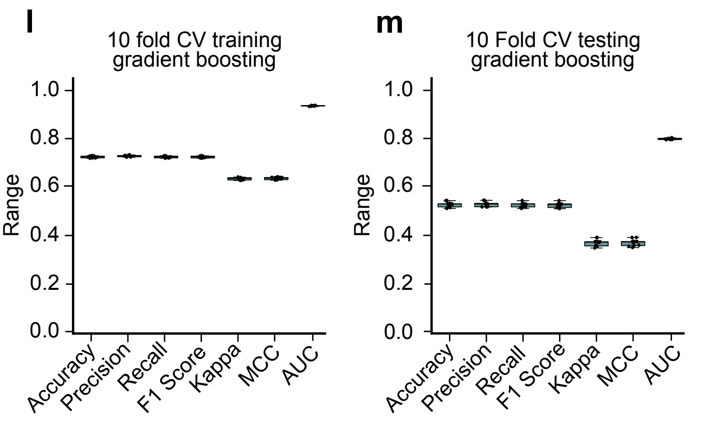


n

| | Test Accuracy | Test precision | Test Recall | Test F1 Score | Test Kappa | Test MCC | Test AUC |
|---|---|---|---|---|---|---|---|
| Upper whisker | 0.54 | 0.54 | 0.54 | 0.54 | 0.39 | 0.39 | 0.80 |
| 3rd Quartile | 0.53 | 0.53 | 0.53 | 0.53 | 0.37 | 0.37 | 0.80 |
| Median | 0.53 | 0.53 | 0.53 | 0.53 | 0.37 | 0.37 | 0.80 |
| 1st Quartile | 0.52 | 0.52 | 0.51 | 0.51 | 0.35 | 0.35 | 0.79 |
| Lower Whisker | 0.51 | 0.51 | 0.51 | 0.51 | 0.34 | 0.35 | 0.79 |
| no of data point | 10 | 10 | 10 | 10 | 10 | 10 | 10 |

**Supplementary Figure 4**
