## supplementary Figure 7 for "Generative AI Framework SynGlue for the Rational Design of Clinically relevant Protein Degraders"

**a**

| Evaluation Criteria | SynGlue | PDP Tool |
|---|---|---|
| Potency Prediction | Quantitative($DC_{50}$ , $D_{max}$) | Normalised (0-1) |
| Predicts $D_{max}$ | Yes | No |
| Predicts $DC_{50}$ | Yes(nM) | No |
| VCaP ranking accuracy | Correct (C1>C2) | Wrong (C1>C2) |
| LNCaP ranking accuracy | Correct | Wrong |
| Correlation to western blot | Strong (C1>C2) | Weak (C1>C2) |
| Interpretability | High | Medium |

**b**

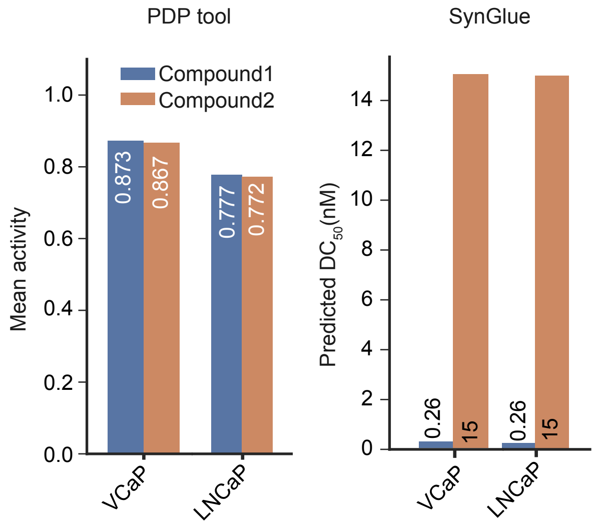


**c**

| Tool Name | Primary Function | Core Methodology | Primary Database/ Training Data | Component Sourcing | Potency Assessment (Method) | Output |
|---|---|---|---|---|---|---|
| SynGlue | De novo Design + Prediction | Data Driven , Structure guided supported by transformer Model | Magnet DB + PROTAC-DB(3.0)+ Protecpedia + curated degraders | W - MagnetDB<br>L - DeNovo<br>E3 - PROTACDB | Transformer based Regression, linker length based multiclass | PROTAC candidate ranked by $DC_{50}$ , $D_{max}$ |
| DiffPROTACs | De novo linker Design | Transformer model | ZINC + GEOM + PROTAC-DB(2.0) | NA | Generative tool | 3D linker structure |
| PROTACable | De novo Design | Reinforcemnet | PROTAC-DB(2.0) | NA | Boosted tree | PROTACs |
| DeepPROTACs | Predict degradation efficiency | GNN +CNN hybrid | PROTAC-DB + curated degraders | NA | Deep Neural Scoring model | predicted $DC_{50}$ , $D_{ma\ x,}$ |
| PRODE | PROTAC scoring & property evaluation | Ensemble ML Model | PROTAC-DB + PRODE internal dataset | NA | Ensemble potency predictor | Degradation score, ADMET |
| PRoSettaC | Structure Modeling of ternary complexes | Rosetta-based docking + conformational sampling | PDB + modeled structures | E3+POI crystal structure | Rosetta energy scoring | Ternary complex poses + G predictions |
| Schrodinger Toolkit | Structure based PROTAC design | Molecular docking, MD,FEP+,QM Methods | PDB + proprietary DB | Ligand/E3 warhead libraries | FEP +potency prediction | Ternary complex model G predictions |

**Supplementary Figure 7**
