## supplementary Figure 10 for "Generative AI Framework SynGlue for the Rational Design of Clinically relevant Protein Degraders"

### Physiochemical filtering

a

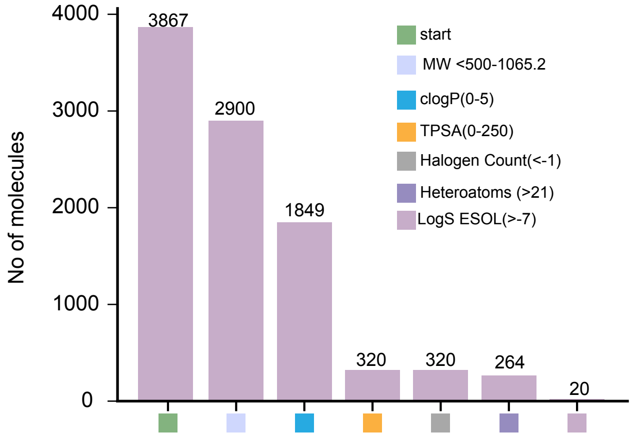


b

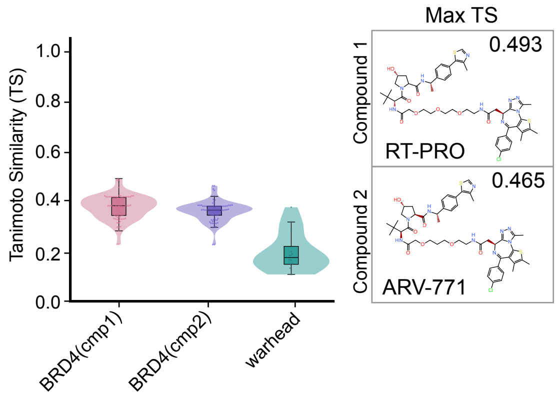


c

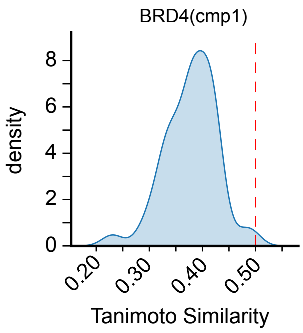


d

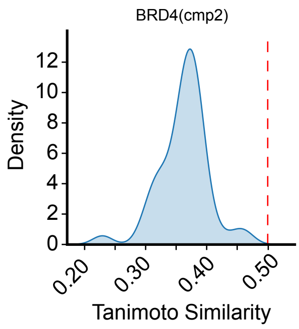


e

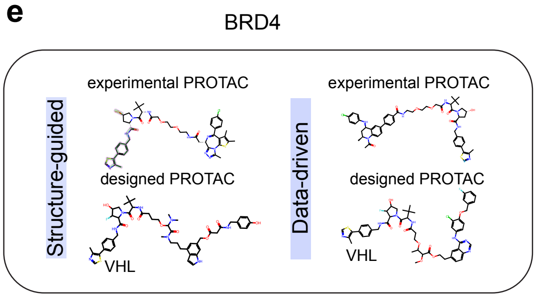


BRD4 structure guided cavity

| Cavity | Volume | Average pKd | Drug Score | Druggability |
|---|---|---|---|---|
| 1 | 384.5 | 6.87 | 1926 | Druggable |
| 2 | 384.5 | 6.83 | -743 | Undruggable |
| 3 | 384.5 | 6.38 | -812 | Undruggable |
| 4 | 384.5 | 4.98 | -1541 | Undruggable |
| 5 | 384.5 | 4.81 | -1614 | Undruggable |

### GSPT1 PROTAC Priortisation

#### warhead priortization

e

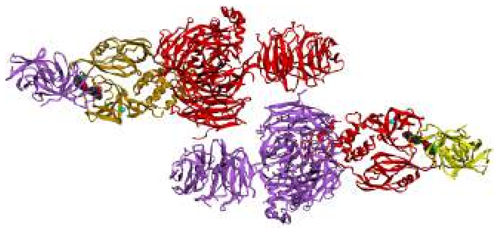

PDB ID - 5HXB
Resolution -3.60A

f

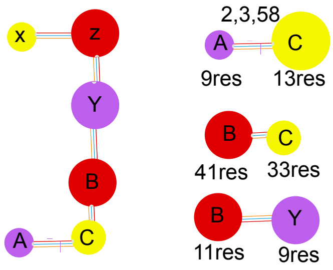


g

| chains | no of interface residue | interface area(A²) | No of salt bridge | No of H bonds | No of non bonding contacts |
|---|---|---|---|---|---|
| A-C | 9:13 | 618:617 | 2 | 3 | 0.39 |
| X-Z | 9:13 | 653:656 | 2 | 5 | 0.37 |
| B-C | 41:33 | 2042:2223 | | 6 | 0.37 |
| Y-Z | 41:36 | 2107:2272 | | 8 | 0.35 |
| B-Y | 11:9 | 445:498 | | 2 | 0.34 |

equivalent chains X =A ,Y=B ,Z=C

h

| Entity ID | Chain ID | Description | Type | Chain Length |
|---|---|---|---|---|
| 1 | A,X | eRF3a,G1 to S phase transition protein 1 homolog | polymer | 0.39 |
| 2 | B,Y | DNA damage-binding protein 1 | polymer | 0.37 |
| 3 | C,Z | Protein cereblon | polymer | 0.37 |
| 4 | G,I | Zinc ion | non polymer | 0.35 |

i

Molecule1 GSPT1

| Mol | Chain | Atoms | | | | | Residues |
|---|---|---|---|---|---|---|---|
| 1 | X | Total 1481 | C 941 | N 254 | O 275 | S 11 | 195 |
| 1 | A | Total 1441 | C 919 | N 245 | O 267 | S 10 | 196 |

**Supplementary Figure 10**
