## supplementary Figure 12 for "Generative AI Framework SynGlue for the Rational Design of Clinically relevant Protein Degraders"

**a**

HPLC of BRD4(cmp1)

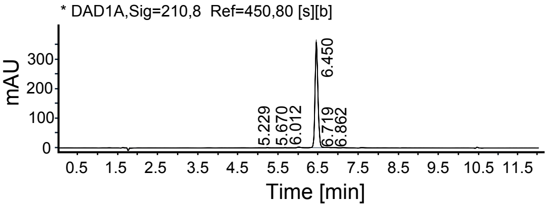


| RT[min] | Width[min] | Area | Area % |
|---|---|---|---|
| 5.229 | 0.14 | 1.90 | 0.10 |
| 5.670 | 0.11 | 0.90 | 0.05 |
| 6.012 | 0.24 | 13.38 | 0.73 |
| 6.450 | 0.39 | 1799.56 | 98.37 |
| 6.719 | 0.16 | 9.54 | 0.52 |
| 6.862 | 0.20 | 4.17 | 0.23 |

**b**

HRMS of BRD4(cmp1)

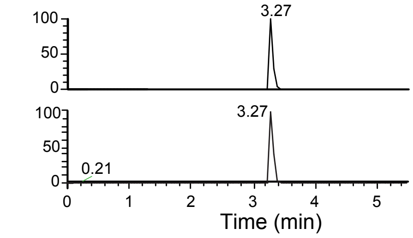


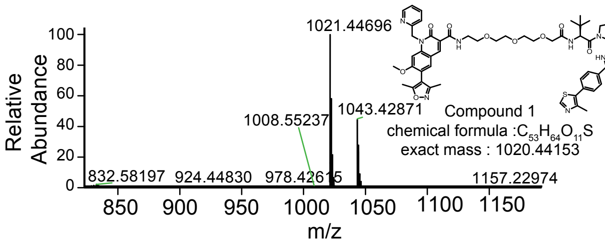


**c**

[1]H NMR of BRD4(cmp1)

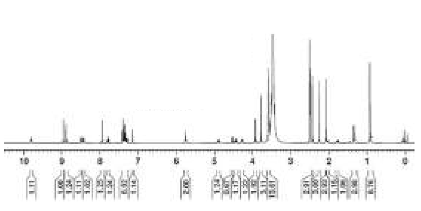

[13]C NMR of BRD4(cmp1)

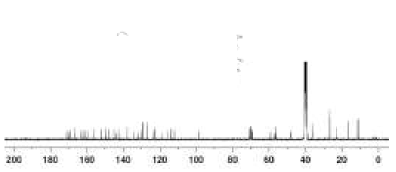

**d**

HPLC of BRD4(cmp2)

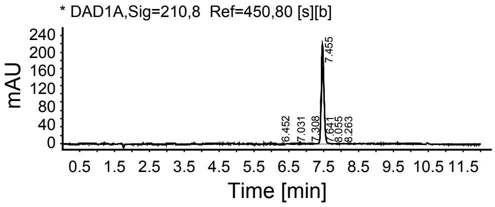


| RT[min] | Width[min] | Area | Area % |
|---|---|---|---|
| 6.452 | 0.17 | 2.67 | 0.22 |
| 7.031 | 0.13 | 0.92 | 0.07 |
| 7.308 | 0.12 | 2.37 | 0.18 |
| 7.455 | 0.32 | 1278.51 | 98.26 |
| 7.641 | 0.24 | 9.81 | 0.75 |
| 8.055 | 0.32 | 5.50 | 0.42 |
| 8.263 | 0.14 | 1.23 | 0.09 |

**e**

HRMS of BRD4(cmp2)

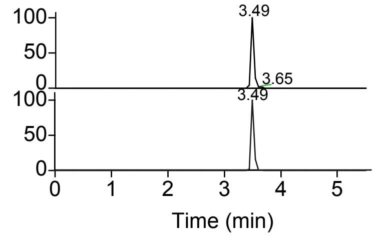


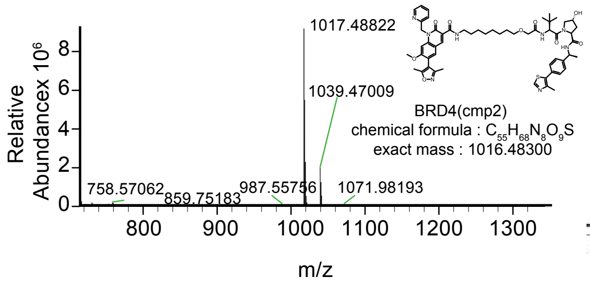


**f**

[1]H NMR of BRD4(cmp2)

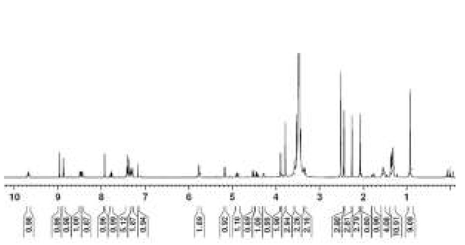

[13]C NMR of BRD4(cmp2)

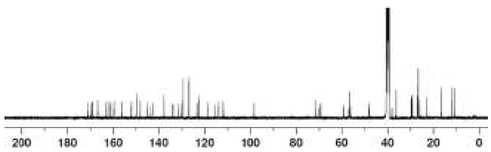

**g**

HPLC of Compound 1 Inactive Isomer

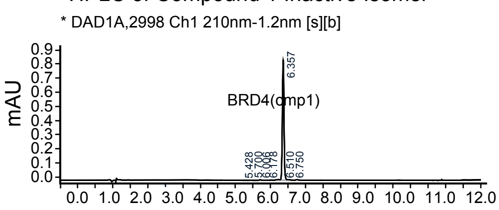


| RT[min] | Width[min] | Area | Area % |
|---|---|---|---|
| 5.428 | 0.14 | 0.00 | 0.15 |
| 5.700 | 0.14 | 0.01 | 0.42 |
| 6.006 | 0.10 | 0.00 | 0.07 |
| 6.178 | 0.09 | 0.00 | 0.17 |
| 6.218 | 0.09 | 0.01 | 0.52 |
| 6.357 | 0.20 | 2.76 | 97.69 |
| 6.510 | 0.16 | 0.01 | 0.50 |
| 6.750 | 0.19 | 0.01 | 0.48 |

**h**

HRMS of Inactive Isomer Compound 1

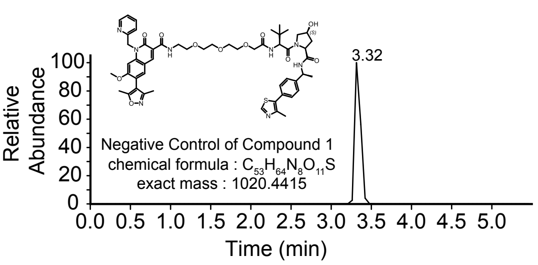


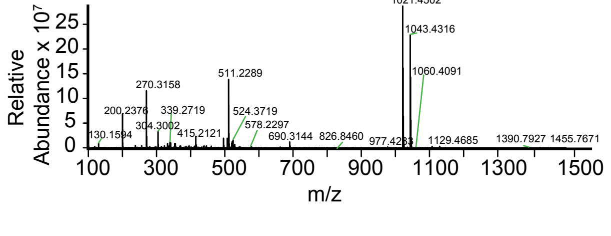


**i**

[1]H NMR spectrum of Inactive Isomer Compound 1

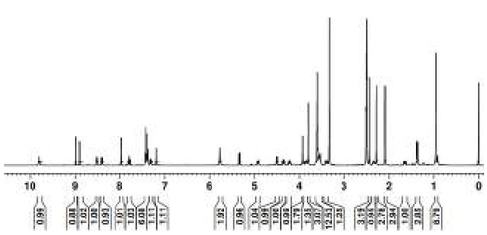

[13]C NMR spectrum of inactive Isomer Compound 1

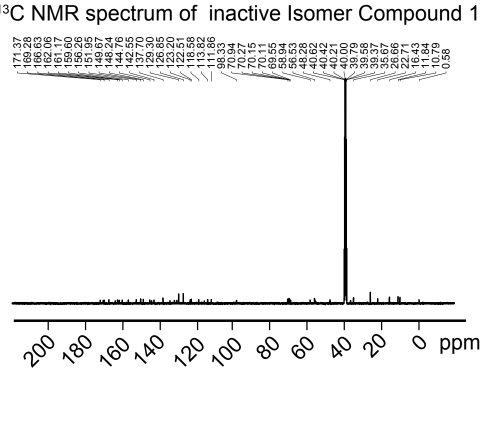


**Supplementary Figure 12**
