## supplementary Figure 13 for "Generative AI Framework SynGlue for the Rational Design of Clinically relevant Protein Degraders"

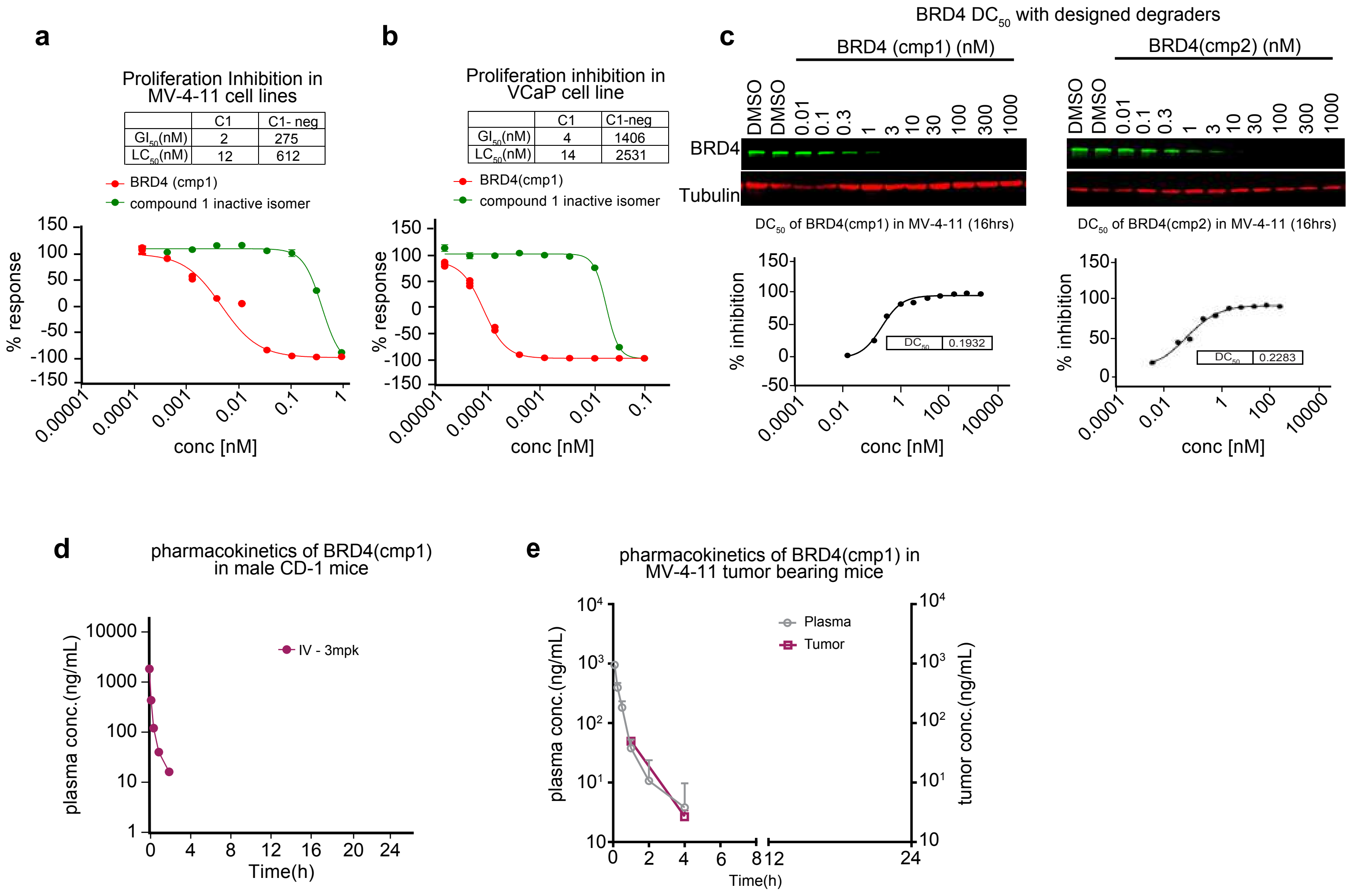


f

| Structure | 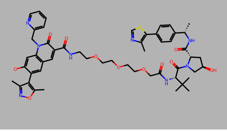 | 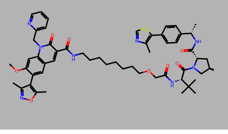 |
|---|---|---|
| Compound | Compound 1 | Compound 2 |
| MW/clogP | 1021/2.5 | 1017/5.1 |
| BRD4 BD1 $IC_{50}$(nM) | 84 | 744/TBR |
| MV-4-11 $GI_{50}$ (nM) | 1.1(n=2) | 7.6(n=2) |
| VCap $GI_{50}$nM/cMYC $IC_{50}$ | 3 (n=2) | 7(n=2) |
| LnCap $GI_{50}$nM/cMYC $IC_{50}$ | 11(n=2) | 9(n=2) |
| MCF7 $GI_{50}$ (nM) | 12 | 18 |
| Eq.sol@pH 7.4(uM) | 4.53 | 0.3 |
| MiLM% rem @30/60 min | 35 / 8 (15/60 min.) | 45 / 21 |
| PPB mouse : % bound | 99 | ND |
| Caco2 permability/efflux | Poor permeability/ high efflux | ND |
| Mouse PK IV, 3mpk — $t_{1/2}$(h) | 0.51 | 2.19 |
| Mouse PK IV, 3mpk — $C_0$(ng/mL) | 2472 | 3690 |
| Mouse PK IV, 3mpk — AUC(0-t)(hr*ng/mL) | 432 | 2811 |
| Mouse PK IV, 3mpk — Cl (mL/min/kg) | 113 | 17.5 |

**Supplementary Figure 13**
