## supplementary Figure 14 for "Generative AI Framework SynGlue for the Rational Design of Clinically relevant Protein Degraders"

### GSPT1 (cmp1) characterization

**a** HPLC of GSPT1(cmp1)

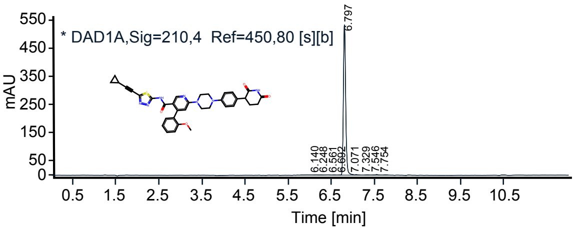


| RT[min] | Width[min] | Area | Area % |
|---|---|---|---|
| 6.140 | 0.13 | 2.46 | 0.12 |
| 6.248 | 0.11 | 1.39 | 0.07 |
| 6.561 | 0.13 | 1.80 | 0.09 |
| 6.692 | 0.07 | 1.89 | 0.09 |
| 6.797 | 0.33 | 2008.22 | 98.70 |
| 7.071 | 0.13 | 6.23 | 0.31 |
| 7.329 | 0.13 | 2.22 | 0.11 |
| 7.546 | 0.18 | 6.68 | 0.33 |
| 7.754 | 0.12 | 3.76 | 0.18 |

**b** LCMS of GSTP1 (cmp1)

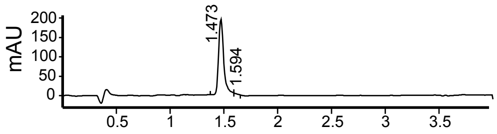


| Peak | RT | Area | Area % |
|---|---|---|---|
| 1 | 1.473 | 6.477e+002 | 98.689 |
| 2 | 1.594 | 8.600e+000 | 1.311 |

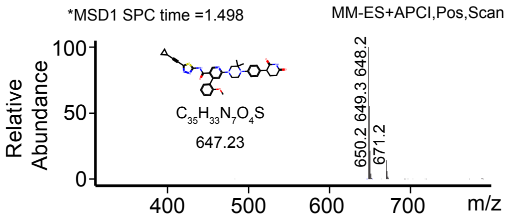


**c** $^1$H NMR of GSTP1 (cmp1)

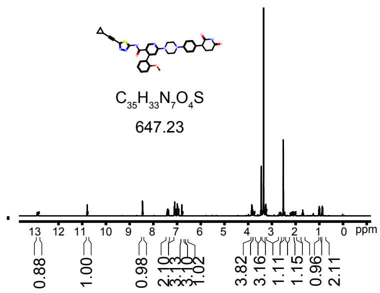


### GSPT1 (cmp2) characterization

**d** HPLC of GSTP1 (cmp2)

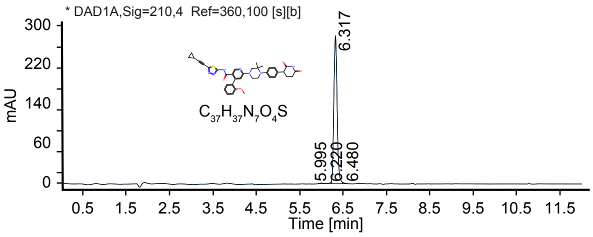


| RT[min] | Width[min] | Area | Area % |
|---|---|---|---|
| 5.995 | 0.17 | 4.45 | 0.30 |
| 6.220 | 0.10 | 5.42 | 0.36 |
| 6.317 | 0.26 | 1473.97 | 98.89 |
| 6.480 | 0.14 | 6.69 | 0.45 |

**e** LCMS of GSTP1 (cmp2)

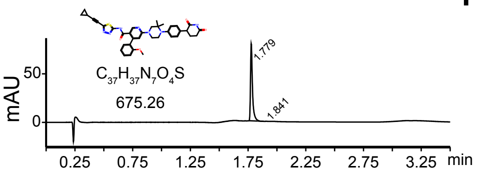


| Peak | RT | Area | Area % |
|---|---|---|---|
| 1 | 1.779 | 930254 | 99.555 |
| 2 | 1.849 | 4155 | 0.445 |

Peak #1 R.time :1.785
MasPeaks :498 Polarity:Positive
Spectrum Avg :1.778-1.792

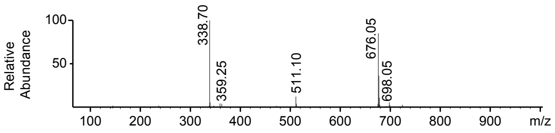


**f** $^1$H NMR of GSTP1 (cmp2)

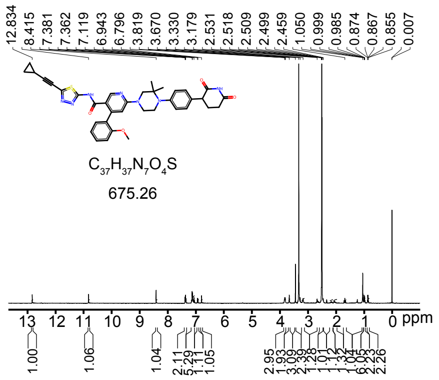


**Supplementary Figure 14**
