## Supplementary_Notes for "Generative AI Framework SynGlue for the Rational Design of Clinically relevant Protein Degraders"

### Supplementary Note 1

#### SynGlue: A Generative AI Platform for the Design and Validation of Polypharmacological Drugs and PROTACs

Saveena Solanki[1,†], Sanjay Kumar Mohanty[1,†], Shiva Satija[1], Sonam Chauhan[1], N.V.M. Rao Bandaru[3], Sandeep Dukare[3], Nirbhay Kumar Tiwari[3], Naveen Kumar R[3], Aravind A B[3], Subhendu Mukherjee [3], Dinesh Chikkanna, Wesley Roy Balasubramanian, Srinivasa Raju Sammeta, Vishakha Gautam[1], Sakshi Arora[1], Suvendu Kumar[1], Subhadeep Duari[1], Arushi Sharma[1], Raidhani Shome[1], Debarka Sengupta[1,2], Chandrasekhar Abbineni[3], Susanta Samajdar[3,#], Gaurav Ahuja[1,2,#]

[1]Department of Computational Biology, Indraprastha Institute of Information Technology-Delhi (IIIT-Delhi), Okhla, Phase III, New Delhi, 110020, India.
[2]Infosys Centre for AI, Indraprastha Institute of Information Technology-Delhi (IIIT-Delhi), Okhla, Phase III, New Delhi, 110020, India.
[3]Aurigene Oncology Limited, Electronic City Phase II, Bangalore, India.

[†]Shared First Authors

**[#]Correspondence:**

Gaurav Ahuja;

Susanta Samajdar;

#### General method

Starting materials and solvents of reagent grade were purchased from commercial suppliers and used without further purification. All non-aqueous reactions were performed under a nitrogen atmosphere in oven-dried glassware with magnetic stirring. The reaction progress was monitored by TLC on $SiO_2$. TLC was carried out employing silica gel 60 $F_{254}$ plates (Merck, Darmstadt), and developed chromatograms were visualized by UV (254 nm). $^{1}$H NMR and $^{13}$C NMR spectra were recorded at room temperature at 400 MHz and 101 MHz, respectively. $^{1}$H NMR spectra were recorded using tetramethyl silane (TMS) as an internal reference, and $^{13}$C NMR spectra were recorded using the deuterated DMSO peak as an internal reference. Chemical shifts are expressed in δ (ppm), and *J* values are given in hertz (Hz). Liquid chromatography mass spectra (LCMS) were obtained from Agilent 1100-LC/MSD VL using mobile phase: 0.1% formic acid- acid-acetonitrile or water-methanol; ionization was achieved either by positive or negative mode. High-Resolution Mass Spectroscopy (HRMS) analysis was carried out on a Thermo Orbitrap Q-Exactive plus LC-MS/MS high-resolution mass spectrometer. Data was acquired with the electrospray ionization (ESI) method using an H-ESI source hyphenated with a Dionex UHPLC system. The purity of final compounds was determined by high-performance liquid chromatography (HPLC).

#### Chemical Synthesis and Characterization of BRD4 compounds

- AU-14555 , AU-16914, AU-14849

#### Scheme-1: Synthesis of intermediate-9:

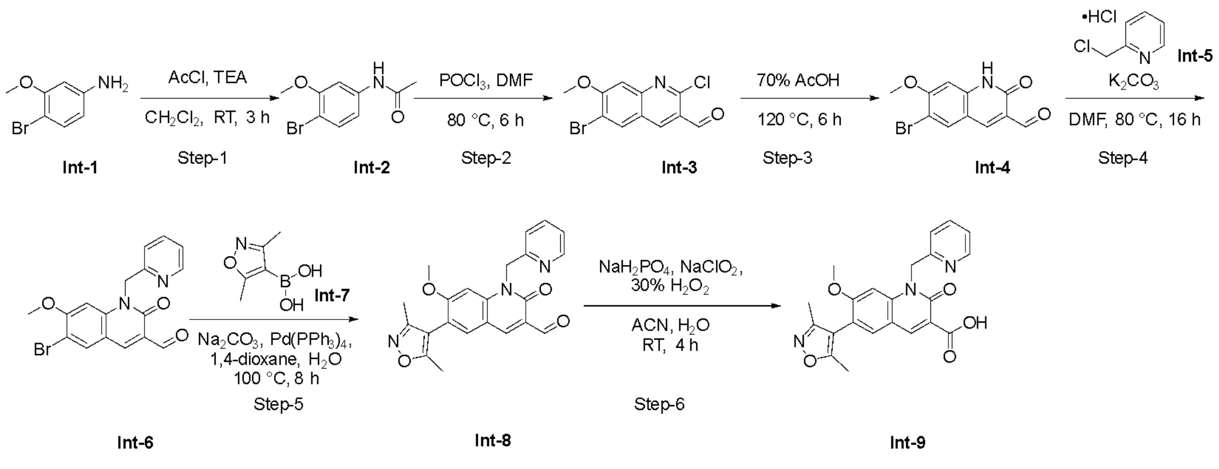

**Step-1: Synthesis of *N*-(4-Bromo-3-methoxyphenyl) acetamide (Intermediate-2)**

To an ice-cooled solution of 4-bromo-3-methoxyaniline **(Intermediate-1)** (2.0 g, 9.90 mmol) in $CH_2Cl_2$ (25 mL) was added triethylamine (4.1 mL, 29.7 mmol) followed by stirring for 5 min at room temperature. Acetyl chloride (1.05 mL, 14.85 mmol) was added to the reaction mixture and stirred for 3 h at room temperature. The reaction mixture was quenched by addition of aqueous $NaHCO_3$ solution (up to pH ~ 8). The mixture was extracted with $CH_2Cl_2$ (2 x 200 mL) and combined organic layers were washed with water (200 mL) and brine (200 mL), dried over sodium sulphate and concentrated under reduced pressure. The obtained **intermediate-2** (2.5 g, crude yield) was directly used for the next step without further purification; $^1$H NMR (400 MHz, DMSO-$d_6$) $\delta$ 10.06 (s, 1H), 7.45–7.43 (m, 2H), 7.10 (dd, $J$ = 8.3 Hz, 2.0 Hz, 1H), 3.79 (s, 3H), 2.04 (s, 3H); LC-MS (*m/z*): calculated for $C_9H_{11}BrNO_2$, 244.0 $[M + H]^+$, found 244.1.

**Step-2: Synthesis of 6-Bromo-2-chloro-7-methoxyquinoline-3-carbaldehyde (Intermediate-3)**

$POCl_3$ (7.6 mL, 81.96 mmol) was added dropwise to DMF (2.5 mL, 32.78 mmol) at 0 °C and stirred for 5 min. *N*-(4-bromo-3-methoxyphenyl) acetamide **(Intermediate-2)** (2.0 g, 8.26 mmol) was added to the reaction mixture and the resulting solution was heated to 80 °C for 16 h. The reaction mixture was cooled to room temperature and poured into crushed ice and extracted with EtOAc (2 x 200 mL). The combined organic layers were washed with water (200 mL), brine (200 mL), dried over sodium sulphate and concentrated under reduced pressure to get i**ntermediate-3** (2.0 g, crude yield) which was directly used for the next step without further purification.; $^1$H NMR (400 MHz, DMSO-$d_6$) $\delta$ 10.33 (s, 1H), 8.88 (s, 1H), 8.64 (s, 1H), 7.59 (s, 1H), 4.07 (s, 3H); LC-MS (*m/z*): calculated for $C_{11}H_8BrClNO_2$, 299.9 $[M + H]^+$, found 300.

**Step-3: Synthesis of 6-Bromo-7-methoxy-2-oxo-1,2-dihydroquinoline-3-carbaldehyde (Intermediate-4)**

A suspension of 6-bromo-2-chloro-7-methoxyquinoline-3-carbaldehyde (**Intermediate-3**) (2.0 g, 6.65 mmol) in 70 % acetic acid (40 mL) was heated to reflux for 6 h. Upon cooling the reaction mixture to room temperature a solid product was precipitated which was filtered and washed with water and dried under reduced pressure to afford the **intermediate -4** as brown solid (1.5 g, 80%); $^1$H NMR (400 MHz, DMSO-$d_6$) $\delta$ 12.18 (s, 1H), 10.17 (s, 1H), 8.42 (s, 1H), 8.22 (s, 1H),

6.93 (s, 1H), 3.94 (s, 3H); LC-MS (*m/z*): calculated for $C_{11}H_9BrNO_3$, 282.0 [M + H]$^+$, found 282.1.

**Step-4: Synthesis of 6-Bromo-7-methoxy-2-oxo-1-(pyridin-2-ylmethyl)-1,2-dihydroquinoline-3-carbaldehyde (Intermediate-6)**

To a solution of 6-bromo-7-methoxy-2-oxo-1, 2-dihydroquinoline-3-carbaldehyde (**Intermediate-4)** (9 g, 31.91 mmol) in DMF (80 mL) were added potassium carbonate (13.2 g, 95.73 mmol) followed by 2-(chloromethyl) pyridine hydrochloride (**Intermediate-5)** (6.4 g, 35.1 mmol) and stirred at 80 ºC for 16 h. Reaction mixture was diluted with water and extracted with EtOAc (2 x 400 mL). The combined organic layers were washed with water (400 mL), brine (300 mL), dried over sodium sulphate and concentrated under reduced pressure. The obtained i**ntermediate-6** was directly used for the next step without further purification (7.5 g, crude); $^1$H NMR (400 MHz, DMSO-$d_6$) δ 10.25 (s, 1H), 8.51–8.48 (m, 2H), 8.31 (s, 1H), 7.78 (t, *J* = 7.9 Hz, 1H), 7.40 (d, *J* = 8.4 Hz, 1H), 7.31–7.28 (m, 1H), 7.13 (s, 1H), 5.70 (s, 2H), 3.86 (s, 3H); LC-MS (*m/z*): calculated for $C_{17}H_{14}BrN_2O_3$ 373.0 [M + H]$^+$, found 373.0.

**Step-5: Synthesis of 6-(3,5-Dimethylisoxazol-4-yl)-7-methoxy-2-oxo-1-(pyridin-2-ylmethyl)-1,2-dihydroquinoline-3-carbaldehyde (Intermediate-8)**

To a solution of 6-bromo-7-methoxy-2-oxo-1-(pyridin-2-ylmethyl)-1,2-dihydro quinoline-3-carbaldehyde (**Intermediate-6)** (4.0 g, 10.72 mmol) in 1,4-dioxane (40 mL) and $H_2O$ (10 mL) were added 3,5-dimethylisoxazoleboronic acid (**Intermediate-7)** (2.30 g, 16.08 mmol) and sodium carbonate (3.41 g, 32.16 mmol) followed by degassing with nitrogen purging for 20 min. Tetrakis(triphenylphosphine)palladium (2.47 g, 2.14 mmol) was added followed by heated at 100 °C for 8 h. The reaction mixture was concentrated under reduced pressure. The residue was diluted with EtOAc (200 ml), washed with water (200 mL), brine (200 mL), dried over sodium sulphate and concentrated in vacuo. The residue was washed with hexane to give the title **intermediate-8** as yellow solid (3.2 g, 76 %); $^1$H NMR (400 MHz, DMSO-$d_6$) δ 10.28 (s, 1H), 8.54 (s, 1H), 8.52 (d, *J* = 4.4 Hz, 1H), 7.94 (s, 1H), 7.82–7.77 (m, 1H), 7.44 (d, *J* = 7.8 Hz, 1H), 7.33–7.29 (m, 1H), 7.17 (s, 1H), 5.72 (s, 2H), 3.81 (s, 3H), 2.27 (s, 3H), 2.08 (s, 3H); LC-MS (*m/z*): calculated for $C_{20}H_{20}N_3O_4$ 390.1 [M + H]$^+$, found 390.1.

**Step-6: Synthesis of 6-(3, 5-Dimethylisoxazol-4-yl)-7-methoxy-2-oxo-1-(pyridin-2-ylmethyl)-1, 2-dihydroquinoline-3-carboxylic acid (Intermediate-9)**

To a solution of 6-(3,5-dimethylisoxazol-4-yl)-7-methoxy-2-oxo-1-(pyridin-2-ylmethyl)-1,2-dihydroquinoline-3-carbaldehyde (**Intermediate-8)** (1.2 g, 3.08 mmol) in a mixture of acetonitrile (12 mL) and $H_2O$ (6 mL) were added sodium dihydrogen phosphate (1.6 g), hydrogen peroxide 30 % (1 mL) and sodium chlorite (0.85 g). The reaction mixture was stirred at room temperature for 4 h. The reaction mixture was poured into crushed ice and the solid was isolated by filtering. The solid was washed thoroughly with water and vacuum dried to give the title **intermediate-9** as yellow solid (0.85 g, 68 %); $^1H$ NMR (400 MHz, DMSO-$d_6$) $\delta$ 14.36 (s, 1H), 8.99 (s, 1H), 8.50 (d, $J$ = 4.4 Hz, 1H), 8.05 (s, 1H), 7.84–7.80 (m, 1H), 7.51 (d, $J$ = 7.8 Hz, 1H), 7.34–7.31 (m, 1H), 7.28 (s, 1H), 5.84 (s, 2H), 3.84 (s, 3H), 2.28 (s, 3H), 2.09 (s, 3H); LC-MS (*m/z*): calculated for $C_{22}H_{20}N_3O_5$ 406.1 $[M + H]^+$, found 406.2.

**Scheme-2: Synthesis of AU-14555 (BRD4 Compound 1) :**

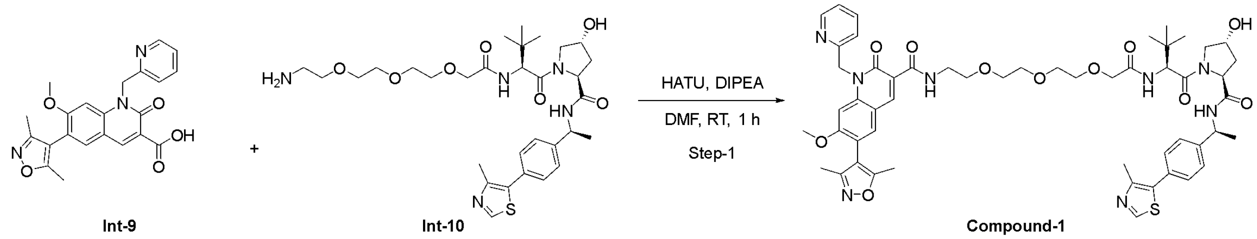


Int-9 Int-10 Compound-1

**Step-1: Synthesis of 6-(3,5-Dimethylisoxazol-4-yl)-*N*-((*S*)-13-((2*S*,4*R*)-4-hydroxy-2-(((*S*)-1-(4-(4-methylthiazol-5-yl)phenyl)ethyl)carbamoyl)pyrrolidine-1-carbonyl)-14,14-dimethyl-11-oxo-3,6,9-trioxa-12-azapentadecyl)-7-methoxy-2-oxo-1-(pyridin-2-ylmethyl)-1,2-dihydroquinoline-3-carboxamide: (AU-14555)**

To a solution of 6-(3,5-dimethylisoxazol-4-yl)-7-methoxy-2-oxo-1-(pyridin-2-ylmethyl)-1,2-dihydroquinoline-3-carboxylic acid **(Intermediate-9)** (1.4 g, 3.47 mmol) and (2*S*,4*R*)-1-((*S*)-14-amino-2-(*tert*-butyl)-4-oxo-6,9,12-trioxa-3-azatetradecanoyl)-4-hydroxy-*N*-((S

)-1-(4-(4-methylthiazol-5-yl)phenyl)ethyl)pyrrolidine-2-carboxamide (**Intermediate-10)** (2 g, 3.15 mmol) in DMF (30.0 mL) were added HATU (1.8 g, 4.73 mmol) and DIPEA (2.2 mL, 12.62 mmol) at 0 $^0$C. The mixture was stirred at room temperature for 1 h. TLC was monitored, and the reaction mixture was diluted with aqueous $NH_4Cl$ (50 mL) and extracted with EtOAc (2 x 100 mL). The organic layer was dried over sodium sulphate and concentrated under reduced pressure. The residue was purified by CombiFlash® to afford the tittle compound **(AU-14555)** as off-white solid (Yield: 1.7 g, 53%); $^1$H NMR (400 MHz, DMSO-$d_6$): $\delta$ $^1$H NMR (400 MHz, DMSO-$d_6$): $\delta$ 9.81 (t, $J$ = 5.2 Hz, 1H), 8.96 (s, 1H), 8.89 (s, 1H), 8.51-8.48 (m, 1H), 8.44 (d, $J$ = 8.0 Hz, 1H), 7.94 (s, 1H), 7.81-7.76 (m, 1H), 7.44-7.28 (m, 7H), 7.16 (s, 1H), 5.76 (s, 2H), 4.91-4.85 (m, 1H), 4.54 (d, $J$ = 9.4 Hz, 1H), 4.44 (t, $J$ = 8.4 Hz, 1H), 4.29 (brs, 1H), 3.93 (s, 2H), 3.79 (s, 3H), 3.61-3.52 (m, 14H), 2.44 (s, 3H), 2.26 (s, 3H), 2.07 (s, 3H), 2.05-2.02 (m, 1H), 1.81-1.74 (m, 1H), 1.36 (d, $J$ = 6.8 Hz, 3H), 0.93 (s, 9H); $^{13}$C NMR (101 MHz, DMSO-$d_6$): $\delta$ 170. 98, 169.48, 169.10, 166.68, 163.41, 162.06, 161.19, 159.63, 156.16, 151.97, 149.64, 148.18, 145.11, 143.85, 142.53, 137.77, 133.93, 131.58, 130.12, 129.28, 126.79, 126.73, 123.25, 122.52, 118.52, 115.60, 113.80, 111.84, 98.29, 70.28, 70.15, 70.09, 70.02, 69.50, 69.24, 59.04, 56.97, 56.51, 56.19, 48.24, 48.01, 38.13, 36.19, 26.72, 26.64, 22.85, 16.39, 11.80, 10.76 ; HPLC: 98.37%; HR-MS ($m/z$): calculated for $C_{53}H_{65}N_8O_{11}S$, 1021.44935 $[M + H]^+$, found, 1021.44696.

**Scheme-3: Synthesis of AU-14849 (BRD4 Compound 2)**

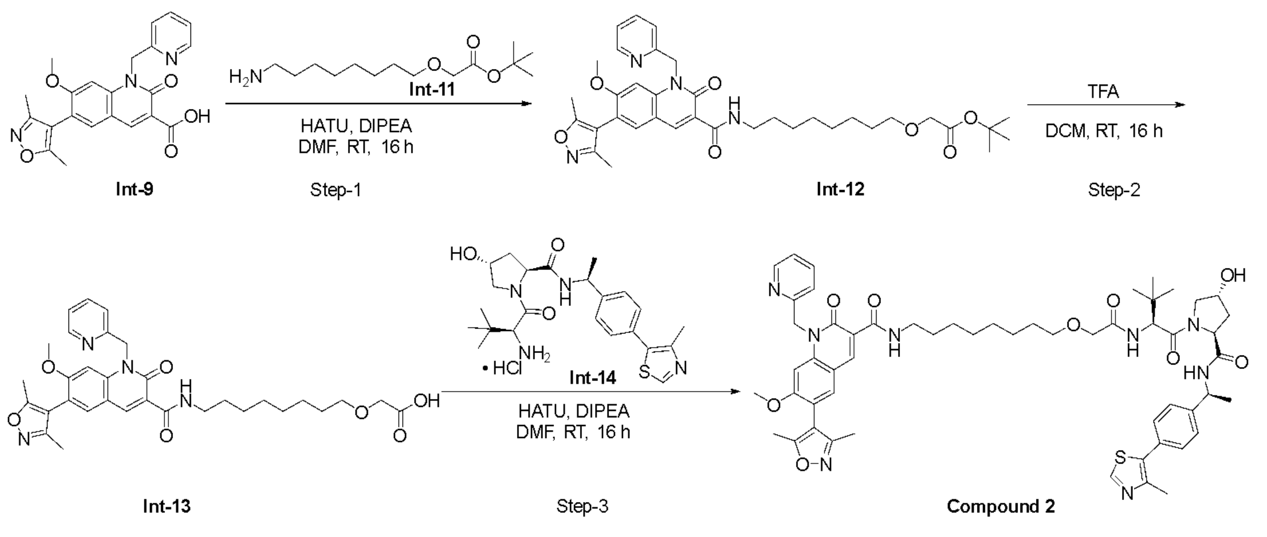

**Step-1: Synthesis of *tert*-butyl 2-((8-(6-(3,5-dimethylisoxazol-4-yl)-7-methoxy-2-oxo-1-(pyridin-2-ylmethyl)-1,2-dihydroquinoline-3-carboxamido)octyl)oxy)acetate (Intermediate-12)**

To a solution of 6-(3,5-dimethylisoxazol-4-yl)-7-methoxy-2-oxo-1-(pyridin-2-ylmethyl)-1,2-dihydroquinoline-3-carboxylic acid (**Intermediate-9)** (1 g, 2.46 mmol) and *tert*-butyl 2-((8-aminooctyl)oxy)acetate (**Intermediate-11)** (0.64 g, 2.46 mmol) in DMF (10.0 mL) was added HATU (1.4 g, 3.69 mmol) followed by DIPEA (1.72 mL, 9.86 mmol) at 0 $^0$C. The mixture was stirred at room temperature for 16 h. TLC was monitored, and the reaction mixture was diluted with aqueous $NH_4Cl$ (60 mL) and extracted with $CH_2Cl_2$ (60 mL x 2). The organic layer was dried over sodium sulphate and concentrated under reduced pressure. The residue was purified by CombiFlash® to afford the **intermediate-12** as off-white solid (0.85 g, 53 %); LC-MS (*m/z*): calculated for $C_{36}H_{47}N_4O_7$ 647.3 $[M + H]^+$, found, 647.2

**Step-2: Synthesis of 2-((8-(6-(3,5-dimethylisoxazol-4-yl)-7-methoxy-2-oxo-1-(pyridin-2-ylmethyl)-1,2-dihydroquinoline-3-carboxamido)octyl)oxy)acetic acid trifluoro acetate (Intermediate -13)**

To the stirred solution of *tert*-Butyl 2-((11-(6-(3,5-dimethylisoxazol-4-yl)-7-methoxy-2-oxo-1-(pyridin-2-ylmethyl)-1,2-dihydroquinoline-3-carboxamido)undecyl)oxy)acetate (0.85 g, 1.31 mmol) **(Intermediate-12)** in $CH_2Cl_2$ (10 mL) was added TFA (0.74g, 6.55 mmol) at 0 $^0$C, then stirred at room temperature for 16h. Reaction was monitored by TLC. After completion, the reaction mixture was concentrated under reduced pressure to afford the **intermediate-13** (0.7 g, crude); LC-MS (*m/z*): calculated for $C_{32}H_{39}N_4O_7$ 591.3 $[M + H]^+$, found 591.3.

**Step-3: Synthesis of 6-(3,5-dimethylisoxazol-4-yl)-*N*-(8-(2-(((*S*)-1-((2*S*,4*R*)-4-hydroxy-2-(((*S*)-1-(4-(4-methylthiazol-5-yl)phenyl)ethyl)carbamoyl)pyrrolidin-1-yl)-3,3-dimethyl-1-oxobutan-2-yl)amino)-2-oxoethoxy)octyl)-7-methoxy-2-oxo-1-(pyridin-2-ylmethyl)-1,2-dihydroquinoline-3-carboxamide (AU-14849)**

To a solution of 2-((8-(6-(3,5-dimethylisoxazol-4-yl)-7-methoxy-2-oxo-1-(pyridin-2-ylmethyl)-1,2-dihydroquinoline-3-carboxamido)octyl)oxy)acetic acid trifluoro acetate **(Intermediate-13)** (0.5 g, 0.84 mmol) and (2*S*,4*R*)-1-((*S*)-2-amino-3,3-dimethylbutanoyl)-4-hydroxy-*N*-((S)-1-(4-(4-methylthiazol-5-yl)phenyl)ethyl)pyrrolidine-2-carboxamide hydrochloride (**Intermediate-14)** (0.4 g, 0.84 mmol) in DMF (10 mL) was added HATU (0.483 g, 1.26 mmol) followed by DIPEA (0.59 mL, 3.38 mmol) at 0 $^0$C. The mixture was stirred at room temperature for 16 h. TLC was monitored, and the reaction mixture was diluted with aqueous $NH_4Cl$ (10 mL) solution and extracted with $CH_2Cl_2$ (2 x 20 mL). The organic layer was dried over sodium sulphate and concentrated under reduced pressure. The residue was purified by CombiFlash® to afford the **compound-2 (AU-14849)** as off-white solid (Yield: 0.52 g, 60%); $^1$H NMR (400 MH*z*, DMSO-$d_6$): *δ* 9.67–9.64 (m, 1H), 8.98 (d, *J* = 3.4 Hz, 1H), 8.88 (s, 1H), 8.50 (d, *J* = 4.4 Hz, 1H), 8.42 (d, *J* = 7.8 Hz, 1H), 7.95 (*s*, 1H), 7.80–7.76 (m, 1H), 7.42–7.34 (m, 5H), 7.31–7.28 (m, 2H), 7.18 (s, 1H), 5.75 (s, 2H), 5.12 (d, *J* = 3.5 Hz, 1H), 4.91–4.88 (m, 1H), 4.53 (d, *J* = 9.3 Hz, 1H), 4.45 (t, *J* = 8.3 Hz, 1H), 4.29–4.27 (m, 1H), 3.89 (s, 2H), 3.79 (s, 3H), 3.59–3.51 (m, 2H), 3.48–3.44 (m, 2H), 2.44 (s, 3H), 2.26 (s, 3H), 2.08 (s, 3H), 2.06–2.04 (m, 1H), 1.77–1.75 (m, 1H), 1.47–1.45 (m, 5H), 1.37–1.23 (m, 12H), 0.93(s, 9H); $^{13}$C NMR (101 MHz, DMSO-$d_6$): *δ* 170. 94, 169.48, 169.05, 166.67, 163.22, 162.14, 161.13, 159.63, 156.18, 151.96, 149.65, 148.18, 145.15, 143.68, 142.44, 137.75, 133.88, 131.58, 130.11, 129.35, 129.27, 126.77, 123.24, 122.51, 118.71, 115.59, 113.83, 111.85, 98.30, 71.40, 69.83, 69.24, 59.02, 57.01, 56.50, 56.08, 48.25, 48.05, 38.14, 36.65, 36.30, 29.55, 29.47, 29.20, 29.15, 26.92, 26.62, 26.03, 22.86, 16.39, 11.79, 10.76; HPLC: 98.26%; HR-MS (*m/z*): calculated for $C_{55}H_{69}N_8O_9S$, 1017.49082 $[M + H]^+$, found 1017.48822.

**Scheme-4: Synthesis of negative control of BRD4 Compound 1(AU-16914):**

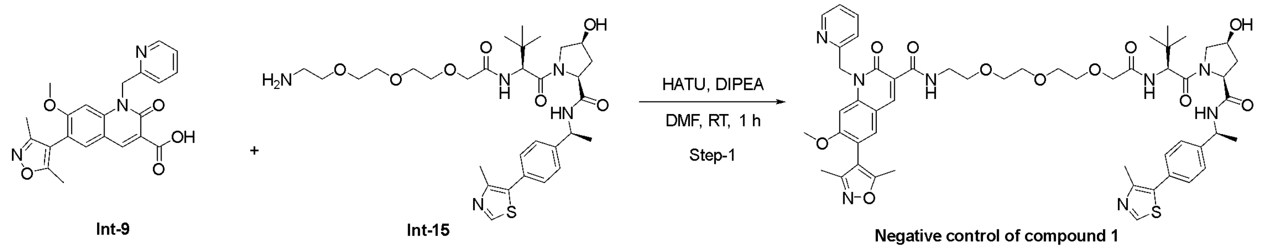


Int-9 Int-15 Negative control of compound 1

**Step-1: Synthesis of 6-(3,5-dimethylisoxazol-4-yl)-N-((S)-13-((2S,4S)-4-hydroxy-2-(((S)-1-(4-(4-methylthiazol-5-yl)phenyl)ethyl)carbamoyl)pyrrolidine-1-carbonyl)-14,14-dimethyl-11-oxo-3,6,9-trioxa-12-azapentadecyl)-7-methoxy-2-oxo-1-(pyridin-2-ylmethyl)-1,2-dihydroquinoline-3-carboxamide : (**Negative control of Compound **1)**

To a solution of 6-(3,5-dimethylisoxazol-4-yl)-7-methoxy-2-oxo-1-(pyridin-2-ylmethyl)-1,2-dihydroquinoline-3-carboxylic acid (**Intermediate-9)** (0.13 g, 0.32 mmol) and (2*S*,4*R*)-1-((*S*)-14-amino-2-(*tert*-butyl)-4-oxo-6,9,12-trioxa-3-azatetradecanoyl)-4-hydroxy-*N*-((S)-1-(4-(4-methylthiazol-5-yl)phenyl)ethyl)pyrrolidine-2-carboxamide (**Intermediate-15**[1]**)** (0.18 g, 0.28 mmol) in DMF (10.0 mL) were added HATU (0.18 g, 0.47 mmol) and DIPEA (0.18 mL, 1.00 mmol) at 0 $^0$C. The mixture was stirred at room temperature for 1 h. TLC was monitored, and the reaction mixture was diluted with aqueous $NH_4Cl$ (20 mL) and extracted with EtOAc (2 x 50 mL). The organic layer was dried over sodium sulphate and concentrated under reduced pressure. The residue was purified by CombiFlash® to afford the negative control of Compound 1 as off-white solid (Yield: 0.2 g, 71%); $\delta$ $^1$H NMR (400 MHz, DMSO-$d_6$): $\delta$ 9.81 (t, $J$ = 5.2 Hz, 1H), 8.96 (s, 1H), 8.91 (s, 1H), 8.52-8.49 (m, 1H), 8.40 (d, $J$ = 7.6 Hz, 1H), 7.96 (s, 1H), 7.81-7.76 (m, 1H), 7.44-7.36 (m, 6H), 7.32-7.29 (m, 1H), 7.18 (s, 1H), 5.77 (s, 2H), 5.34-5.32 (m, 1H), 4.94-4.88 (m, 1H), 4.50 (d, $J$ = 9.2 Hz, 1H), 4.37-4.33 (m, 1H), 4.23-4.18 (m, 1H), 3.93 (s, 2H), 3.89-3.83 (m, 1H), 3.80 (s, 3H), 3.65-3.52 (m, 12H), 3.41-3.39 (m, 1H), 2.45 (s, 3H), 2.37-2.31 (m, 1H), 2.27 (s, 3H), 2.09 (s, 3H), 1.67-1.65 (m, 1H), 1.37 (d, $J$ = 6.8 Hz, 3H), 0.95 (s, 9H); $^{13}$C NMR (101 MHz, DMSO-$d_6$): $\delta$ 171. 37, 169.28, 166.63, 162.06, 161.17, 159.60, 156.26, 151.95, 149.67, 148.24, 144.76, 142.55, 137.70, 129.30, 126.85, 123.20, 122.51, 118.58, 113.82, 111.86, 98.33, 70.94, 70.27, 70.15, 70.11, 69.55, 58.94, 56.53, 48.28, 39.37, 35.67, 26.66, 22.71, 16.43, 11.84, 10.79 ; HPLC: 97.69%; HR-MS (*m/z*): calculated for $C_{53}H_{65}N_8O_{11}S$, 1021.44935 $[M + H]^+$, found, 1021.4502.

#### Chemical Synthesis and Characterization of GSPT1 compounds

- AU-29140, AU-30489.

**Scheme-5: Synthesis of intermediate-20:**

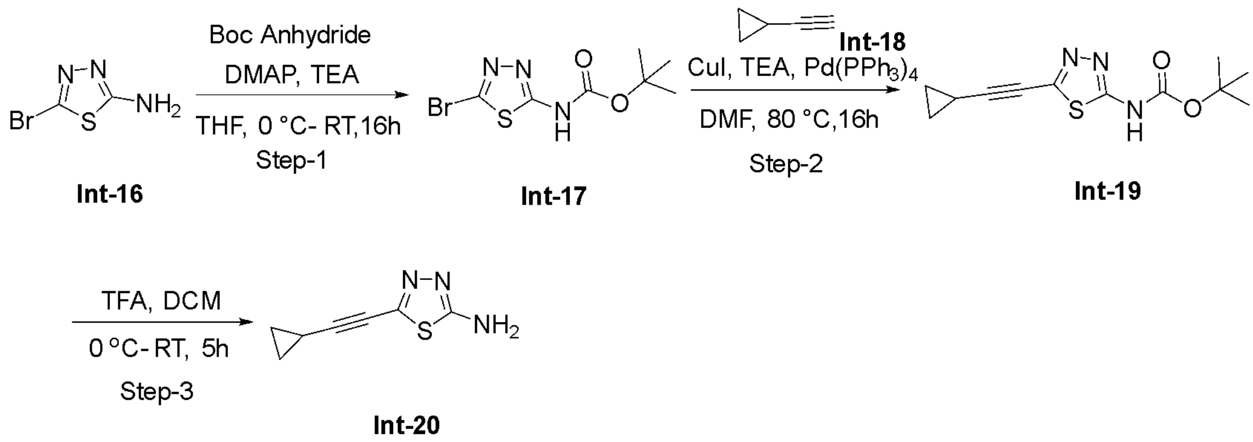


**Step-1: Synthesis of tert-butyl (5-bromo-1,3,4-thiadiazol-2-yl)carbamate (Intermediate-17)**

To a solution of 5-bromo-1,3,4-thiadiazol-2-amine **(Intermediate-16)** (1.0 g, 5.554 mmol) in 10 mL of THF were added DMAP (0.14 g, 1.110 mmol), triethylamine (1.12 g, 11.110 mmol) at 0 °C, and then Boc-anhydride was slowly added to the reaction mixture at 0 °C. The reaction mixture was stirred for 16 hours under argon atmosphere at room temperature. Upon completion, the reaction mixture was quenched with chilled water and extracted with ethyl acetate. The ethyl acetate layer was washed with brine solution, dried over anhydrous sodium sulphate and concentrated in vacuo. The residue was purified by column chromatography on silica gel (20% ethyl acetate/hexane) to give the title compound **(Intermediate-17)** (0.65 g, 42%) of tert-butyl (5-bromo-1,3,4-thiadiazol-2-yl) carbamate. $^{1}$H NMR (400 MHz, DMSO-d6): δ 12.27 (bs, 1H), 1.54 (s, 9H); LCMS: 279.9[M]$^{+}$.

**Step-2: Synthesis of tert-butyl (5-(cyclopropyl ethynyl)-1,3,4-thiadiazol-2-yl) carbamate (Intermediate-19)**

To a solution of tert-butyl (5-bromo-1,3,4-thiadiazol-2-yl) carbamate **(Intermediate-17)** (0.65 g, 2.318 mmol) in 6 mL of DMF were added triethylamine (0.704 g, 6.950 mmol), Copper Iodide (0.044 g, 0.230 mmol) and degassed with argon for 10 minutes. Then ethynyl cyclopropane**(Intermediate-18)** (0.92 g, 13.913 mmol) and Tetrakis(triphenylphosphine) palladium (0) (0.268 g, 0.230 mmol) were added to the reaction mixture and heated to 80 °C. The reaction mixture continued for 16 hours at 80 °C. Upon completion, the reaction mixture was concentrated in vacuo. The residue was purified by column chromatography on silica gel (20% ethyl acetate/hexane) to give the title compound **(Intermediate-19)** (0.11 g, 18 %) of tert-butyl (5-(cyclopropyl ethynyl)-1,3,4-thiadiazol-2-yl) carbamate. $^{1}$H NMR (400 MHz, CDCl3): δ 8.23 (bs, 1H), 1.58 (s, 9H), 1.575– 1.56 (m, 1H), 1.0 – 0.95 (m, 4H); LCMS: 266.10 $[M+H]^{+}$.

**Step-3: Synthesis of 5-(cyclopropyl ethynyl)-1,3,4-thiadiazol-2-amine (Intermediate-20)**

To a solution of tert-butyl (5-(cyclopropyl ethynyl)-1,3,4-thiadiazol-2-yl) carbamate **(Intermediate-19)** (0.11 g, 0.415 mmol) in 2 mL of DCM was added trifluoro acetic acid (0.08 g, 0.830 mmol) at 0°C. The reaction mixture was stirred for 5 hours under argon atmosphere at room temperature. Upon completion, the reaction mixture was concentrated in vacuo. The obtained residue was diluted with ethyl acetate and water and basified with 2N NaOH. The organic layer was separated and washed with brine solution, dried over anhydrous sodium sulphate and concentrated in vacuo to give the title compound **(Intermediate-20)** (0.065 g, 95 %) of 5-(cyclopropyl ethynyl)-1,3,4-thiadiazol-2-amine. $^{1}$H NMR (400 MHz, DMSO-d6): δ 7.50 (s, 2H), 1.65 – 1.60 (m, 1H), 0.97– 0.94 (m, 2H), 0.93 – 0.82 (m, 2H); LCMS: 166.0 $[M+H]^{+}$.

LC-MS (*m/z*): calculated for $C_9H_{11}BrNO_2$, 244.0 $[M + H]^{+}$, found 244.1.

**Scheme-6: Synthesis of intermediate-25:**

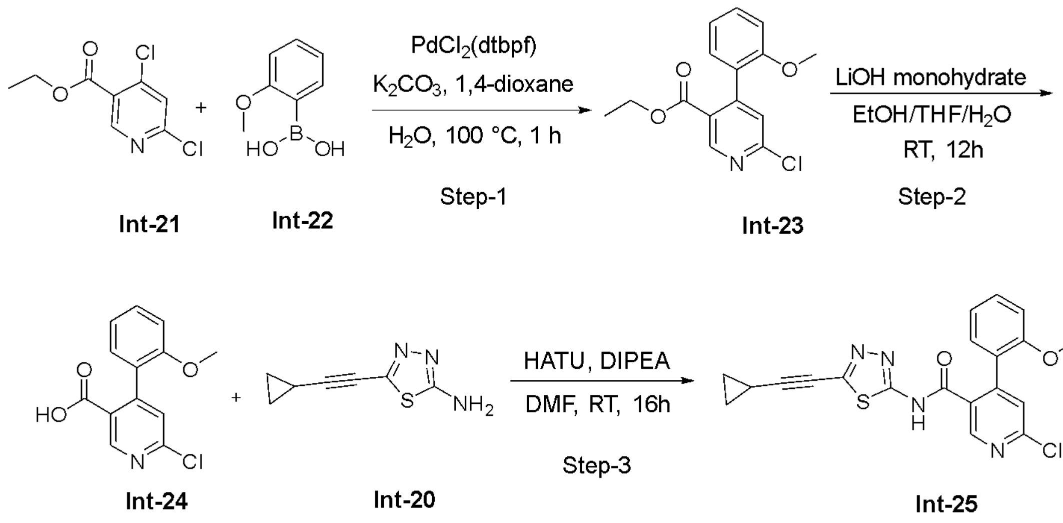


**Step-1: Synthesis of ethyl 6-chloro-4-(2-methoxyphenyl) nicotinate (Intermediate-23)**

To a solution of ethyl 4,6-dichloronicotinate **(Intermediate-21)** (3.2 g, 14.541 mmol) in 30 mL of 1,4-dioxane and 6 mL of water, was added (2-methoxyphenyl) boronic acid **(Intermediate-22)** (3.31 g, 21.812 mmol) and potassium carbonate (4.02 g, 29.080 mmol) and degassed with argon gas for 10 minutes. Then $PdCl_2$(dtbpf) (0.94 g, 1.454 mmol) was added to the reaction mixture and heated to 100 °C. The reaction mixture was stirred for 1 hour under argon atmosphere at 100°C and cooled to room temperature, filtered through celite bed and washed with ethyl acetate. The filtrate was washed with water, brine solution, dried over anhydrous sodium sulphate and concentrated in vacuo. The residue was purified by column chromatography on silica gel (30% ethyl acetate and hexane) to give the title compound **(Intermediate-23)** (2.5 g, 59%) of ethyl 6-chloro-4-(2-methoxyphenyl) nicotinate. $^1$H NMR (400 MHz, $CDCl_3$): δ 8.81 (s, 1H), 7.43 – 7.38 (m, 1H), 7.26 – 7.21 (m, 2H), 7.07 – 7.03 (m, 1H), 6.92 – 6.90 (m, 1H), 4.16 – 4.11 (m, 2H), 3.72 (s, 3H), 1.09 – 1.06 (m, 3H); LCMS: 291.95 $[M+H]^+$.

**Step-2: Synthesis of 6-chloro-4-(2-methoxyphenyl) nicotinic acid (Intermediate-24)**.

To a solution of ethyl 6-chloro-4-(2-methoxyphenyl) nicotinate **(Intermediate-23)** (1.2 g, 4.113 mmol) in 5 mL of ethanol was added 5 mL of THF, 5 mL of water and lithium hydroxide monohydrate (0.51 g, 12.330 mmol) at 0 °C. The reaction mixture was stirred for 12 hours at room temperature. Upon completion, the reaction mixture was concentrated in vacuo. The residue was diluted with chilled water, pH was adjusted to 2 with 1N HCl solution and extracted with ethyl acetate, dried over anhydrous sodium sulphate and concentrated in vacuo to give the title compound **(Intermediate-24)** (1.0 g, 92%) of 6-chloro-4-(2-methoxyphenyl) nicotinic acid. $^1$H NMR (400 MHz, CDCl3): δ 8.91 (s, 1H), 7.46 – 7.42 (m, 1H), 7.31 – 7.29 (m, 1H), 7.27 – 7.24 (m, 1H), 7.13 – 7.09 (m, 1H), 6.92 – 6.90 (m, 1H), 3.93 (s, 3H); LCMS: 263.95 $[M+H]^+$.

**Step-3: Synthesis of 6-chloro-N-(5-(cyclopropylethynyl)-1,3,4-thiadiazol-2-yl)-4-(2-methoxy phenyl) nicotinamide (Intermediate-25)**.

To a solution of 5-(cyclopropyl ethynyl)-1,3,4-thiadiazol-2-amine **(Intermediate-20)** (0.5 g, 1.895 mmol) and 6-chloro-4-(2,5-dimethoxyphenyl) nicotinic acid **(Intermediate-24)** (0.34 g, 2.085 mmol) in 10 mL of DMF was added HATU (0.67 g, 2.840 mmol) and N, N diisopropylethylamine (0.73 g, 5.680 mmol) at room temperature. The reaction mixture was stirred for 16 hours under argon atmosphere at room temperature. Upon completion, the reaction mixture was quenched with chilled water and extracted with ethyl acetate. The organic layer was washed with brine solution, dried over anhydrous sodium sulphate, and concentrated in vacuo. The residue was purified by column chromatography on silica gel (30% ethyl acetate/hexane) to give the title compound **(Intermediate-25)** (0.30 g, 38 %) of 6-chloro-N-(5-(cyclopropylethynyl)-1,3,4-thiadiazol-2-yl)-4-(2-methoxy phenyl)nicotinamide.$^1$H NMR (400 MHz, DMSO): δ 13.44 (bs, 1H), 8.69 (s, 1H), 7.61 (s, 1H), 7.45 – 7.41 (m, 2H), 7.11 – 7.09 (m, 1H), 6.99 – 6.97 (m, 1H), 3.45 (s, 3H), 1.71– 1.67 (m, 1H), 1.01 – 0.98 (m, 2H), 0.88 – 0.87 (m, 2H) ; LCMS: 411.20 $[M+H]^+$.

**Scheme-7: Synthesis of intermediate-32:**

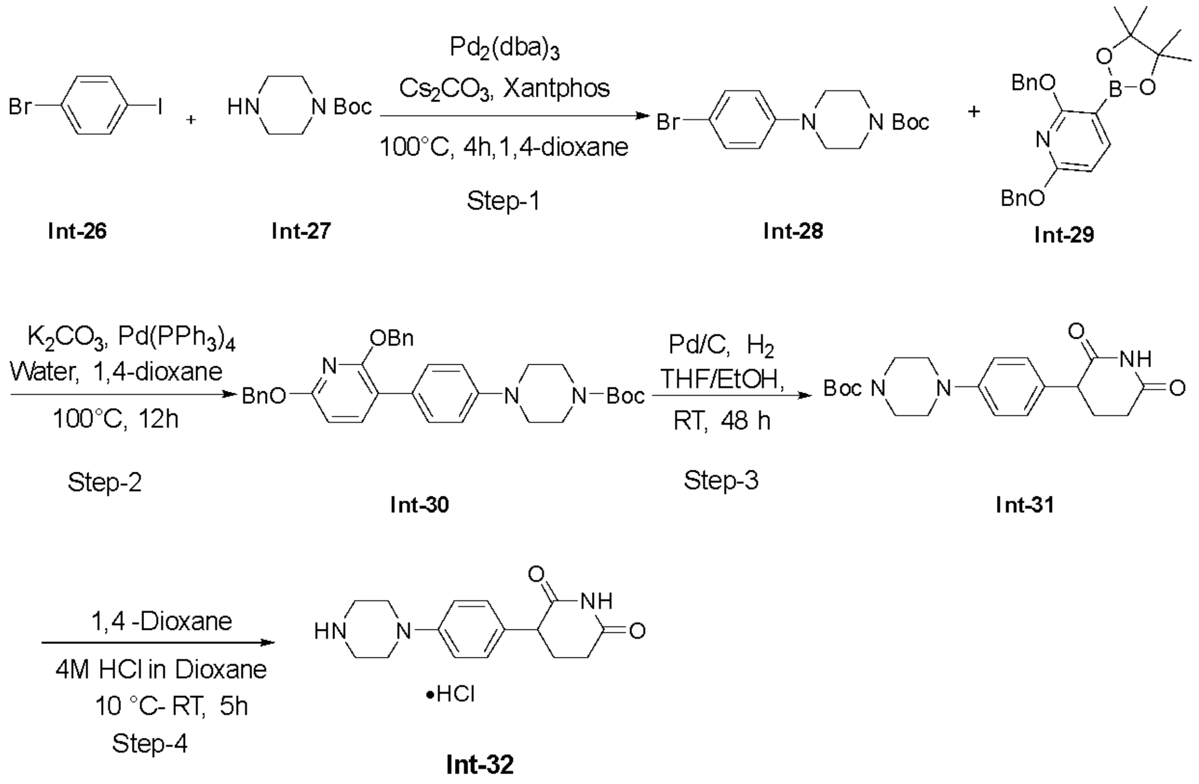


**Step-1: Synthesis of tert-butyl 4-(4-bromophenyl)piperazine-1-carboxylate (Intermediate-28)**

To a solution of 1-bromo-4-iodobenzene **(Intermediate-26)** (25.0 g, 88.367 mmol) and tert-butyl piperazine-1-carboxylate **(Intermediate-27)** (16.46 g, 88.367 mmol) in 300 mL of 1,4-dioxane were added cesium carbonate (71.98 g, 220.910 mmol), Xantphos (5.108 g, 8.830 mmol) and tris(dibenzylideneacetone)dipalladium (0) (0.992 g, 4.410 mmol). The reaction mixture was degassed with Argon gas for 20 minutes. The reaction mixture was stirred at 100 °C for 4 hours under argon atmosphere. Upon completion, the reaction mixture was filtered through celite bed. The filtrate was diluted with ethyl acetate and washed with water, brine solution, dried over anhydrous sodium sulphate and concentrated in vacuo. The residue was purified by column chromatography on silica gel (50% ethyl acetate/hexane) to give the title compound **(Intermediate-28)** (16.0 g, 53%) of tert-butyl 4-(4-bromophenyl)piperazine-1-carboxylate. $^1$H NMR (400 MHz, CDCl3): δ 7.38 – 7.36 (m, 2H), 6.82 – 6.79 (m, 2H), 3.60 – 3.57 (m, 4H), 3.13 – 3.10 (m, 4H), 1.55 (s, 9H); LCMS: 340.95 $[M]^+$.

**Step-2: Synthesis of tert-butyl 4-(4-(2,6-bis(benzyloxy)pyridin-3-yl)phenyl)piperazine-1-carboxylate (Intermediate-30)**

To a solution of tert-butyl 4-(4-bromophenyl)piperazine-1-carboxylate **(Intermediate-28)** (10.0 g, 29.303 mmol) and 2,6-bis(benzyloxy)-3-(4,4,5,5- tetramethyl-1,3,2-dioxaborolan-2-yl)pyridine **(Intermediate-29)** (14.67 g, 35.164 mmol) in 120 mL of 1,4-dioxane and 30 mL of water were added potassium carbonate (12.15 g, 97.900 mmol) and Pd(PPh3)4 (3.386 g, 2.930 mmol). The reaction mixture was degassed with argon gas for 20 minutes, then continued for 12 hours at 100 °C under argon atmosphere. The reaction mixture was filtered through celite bed, the filtrate was diluted with ethyl acetate and washed with water, brine solution, dried over anhydrous sodium sulphate and concentrated in vacuo. The residue was purified by column chromatography on silica gel (10% ethyl acetate and hexane) to give the title compound **(Intermediate-30)** (12.0 g, 74%) of tert-butyl 4-(4-(2,6-bis(benzyloxy)pyridin-3-yl)phenyl)piperazine-1-carboxylate. $^{1}$H NMR (400 MHz, CDCl3): δ 7.63 – 7.59 (m, 1H), 7.58 – 7.51 (m, 1H), 7.46 – 7.28 (m, 10H), 6.98 – 6.96 (m, 2H), 6.49 – 6.46 (m, 2H), 5.45 (s, 2H), 5.35 (s, 2H), 3.62 – 3.60 (m, 4H), 3.20 – 3.18 (m, 4H), 1.49 (s, 9H); LCMS: 552.30 $[M+H]^{+}$.

**Step-3: Synthesis of tert-butyl 4-(4-(2,6-dioxopiperidin-3-yl)phenyl)piperazine-1-carboxylate (Intermediate-31)**

To a solution of tert-butyl 4-(4-(2,6-bis(benzyloxy)pyridin-3-yl)phenyl)piperazine-1-carboxylate **(Intermediate-30)** (12.0 g, 21.751 mmol) in 100 mL of Ethanol and 100 mL of THF was added 10% Palladium on carbon (11.57 g, 108.750 mmol). The reaction mixture was stirred at room temperature under 80psi hydrogen gas pressure for 48 hours. Upon completion, the reaction mixture was filtered through celite bed, the filtrate was concentrated in vacuo to give the title compound **(Intermediate-31)** (8.0 g, 98%) of tert-butyl 4-(4-(2,6-dioxopiperidin-3-yl) phenyl)piperazine-1-carboxylate.$^{1}$H NMR (400 MHz, DMSO-d6): δ 10.77 (s, 1H), 7.08 – 7.06 (m, 2H), 6.92 – 6.90 (m, 2H), 3.76 – 3.72 (m, 1H), 3.46 – 3.44 (m, 4H), 3.09 – 3.06 (m, 4H), 2.68– 2.58 (m, 1H), 2.48– 2.44 (m, 1H), 2.16 –2.11 (m, 1H), 2.03 –1.98 (m, 1H), 1.42 (s, 9H); LCMS: 374.15 $[M+H]^{+}$.

**Step-4: Synthesis of 3-(4-(piperazin-1-yl)phenyl)piperidine-2,6-dione hydrochloride (Intermediate-32)**

To a solution of tert-butyl 4-(4-(2,6-dioxopiperidin-3-yl)phenyl)piperazine-1-carboxylate **(Intermediate-31)** (8.0 g, 21.422 mmol) in 40 mL of 1,4-dioxane was added 4M HCl in 1,4 dioxane (80 mL, 10v) at 10 °C under nitrogen atmosphere. The reaction mixture was stirred for 5 hours at room temperature. Upon completion, the reaction mixture was concentrated in vacuo. The residue was washed with diethyl ether and dried under reduced pressure to give the title compound **(Intermediate-32)** (6.3 g, 95%) of 3-(4-(piperazin-1-yl) phenoxy) piperidine-2,6-dione hydrochloride.$^{1}$H NMR (400 MHz, DMSO-d6) δ 10.79 (s, 1H), 9.26- 9.18 (bs, 2H), 7.12 – 7.10 (m, 2H), 6.97 – 6.95 (m, 2H), 3.78 – 3.74 (m, 1H), 3.39 – 3.34 (m, 4H), 3.23 – 3.18 (m, 4H), 2.68– 2.60 (m, 1H), 2.48– 2.44 (m, 1H), 2.16 –2.13 (m, 1H), 2.03 –1.98 (m, 1H); LCMS: 274.10 $[M+H]^+$.

**Scheme-8: Synthesis of intermediate-37:**

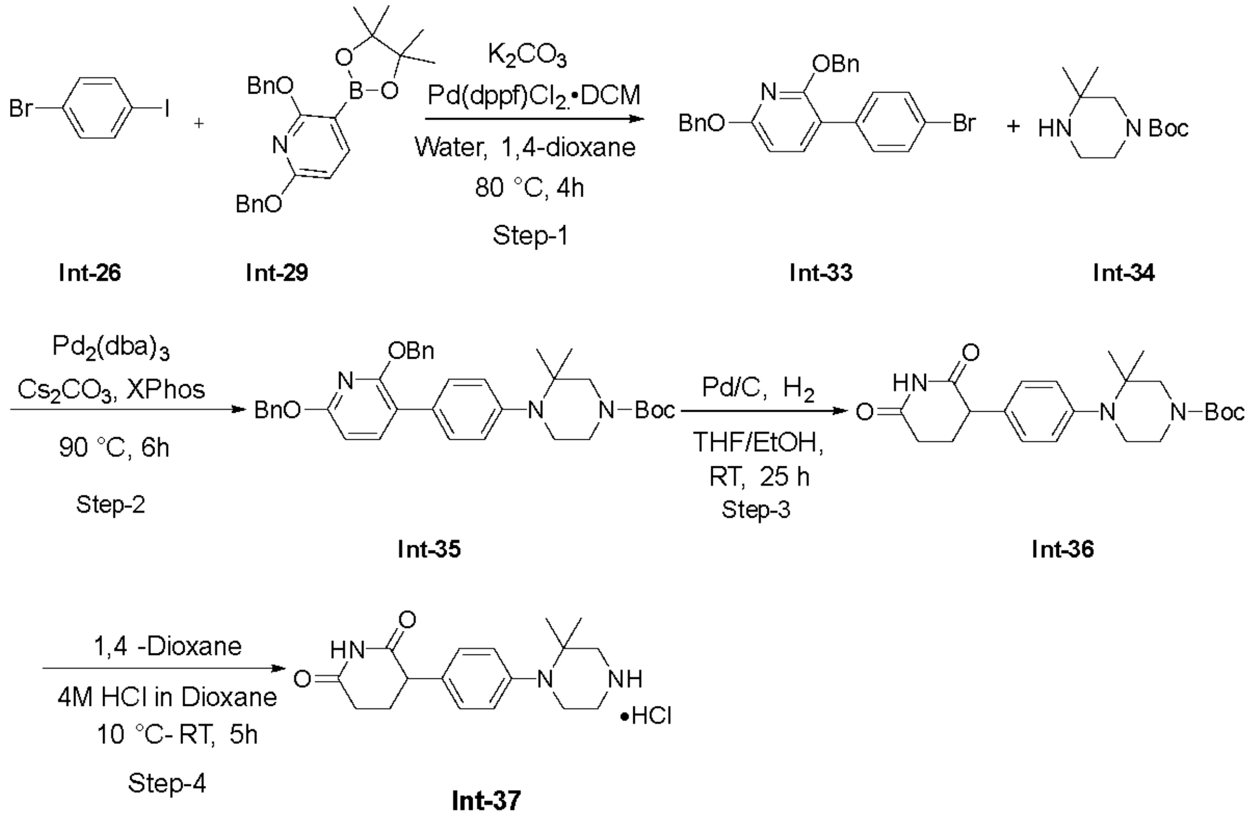


**Step-1: Synthesis of 2,6-bis(benzyloxy)-3-(4-bromophenyl) pyridine (Intermediate-33)**

To a solution of 1-bromo-4-iodobenzene **(compound-26)** (2.0 g, 7.069 mmol) and 2,6-bis(benzyloxy)-3-(4,4,5,5- tetramethyl-1,3,2-dioxaborolan-2- yl)pyridine **(compound-29)** (2.36 g, 5.655 mmol) in 24 mL of 1,4-dioxane and 6 mL of water were added potassium carbonate (2.44 g, 17.673 mmol), and Pd(dppf)$Cl_2$.DCM (0.57 g, 0.707 mmol). The reaction mixture was degassed with Argon gas for 20 minutes, then stirred at 80 °C for 4 hours under argon atmosphere. Upon completion the reaction mixture was filtered through celite bed, the filtrate was diluted with ethyl acetate and washed with water, brine solution, dried over anhydrous sodium sulphate and concentrated under reduced pressure. The residue was purified by column chromatography on silica gel (50% ethyl acetate/hexane) to give the title compound **(Intermediate-33)** (1.7 g, 54 %) of 2,6-bis(benzyloxy)-3-(4-bromophenyl)pyridine. $^1$H NMR (400 MHz, CDCl3): δ 7.58 – 7.53 (m, 1H), 7.52 – 7.51 (m, 2H), 7.46 – 7.45 (m, 4H), 7.40 – 7.30 (m, 8H), 6.51 – 6.49 (m, 1H), 5.44 (s, 2H), 5.39 (s, 2H); LCMS: 448.0 $[M+2]^+$.

**Step-2: Synthesis of tert-butyl 4-(4-(2,6-bis(benzyloxy)pyridin-3-yl)phenyl)-3,3-dimethylpiperazine-1-carboxylate (Intermediate-35)**

To a solution of 2,6-bis(benzyloxy)-3-(4-bromophenyl)pyridine **(Intermediate-33)** (1.0 g, 2.24 mmol) and tert-butyl piperazine-1-carboxylate **(Intermediate-34)** (0.576 g, 2.688 mmol) in 30 mL of 1,4-dioxane were added cesium carbonate (1.825 g, 5.6 mmol) XPhos (0.107 g, 0.22 mmol) and tris(dibenzylideneacetone)dipalladium (0) (0.205 g, 0.22 mmol). The reaction mixture was degassed with argon gas for 20 minutes, then stirred at 90 °C for 6 hours under argon atmosphere. Upon completion, the reaction mixture was filtered through celite bed, the filtrate was diluted with ethyl acetate and washed with water, brine solution, dried over anhydrous sodium sulphate and concentrated in vacuo. The residue was purified by column chromatography on silica gel (10% ethyl acetate and hexane) to give the title compound **(Intermediate-35)** (0.35 g, 27 %) of tert-butyl 4-(4-(2,6-bis(benzyloxy)pyridin-3-yl)phenyl)-3,3-dimethylpiperazine-1-carboxylate. LCMS: 580.25 $[M+H]^+$.

**Step-3: Synthesis of tert-butyl 4-(4-(2,6-dioxopiperidin-3-yl)phenyl)-3,3-dimethylpiperazine-1-carboxylate (Intermediate-36)**

To a solution of tert-butyl 4-(4-(2,6-bis(benzyloxy)pyridin-3-yl)phenyl)-3,3-dimethylpiperazine-1-carboxylate **(Intermediate-35)** (0.29 g, 0.5 mmol) in 15 mL of Ethanol and 15 mL of THF was added 10% Palladium on carbon (0.26 g, 2.5 mmol). The reaction mixture was stirred at room temperature under 80psi hydrogen gas pressure for 24 hours. Upon completion, the reaction mixture was filtered through celite bed, the filtrate was concentrated under reduced pressure to give the title compound **(Intermediate-36)** (0.20 g, 99 %) of tert-butyl 4-(4-(2,6-dioxopiperidin-3-yl)phenyl)-3,3-dimethylpiperazine-1-carboxylate. $^{1}$H NMR (400 MHz, DMSO-d6): δ 10.82 (s, 1H), 7.13 – 7.06 (m, 2H), 6.92 – 6.90 (m, 2H), 3.86 – 3.78 (m, 1H), 3.44 – 3.39 (m, 2H), 3.25 – 3.23 (m, 2H), 3.09 – 3.06 (m, 2H), 2.68– 2.58 (m, 1H), 2.48– 2.44 (m, 1H), 2.16 –2.11 (m, 1H), 2.03 –1.98 (m, 1H), 1.42 (s, 9H), 1.36 (s, 6H); LCMS: 402.2 $[M+H]^+$.

**Step-4: Synthesis of 3-(4-(2,2-dimethylpiperazin-1-yl)phenyl)piperidine-2,6-dione hydrochloride (Intermediate-37)**

To a solution of tert-butyl 4-(4-(2,6-dioxopiperidin-3-yl)phenyl)-3,3-dimethylpiperazine-1-carboxylate **(Intermediate-36)** (0.20 g, 0.498 mmol) in 2 mL of 1,4-dioxane was added 4M HCl in 1,4 dioxane (0.4 mL, 20v) at 10 °C under nitrogen atmosphere. The reaction mixture was stirred for 5 hours at room temperature. Upon completion, the reaction mixture was concentrated in vacuo. The residue was washed with diethyl ether and dried under reduced pressure to give the title compound **(Intermediate-37)** (0.18 g, 95%) of 3-(4-(2,2-dimethylpiperazin-1-yl)phenyl)piperidine-2,6-dione hydrochloride. LCMS: 302.15 $[M+H]^+$.

**Scheme-9: Synthesis of Intermediate-42:**

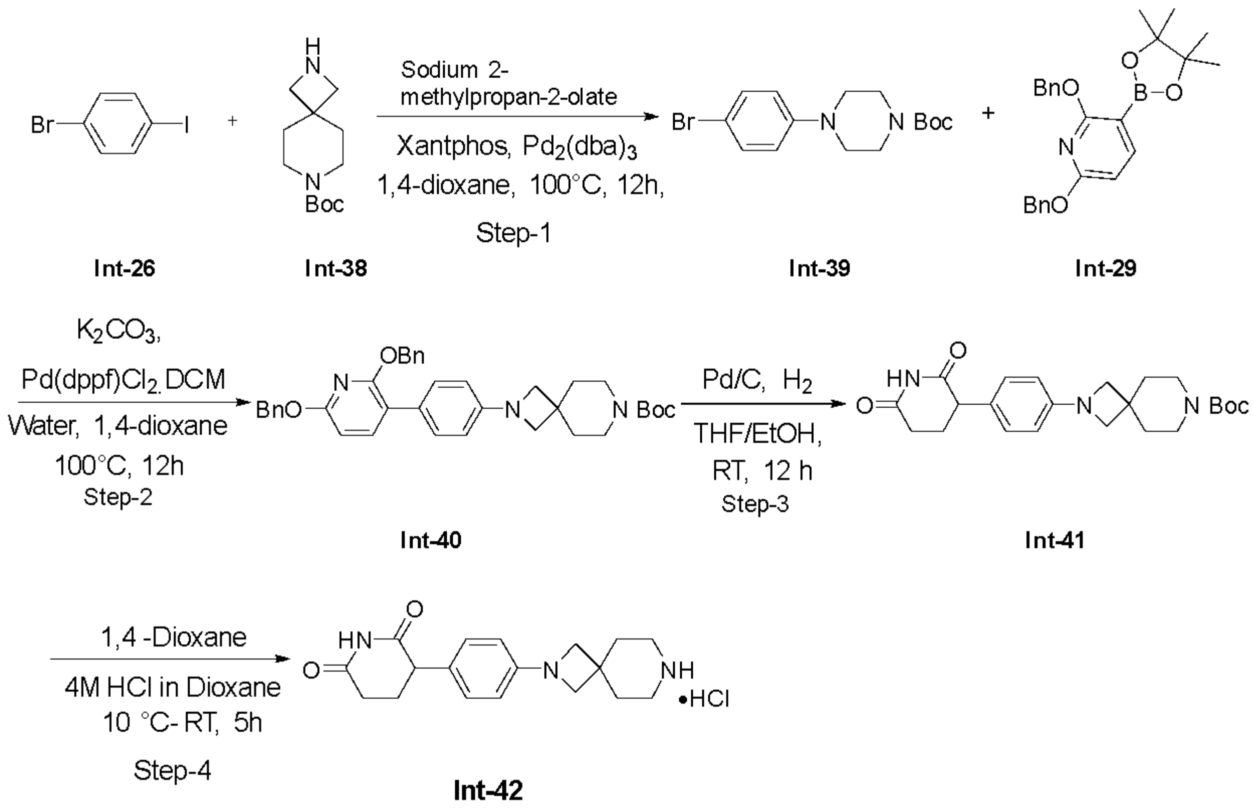


**Step-1: Synthesis of tert-butyl 2-(4-bromophenyl)-2,7-diazaspiro[3.5]nonane-7-carboxylate (Intermediate-39)**

To a solution of 1-bromo-4-iodobenzene **(Intermediate-26)** (1.0 g, 3.535 mmol) and tert-butyl 2,7-diazaspiro[3.5]nonane-7-carboxylate **(Intermediate-38)** (1.2 g, 5.302 mmol) in 10 mL of 1,4-dioxane were added Sodium 2-methylpropan-2-olate (1.01 g, 10.6 mmol), Xantphos (0.256 g, 0.44 mmol) and tris(dibenzylideneacetone)dipalladium (0) (0.162 g, 0.17 mmol). The reaction mixture was degassed with Argon gas for 20 minutes, then continued for 12 hours at 100 °C under argon atmosphere. Upon completion the reaction mixture was filtered through celite bed, the filtrate was diluted with ethyl acetate and washed with water, brine solution, dried over anhydrous sodium sulphate and concentrated in vacuo. The residue was purified by column chromatography on silica gel (50% ethyl acetate/hexane) to give the title compound **(Intermediate-39)** (1.2 g, 89 %) of tert-butyl 2-(4-bromophenyl)-2,7-diazaspiro[3.5]nonane-7-carboxylate. LCMS: 381.3 $[M]^+$.

**Step-2: Synthesis of tert-butyl 2-(4-(2,6-bis(benzyloxy)pyridin-3-yl)phenyl)-2,7-diazaspiro[3.5]nonane-7-carboxylate (Intermediate-40)**

To a solution of tert-butyl 2-(4-bromophenyl)-2,7-diazaspiro[3.5]nonane-7-carboxylate **(compound-39)** (1.1 g, 2.88 mmol) and 2,6-bis(benzyloxy)-3-(4,4,5,5-tetramethyl-1,3,2-dioxaborolan-2- yl)pyridine **(Intermediate-29)** (1.80 g, 4.32 mmol) in 16 mL of 1,4-dioxane and 4 mL of water were added potassium carbonate (1.19 g, 8.65 mmol) and Pd(dppf)$Cl_2$.DCM (0.23 g, 0.289 mmol). Then the reaction mixture was degassed with argon gas for 20 minutes. The reaction mixture was stirred at 100 °C for 12 hours under argon atmosphere. Upon completion the reaction mixture was filtered through celite bed, the filtrate was diluted with ethyl acetate and washed with water, brine solution, dried over anhydrous sodium sulphate and concentrated in vacuo. The residue was purified by column chromatography on silica gel (50% ethyl acetate and hexane) to give the title compound **(Intermediate-40)** (0.8 g, 46 %) of tert-butyl 2-(4-(2,6-bis(benzyloxy)pyridin-3-yl)phenyl)-2,7-diazaspiro[3.5]nonane-7-carboxylate. $^1$H NMR (400 MHz, DMSO-d6): δ 7.66 – 7.63 (m, 1H), 7.44 – 7.30 (m, 13H), 6.51 – 6.49 (m, 1H), 6.44 – 6.42 (m, 2H), 5.38 (s, 2H), 5.35 (s, 2H), 3.58 (bs, 4H), 3.24 – 3.22 (m, 4H), 1.70 – 1.67 (m, 4H), 1.40 (s, 9H); LCMS: 592.3 $[M+H]^+$.

**Step-3: Synthesis of tert-butyl 2-(4-(2,6-dioxopiperidin-3-yl)phenyl)-2,7-diazaspiro[3.5]nonane-7-carboxylate (Intermediate-41)**

To a solution of tert-butyl 2-(4-(2,6-bis(benzyloxy)pyridin-3-yl)phenyl)-2,7-diazaspiro[3.5]nonane-7-carboxylate **(Intermediate-40)** (0.8 g, 1.352 mmol) in 20 mL of Ethanol and 20 mL of THF was added 10% Palladium on carbon (0.72 g, 6.76 mmol). The reaction mixture was stirred at room temperature under 80 psi hydrogen gas pressure for 12 hours. Upon completion, the reaction mixture was filtered through celite bed, the filtrate was concentrated in vacuo to give the title compound **(Intermediate-41)** (0.5 g, 89%) of tert-butyl 2-(4-(2,6-dioxopiperidin-3-yl)phenyl)-2,7-diazaspiro[3.5]nonane-7-carboxylate. LCMS: 414.15 $[M-H]^+$.

**Step-4: Synthesis of 3-(4-(2,7-diazaspiro[3.5]nonan-2-yl)phenyl)piperidine-2,6-dionehydrochloride (Intermediate-42)**

To a solution of 2-(4-(2,6-dioxopiperidin-3-yl)phenyl)-2,7-diazaspiro[3.5]nonane-7-carboxylate **(Intermediate-41)** (0.5 g, 1.209 mmol) in 4 mL of 1,4-dioxane was added 4M HCl in 1,4 dioxane (6 mL, 12v) at 10 °C under nitrogen atmosphere. The reaction mixture was stirred for 5 hours at room temperature. Upon completion, the reaction mixture was concentrated in vacuo. The residue was washed with diethyl ether and dried under reduced pressure to give the title compound **(Intermediate-42)** (0.37 g, 87%) of 3-(4-(2,7-diazaspiro[3.5]nonan-2-yl)phenyl)piperidine-2,6-dione hydrochloride.LCMS: 314.2 $[M+H]^+$.

**Scheme-10: Synthesis of AU-29140 (GSPT1 compound1):**

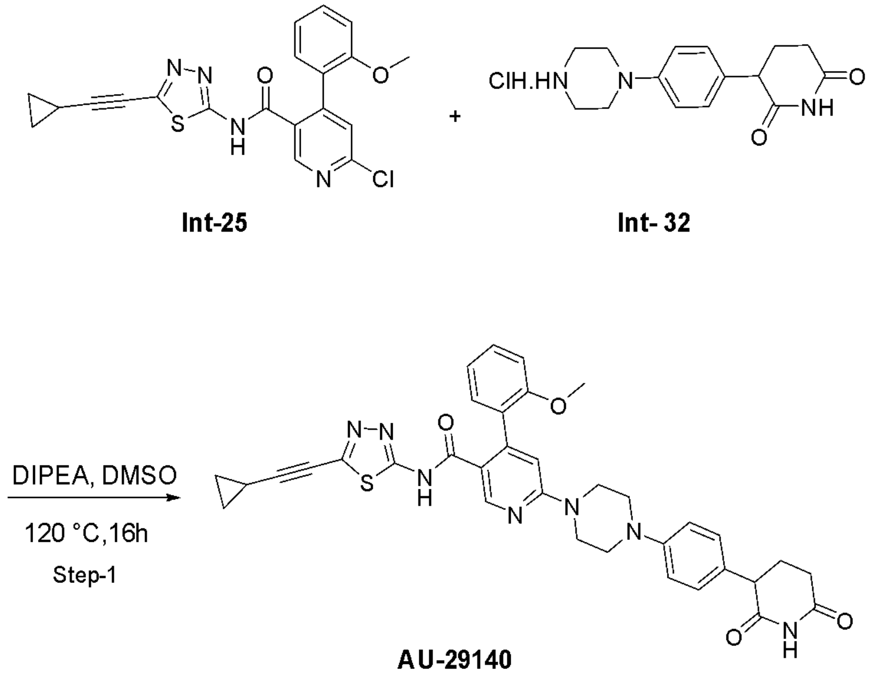


**Step-1: Synthesis of N-(5-(cyclopropylethynyl)-1,3,4-thiadiazol-2-yl)-6-(4-(4-(2,6-dioxopiperidin-3-yl)phenyl) piperazin-1-yl)-4-(2-methoxyphenyl)nicotinamide** (**AU-29140**). To a solution of

6-chloro-N-(5-(cyclopropylethynyl)-1,3,4-thiadiazol-2-yl)-4-(2-methoxyphenyl) nicotinamide (**Intermediate-25)** (0.16 g, 0.389 mmol) in 5 mL of DMSO were added 3-(4-(piperazin-1-yl)phenyl)piperidine-2,6-dione hydrochloride (**Intermediate-32)** (0.213 g, 0.778 mmol), and N,N diiso propyl ethylamine (0.25 g, 1.940 mmol) at room temperature under nitrogen atmosphere. The reaction mixture was heated to 120°C for 16 hours. Upon completion, the reaction mixture was concentrated in vacuo. The residue was purified by reverse phase column chromatography (Mobile Phase; A- 0.1 % formic acid in water, B- Acetonitrile, COLUMN: Kinetex EVO, c18 Dimension 250 x 21.2mm, 5µ) to give the title compound, **AU-29140** (0.025 g, 10 %) of N-(5-(cyclopropylethynyl)-1,3,4-thiadiazol-2-yl)-6-(4-(4-(2,6-dioxopiperidin-3-yl)phenyl)piperazin-1-yl)-4-(2-methoxy phenyl)nicotinamide. $^{1}$H NMR (400 MHz, DMSO- d6) δ 12.86 (s, 1H), 10.78 (s, 1H), 8.44 (s,1H), 7.40 – 7.34 (m, 2H), 7.10 – 7.06 (m, 3H), 7.04 – 6.98 (m, 3H), 6.73 (s,1H), 3.84 – 3.82 (m, 4H), 3.77 – 3.73 (m, 1H), 3.44 (s, 3H), 3.25 – 3.24 (m, 4H), 2.67 – 2.60 (m, 1H), 2.49 – 2.45 (m, 1H), 2.16 – 2.12 (m, 1H) ,2.04 – 1.99 (m, 1H), 1.71 – 1.65 (m, 1H), 1.01 – 0.96 (m, 2H), 0.88 – 0.84 (m, 2H). LCMS:648.20 $[M+H]^{+}$;

**Scheme-11: Synthesis of AU-30489 (GSPT1 compound 2):**

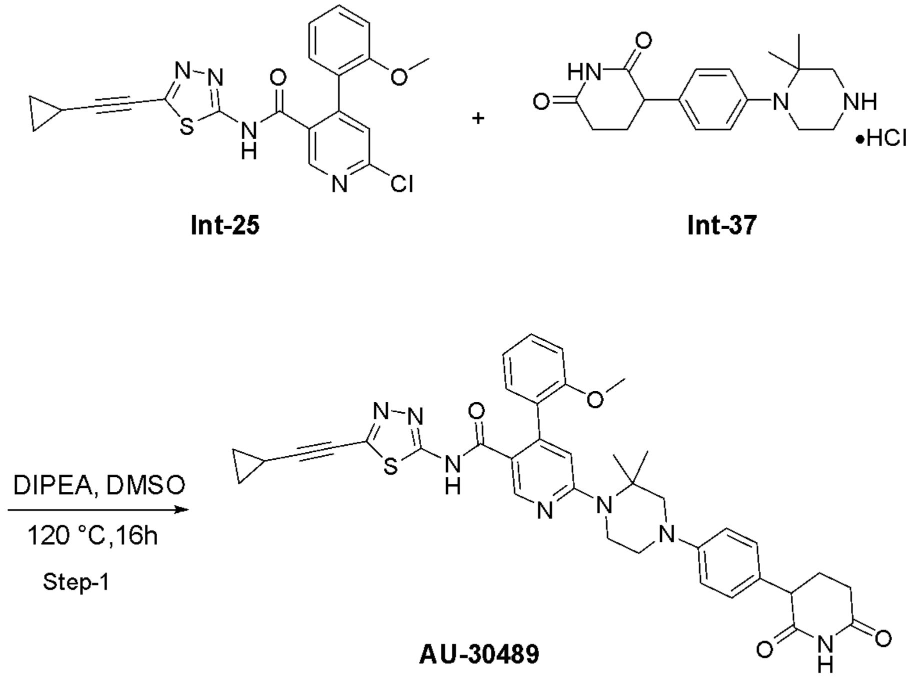

**Step-1: Synthesis of N-(5-(cyclopropylethynyl)-1,3,4-thiadiazol-2-yl)-6-(2-(4-(2,6-dioxopiperidin-3-yl)phenyl)-2,7-diazaspiro[3.5]nonan-7-yl)-4-(2-methoxyphenyl) nicotinamide** (**AU-30489**).

To a solution of 6-chloro-N-(5-(cyclopropylethynyl)-1,3,4-thiadiazol-2-yl)-4-(2-methoxyphenyl) nicotinamide (**Intermediate-25)** (0.08 g, 0.195 mmol) in 5 mL of DMSO were added 3-(4-(2,2-dimethylpiperazin-1-yl)phenyl)piperidine-2,6-dione hydrochloride (**Intermediate-37)** (0.1 g, 0.332 mmol), N,N diiso propyl ethylamine (0.15 g, 1.21 mmol) at room temperature under nitrogen atmosphere. The reaction mixture was heated to 120°C for 16 hours. Upon completion the reaction mixture was concentrated in vacuo. The residue was purified by reverse phase column chromatography (Mobile Phase; A- 0.1 % formic acid in water, B- Acetonitrile, COLUMN: Kinetex EVO, c18 Dimension 250 x 21.2mm, 5μ) to give the title compound, **AU-30489** (0.01 g, 6%) of **(AU-30489)** N-(5-(cyclopropylethynyl)-1,3,4-thiadiazol-2-yl)-6-(2-(4-(2,6-dioxo piperidin-3-yl)phenyl)-2,7-diazaspiro[3.5]nonan-7-yl)-4-(2-methoxyphenyl)nicotinamide. $^1$H NMR (400 MHz, DMSO-d6) δ 12.83 (s, 1H), 10.82 (s, 1H), 8.41 (s,1H), 7.38 – 7.34 (m, 2H), 7.15 – 7.04 (m, 5H), 6.94 – 6.92 (m, 1H), 6.79 (s,1H), 3.84 – 3.80 (m, 3H), 3.67 – 3.65 (m, 2H), 3.45 (s, 3H), 3.17 – 3.14 (m, 2H), 2.67 – 2.60 (m, 1H), 2.49 – 2.45 (m, 1H), 2.16 – 2.12 (m, 1H) ,2.09 – 2.03 (m, 1H), 1.68 – 1.67 (m, 1H), 1.24 (s, 6H), 1.05 – 0.98 (m, 2H), 0.88 – 0.84 (m, 2H). LCMS: 676.05 $[M+H]^+$;
